# Diversity of spider families parasitized by fungal pathogens: a global review

**DOI:** 10.1101/2023.05.11.540451

**Authors:** Martin Nyffeler, Nigel Hywel-Jones

## Abstract

In this paper the findings of a global literature and social media survey of spider mycoses are presented. Our survey revealed that spider mycoses occur in the geographic belt between latitude 78°N and 52°S, and that more than 40 out of the known 135 spider families (ca. 30%) are attacked by fungal pathogens. Jumping spiders (Salticidae), cellar spiders (Pholcidae), and sheet-web spiders (Linyphiidae) are the families most frequently reported to be attacked by fungal pathogens (combined >40% of all reported cases). Ninety-two percent of the infections of spiders can be attributed to pathogens in the order Hypocreales (phylum Ascomycota), and almost exclusively the families Cordycipitaceae and Ophiocordycipitaceae. Within the Hypocreales, the asexually reproductive genus *Gibellula* is an historically species-rich and widespread genus of specific spider-pathogenic fungi. For ca. 70 species of spider-pathogenic fungi their hosts could be identified at least to family level. The data presented here reaffirm the findings of previous studies that spider-pathogenic fungi are most common and widespread in tropical and subtropical forested areas, with free-living cursorial hunters – dominated by Salticidae – being the most frequently infected. Cursorial hunters (especially Salticidae) and subterranean cellar spiders (Pholcidae) are the most frequently fungus-infected spiders in North America, whereas web-weavers (especially Linyphiidae and Pholcidae) are the most common spider hosts in Europe. Our survey implies that spider-pathogenic fungi are an important mortality factor for spiders which has hitherto been underestimated.

---

Fungal pathogens are undoubtedly an important source of mortality for a large number of arthropod groups. Insects alone are attacked by >700 species of entomopathogenic fungi (Hajek & St. Leger 1994; Consolo et al. 2003) and accordingly, these fungi have been extensively studied. By way of contrast, spider-pathogenic fungi have attracted less scientific interest. They have been almost neglected as spider enemies by arachnologists. This is evident from the fact that in most books on spider biology, fungal pathogens were not mentioned in chapters dealing with the spiders’ natural enemies (see Gertsch 1979; Nentwig 1987a; Wise 1993; Dippenaar-Schoeman & Jocqué 1997; Foelix 2011; Bradley 2013).

Although a larger number of papers on the taxonomy of spider-pathogenic fungi exist, most of these studies are of limited value for the arachnologist because the authors of such studies invariably referred to the spider hosts as “unknown spider”. The lack of knowledge of the identity of spider hosts can be explained: 1) by the fact that almost all of these past studies on spider-pathogenic fungi were conducted by mycologists with rather limited interest or expert knowledge regarding the identification of spiders; 2) due to the lack of local experts to correctly identify the host and the difficulties of transferring materials (following Nagoya Protocol and Convention on Biological Diversity) to countries where experts reside oftentimes encountering ‘red tape’; and 3) by the extreme difficulty in reliably identifying parasitized spiders whose identifying features are often completely overgrown by the pathogen itself (Evans 2013; Durkin et al. 2021; Kozlov 2021; Mendes-Pereira et al. 2022).

Evans (2013: p. 107) stated with regards to spider-pathogenic fungi “*This has been a somewhat neglected topic: typically, a no-man’s land between mycologists and arachnologists*”. In the current review, we aim to close the gap between mycology and arachnology – thereby pursuing a mycological-arachnological interdisciplinary approach by looking at spider-pathogenic fungi from an arachnological point of view (Fig. 1). It is hoped that this review will generate interest among arachnologists in the spider-pathogenic fungi and that it will inspire them to start exploring the still unresolved problems on this topic in the future. Aside from that, we hope that it will also benefit mycologists by broadening their horizons in terms of spider host-fungal pathogen relationships. Historically, the mycological literature has lumped all arachnids in with Insecta when dealing with invertebrate-pathogenic fungi. Where there have been distinctions, this has been between mites and ‘spiders’. In the mycological literature ‘spiders’ has also included the harvestmen. In this review, we do not treat the mites or harvestmen and consider only the families within the order Araneae.

**Figure 1.**
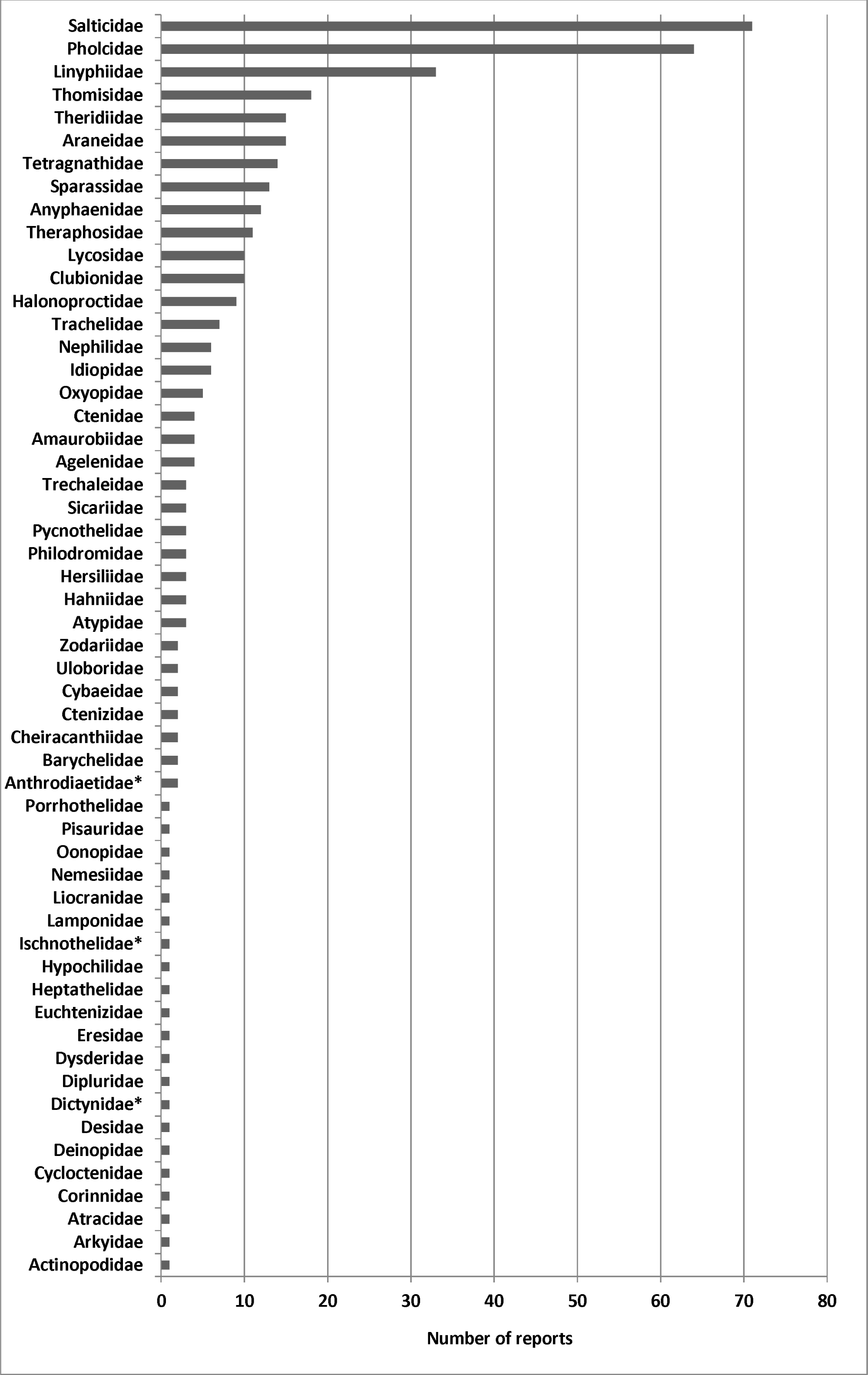
Fifty-five spider families containing species parasitized by fungi (see Supplementary Table S1 for details). * Fungus infections observed exclusively under laboratory conditions. Extinct members of the families Gnaphosidae?, Mysmenidae, and Synotaxidae with fungi growing on their bodies – reported as amber inclusions (see Wunderlich 2004) – are not included here.

## METHODS

### Data collection

We searched published reports on fungal pathogens in the Web of Science, Scopus, Google Search, Google Scholar, Google Books, Google Pictures, ProQuest Dissertations and Theses, as well as USDA-ARS Collection of Entomopathogenic Fungal Cultures – Catalog of Strains (Humber et al. 2014) following the same search method as Nyffeler & Gibbons (2022). Additionally, we made a library search of books and scientific journals not included in the electronic databases. Social media sites (e.g., BugGuide, What’s That Bug, iNaturalist, Facebook, Twitter, Flickr, Getty Images, Reddit, Yahoo, and YouTube) were also searched for information on spider-pathogenic fungi. Our survey resulted in a total of 511 records of entomogenous fungi growing on spiders, 92% of which could be identified, while 8% remained unidentified (see Supplementary Table S1). [“Entomogenous” = fungi growing on or in the bodies of arthropods, including pathogens, facultative pathogens, or fungi whose trophic status is ambiguous.] In 394 cases (roughly 77%) the spider host could be determined at least to family level, whereas in 117 cases (23%) the spider hosts remained unidentified (see Supplementary Table S1).

With regard to the cases with identified spider hosts, information on the place of discovery of the fungus-infected spider was available in 355 cases (see Supplementary Table S1), and these data were used for the statistical comparison of the dominant fungus-infected spider families in different major geographic regions (see Fig. 7). Records referring to the Russian Federation (see Humber et al. 2014) and the arctic island of Jan Mayen (see Bristowe 1948) were listed under Europe in all cases.

### Taxonomic comments and data sources

We have the following comments regarding data sources and the taxonomy of fungi and spiders, respectively.

#### Asexual morph / sexual morph concept in mycology

Fungi in the phyla Ascomycota and Basidiomycota usually have two different reproductive stages: an asexual stage and a sexual stage (Hodge 2003). The asexual and sexual reproductive stages of a particular fungus were historically recognized as distinct species, although they may be genetically identical. Sometimes only one of the two reproductive stages of a fungal pathogen is known. With the advent of molecular phylogenetics the links between asexual and sexual species is now becoming clear. As a result, the last 10+ years has seen a move towards the One Fungus One Name concept (Taylor 2011; Rossman 2014; Crous et al. 2015). This is a dynamic concept that is beyond the scope of this review to fully detail.

Information on the reproductive mode of the fungal species associated with spiders (see Appendix 1, third column) was extracted from various literature sources:

– Samson & Evans 1973; Sung et al. 2001; Swart et al. 2001; Benny 2008; Rivera 2009; Evans 2013; James et al. 2016; Kepler et al. 2017; Walther et al. 2019; St. Leger & Wang 2020; Xu et al. 2021; Yu et al. 2021; Chen et al. 2022a; Mongkolsamrit et al. 2022; Wang et al. 2023a; https://fungalgenera.org/genus/acrodontium.html: *Acrodontium*, *Akanthomyces*, *Aphanocladium*, *Apiotrichum*, *Aspergillus*, *Basidiobolus*, *Beauveria*, *Cladosporium*, *Clathroconium*, *Clonostachys*, *Conidiobolus*, *Engyodontium*, *Gibellula*, *Hevansia*, *Hirsutella*, *Hymenostilbe*, *Jenniferia* (i.e., *Jenniferia cinerea* – other *Jenniferia* spp. classified differently, see below), *Lecanicillium*, *Metarhizium*, *Mucor*, *Neoaraneomyces*, *Parahevansia*, *Parengyodontium*, *Penicillium*, *Pseudogibellula*, *Pseudometarhizium*, *Purpureocillium* [= *Nomuraea*], *Sporodiniella*, *Tolypocladium*, and unknown Hyphomycetes were found with asexual morphs (A).

– Evans 2013; Mongkolsamrit et al. 2022: *Cordyceps*, *Ophiocordyceps*, *Polystromomyces*, and *Torrubiella* represent the sexual reproductive stage (S).
– Mongkolsamrit et al. 2022, 2023: *Bhushaniella rubra*, *Jenniferia griseocinerea* and *J. thomisidarum* represent species which were found with asexual and sexual morphs (A, S).
– Malloch et al. 1978: *Cryptococcus depauperatus* (sexual morph named *Filobasidiella arachnophila* was found on a dead spider) (S).
– Evans & Samson 1982, Evans 2013: *Cordyceps farinosa* (asexual morph *Isaria farinosa* [= *Paecilomyces farinosus*] was found on dead spiders) (A).

– Evans 2013; Thúy et al. 2015: *Cordyceps javanica* (asexual morph [= *Isaria javanica*] was found on dead spiders) (A).

*Status of fungi as pathogens, facultative pathogens, saprobes or other types of fungus/spider associations*: An assignment of the fungal species associated with spiders to a particular infection type (see Appendix 1, second column) was based on the following literature sources:

– Evans & Samson 1987; Bibbs et al. 2013; Evans 2013; Manfrino et al. 2017; Shrestha et al. 2019; Chen et al. 2022a; Wang et al. 2023a; Mongkolsamrit et al. 2023: *Bhushaniella rubra*, *Clathroconium arachnicola*, *Clathroconium* sp., *Clonostachys aranearum*, *Metarhizium anisopliae*, *Mucor fragilis*, *Neoaraneomyces araneicola*, *Pseudometarhizium araneogenum*, *Purpureocillium lilacinum,* all Cordycipitaceae, all Ophiocordycipitaceae, all unidentified Hypocreales considered as pathogens (P).
– Noordam et al. 1998; Koukol 2010; Evans 2013; Chang et al. 2022: *Acrodontium crateriforme*, *Conidiobolus* sp., *Mucor hiemalis*, *Mucor* sp., *Sporodiniella umbellata*, and some unidentifiable Hyphomycetes all characterized as facultative pathogens (FP).
– Tan et al. 2022: *Penicillium tealii* characterized as hyperparasite of another fungus species (i.e., *Gibellula* sp.) (H).
– Greif & Currah 2007; Yoder et al. 2009; Koukol 2010; Nováková et al. 2018a,b; https://fungalgenera.org/genus/acrodontium.html: *Acrodontium crateriforme*, *Aspergillus* spp., *Cladosporium* spp., *Conidiobolus coronatus*, *Mucor* spp., *Penicillium* spp. (with the exception of *P. tealii*) characterized as saprobes (SA) which does not exclude that some of them are also listed as facultative pathogens (see above).
– Malloch et al. 1978; Henriksen et al. 2018: *Basidiobolus* sp., *Cryptococcus depauperatus* [=*Filobasidiella arachnophila*] characterized as non-pathogenic (NP).
– Heneberg & Rezac 2013; Biali et al. 1972/Humber et al. 2014: *Aphanocladium album*, *Apiotrichum* spp. characterized as unknown type of fungus/spider association (U).

#### Identification of unidentified fungi

Some photographs depicting unidentified fungi were identified to the lowest taxon possible. In the case of subterranean fungus-infected pholcids from Europe and North America, it must be said that they all looked very much alike (Fig. 2A–E). Mycosed pholcids are conspicuous by white, fluffy spheres surrounding the spider body and each of the leg joints (Eiseman et al. 2010; Martynenko et al. 2012). The resemblance in appearance of the mycosed pholcids (Fig. 2A–E) indicates that they were all attacked by a group of related fungal pathogens. This is confirmed by the observation that fungi from only one family (i.e., Cordycipitaceae) were isolated from dead subterranean pholcids (see Cokendolpher 1993; Keller 2007; Eiseman et al. 2010; Martynenko et al. 2012; Jent 2013; Humber et al. 2014; Kathie T. Hodge, pers. comm.) which led us to assume that they are exclusively attacked by fungi from this family – at least in Europe and North America. Consequently, fungi on subterranean dead pholcids were consistently assigned by us to the family Cordycipitaceae, as far as fungi from Europe and North America were concerned.

**Figure 2.**
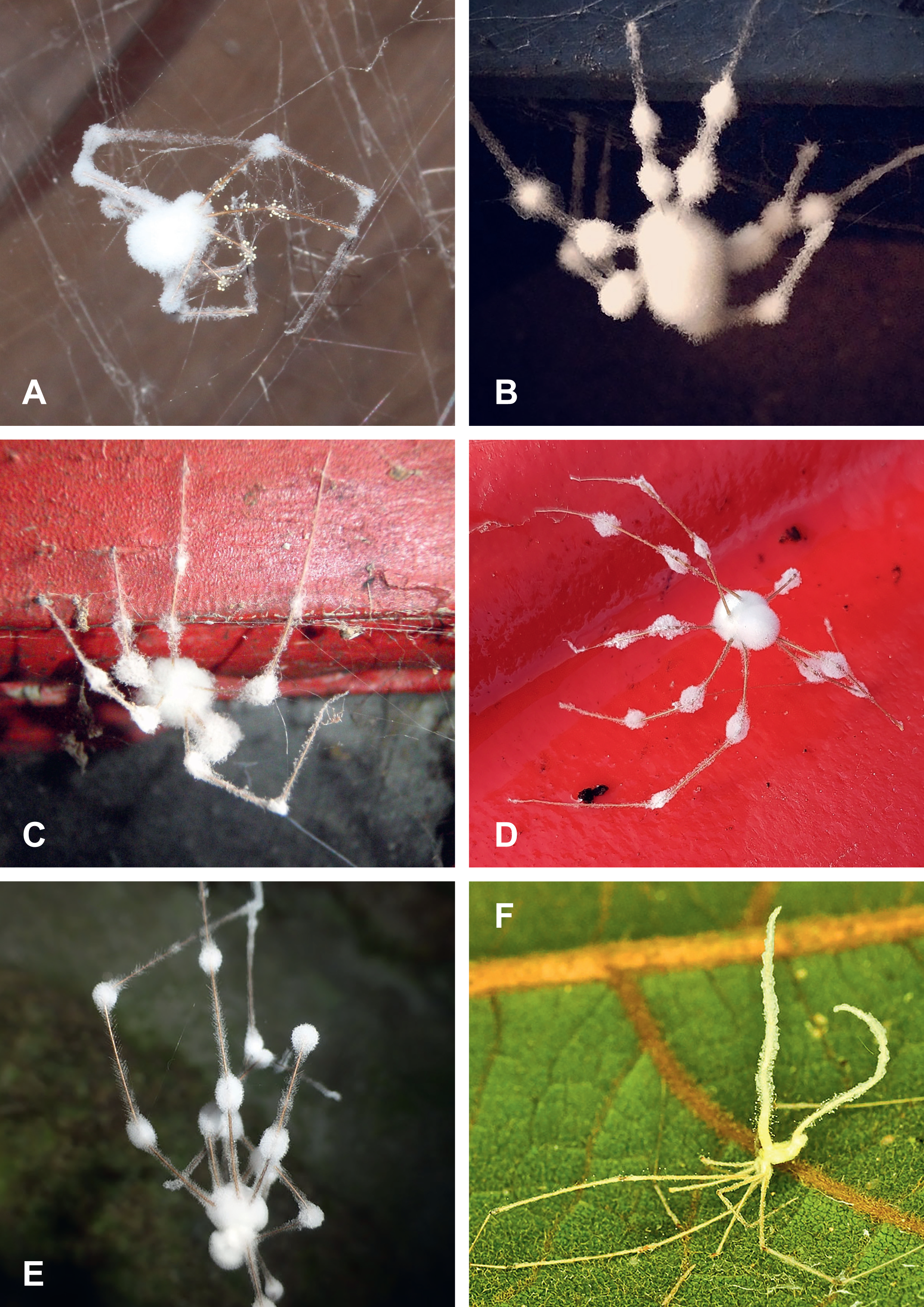
**A.–G.** Cellar spiders (*Pholcus* spp.) infected by hypocrealean fungi (Cordycipitaceae). **A.** Observed in Cottage Grove, Wisconsin, USA (Photo by Kim Wood). **B.** In basement of an old house in Pennsylvania, USA (Photo by Lynn Lunger). **C.** Observed in Wayne, New Jersey, USA (Photo by Jeffrey J. Cook). **D.** Picture taken in Uxbridge, Ontario, Canada (Photo by J. Mass / “iNaturalist” CC BY-NC 4.0). **E.** On a water motor house in Gemunde, Portugal (Photo by Carlos M. Silva). **F.** Pholcid spider *Metagonia taruma* Huber, 2000 parasitized by fungus *Gibellula* sp. on leaf underside – Floresta Nacional de Tapajós, Brazil (Photo by Leonardo Sousa Carvalho).

#### Identification of spiders

Some photographs depicting unidentified spiders were identified to the lowest taxon possible on our behalf by the following spider taxonomists: Anyphaenidae (R. Bennett, A. Dippenaar, C. Haddad, M. Ramírez, D. Ubick, A. Zamani), Cheiracanthiidae (R. Raven), Ctenidae (H. Höfer), Cybaeidae (R. Bennett), Cycloctenidae (M. Ramírez, R. Raven), Linyphiidae (T. Bauer, J. Wunderlich), Philodromidae (B. Cutler), Pholcidae (L. Sousa Carvalho), Salticidae (B. Cutler, G.B. Edwards, D. Hill), Tetragnathidae (T. Bauer, M. Kuntner, G. Oxford, A. Zamani), Theraphosidae (R. West), Thomisidae (B. Cutler), Trachelidae (T. Bauer, C. Haddad, Y. Marusik, M. Ramírez, A. Zamani), Uloboridae (A. Brescovit), and further taxonomic advice (D. Logunov).

#### Further taxonomic comments

Nomenclature of spider taxa was based on the World Spider Catalog (2023). *Argyroneta aquatica* (Clerck, 1757) previously placed in the family Cybaeidae is now placed in Dictynidae (World Spider Catalog 2023).

Several authors reported the Corinnidae among the spider families known to be infected by fungal pathogens (Costa 2014; Shrestha et al. 2019; Arruda 2020; Kuephadungphan et al. 2022). All these reports can be traced back to Costa (2014), who mentioned a fungus-infected spider identified as *Trachelas* cf. *robustus* (Chickering, 1937), a species originally placed in the family Corinnidae (see Platnick 2013). According to the World Spider Catalog (2023), the genus *Trachelas* now belongs to the family Trachelidae which implies that the fungus-infected spider in Costa’s (2014) study had a trachelid host. However, a corinnid spider parasitized by a fungus from the genus *Gibellula* has most recently been discovered in the Brazilian Atlantic forest (Mendes-Pereira et al. 2022). Furthermore, a dead corinnid spider with fungus hyphae growing on its body was found encased in a Baltic amber sample (Wunderlich 2004).

A second taxonomic issue which needs to be addressed refers to the spider family Ctenizidae. Originally, the Ctenizidae was a large family made up of 135 species placed in 9 genera all of which construct burrows in the ground sealed by a trapdoor made of soil, saliva, and silk (i.e., trapdoor spiders; see Gertsch 1979; World Spider Catalog 2018 / Version 18.5). Over many years starting with Raven (1985), many species previously placed in the Ctenizidae were transferred to other families. Accordingly, the family Ctenizidae was reduced from 135 species (in 9 genera) to 53 species (in 2 genera) (see World Spider Catalog 2018).

Today, the revised family Ctenizidae is made up of only 5 species placed in the genera *Cteniza* Latreille, 1829 and *Cyrtocarenum* Ausserer, 1871, all of which occur in the European Mediterranean region (see World Spider Catalog 2023). In the mycology literature we find many reports from Asia and South America of fungus-infected “trapdoor spiders” which were referred to as Ctenizidae (see Yakushiji & Kumazawa 1930; Petch 1937, 1939; Kobayasi 1941; Mains 1954; Haupt 2002; Evans 2013; Hughes et al. 2016; Shrestha et al. 2019). These statements reflected the state of taxonomy of that time. However, according to today’s concept of spider taxonomy, these species are no longer ctenizids, but belong to other families instead (e.g., the Asian trapdoor spider genus *Latouchia* Pocock, 1901 has been transferred to the family Halonoproctidae; see World Spider Catalog 2023). As regards the species of mygalomorph spiders from South America, many of those may be Pycnothelidae now (Martín Ramírez, pers. comm.; see also Montes de Oca et al. 2022). Nevertheless, white/light spots or patches have been observed on adult spiders in the genera *Cteniza* and *Cyrtocarenum* indicating that fungus-infections can occur in spiders of the family Ctenizidae in Europe (Arthur Decae, pers. comm.).

A further taxonomic issue which needs to be addressed refers to the spider *Stenoterommata platensis* Holmberg, 1881 which was cited by Shrestha et al. (2019) as belonging to the family Nemesiidae. Indeed this species used to be placed in the family Nemesiidae at that time (World Spider Catalog 2019). But today it is placed in the family Pycnothelidae (Opatova et al. 2020). Nevertheless, during our literature survey we found another record for a spider from the family Nemesiidae that apparently had been infected by a fungus (see Isaia & Decae 2012).

#### Predation strategies of the documented spider families

Information on the predation strategies of the documented spider families (Appendix 2) was taken from Nentwig (1987b). Additionally, information for Cycloctenidae is based on Kelly et al. (2023) and for Arkyidae on Kulkarni et al. (2023).

### Statistical methods

To test whether the percentages of fungus-infected spiders of a particular spider group differed statistically significantly between two continents, a chi-square test (without Yates’s correction) was performed using MedCalc statistical software (https://www.medcalc.org/calc/comparison_of_proportions.php).

## RESULTS

### Which fungi are engaged in the parasitization of spiders?

The entomogenous fungal taxa associated with spider hosts tracked down during our survey are listed in Tables 1–2 and in Appendix 1–2. Appendix 1 includes roughly 70 different pathogenic fungal taxa with known spider host identity; roughly 20 other types of fungal species (facultative pathogens, saprobes, non-pathogenic fungi, or hyperparasites) with known spider host are also included in this list. Furthermore, for the sake of completeness a larger number of described pathogenic fungal taxa with unidentified spider hosts have been included in the list as well. Based on the >400 records of identified fungi examined in this study (Table 1), it can be concluded that >90% of them clearly classify as spider pathogens, while the remaining <10% are either facultative pathogens, non-pathogens, or fungi whose trophic status is ambiguous. The asexual reproductive stage prevails, comprising roughly two-thirds of all identified fungal species (Appendix 1, third column).

**Table 1.**
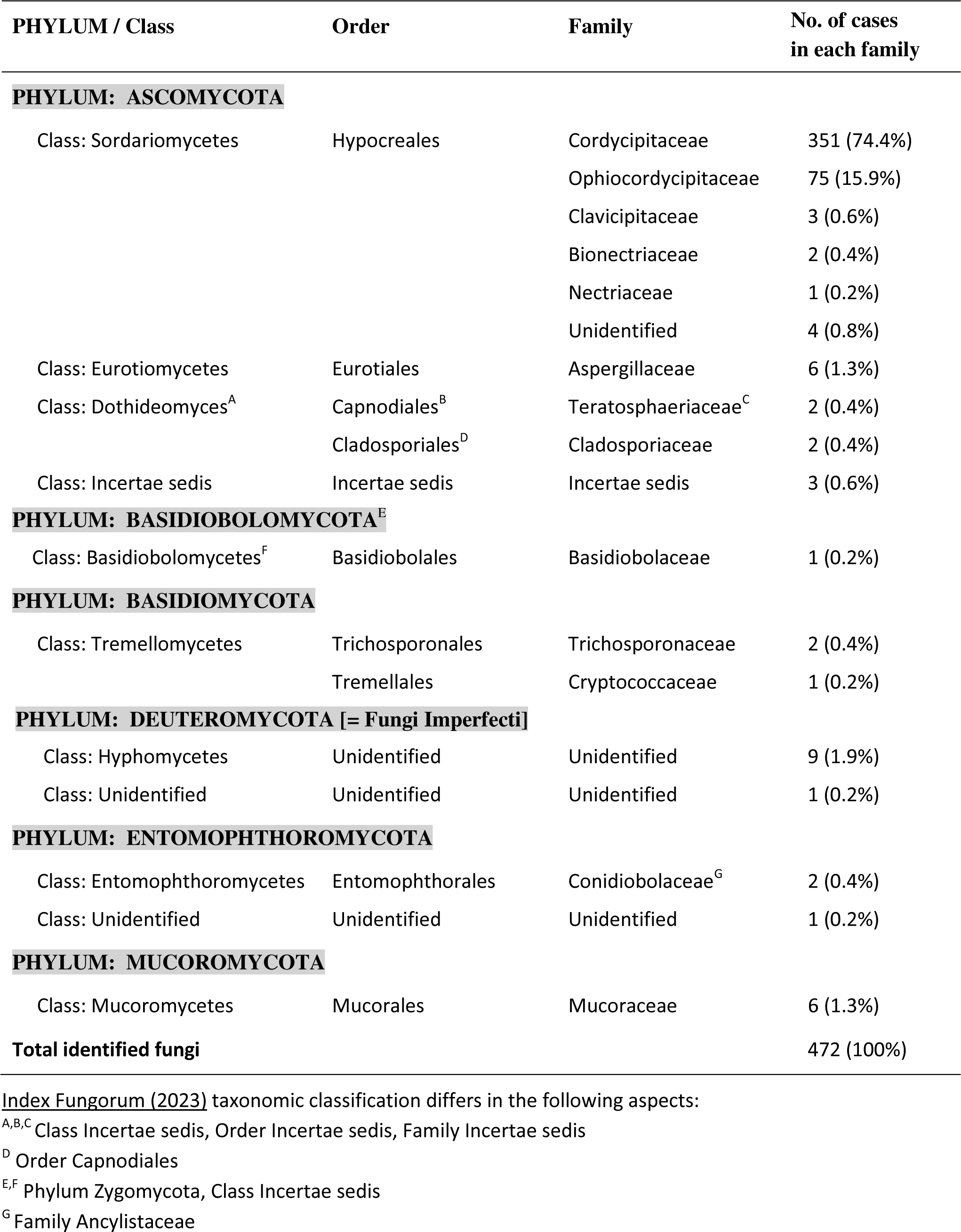
Taxonomical classification of spider-pathogenic fungi and other spider-associated fungi based on MycoBank (2023). Numbers in the last column of the table is based on Supplementary Table S1. Thirty-nine records in which the fungi could not be identified were not included in this compilation.

**Table 2.**
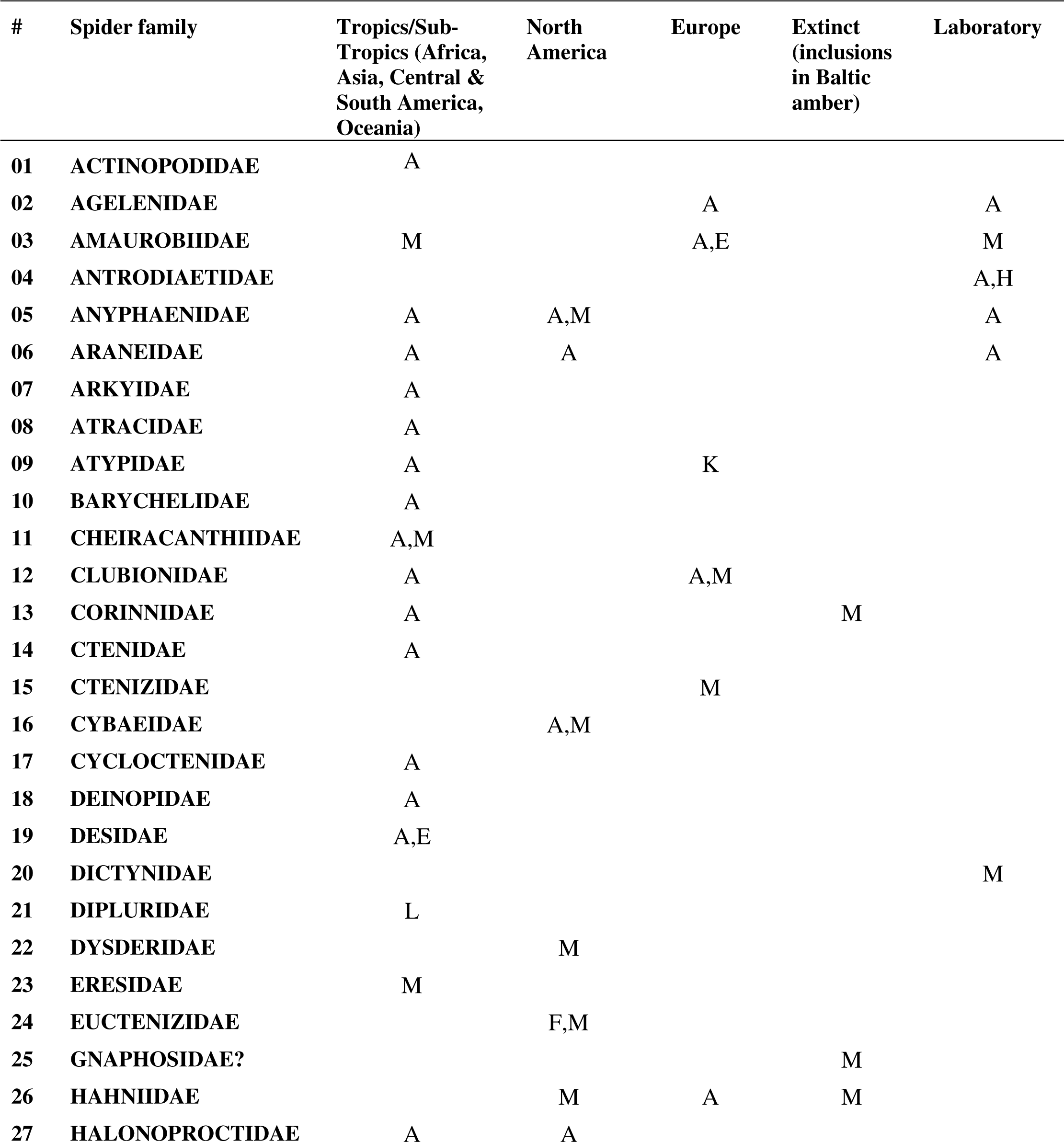
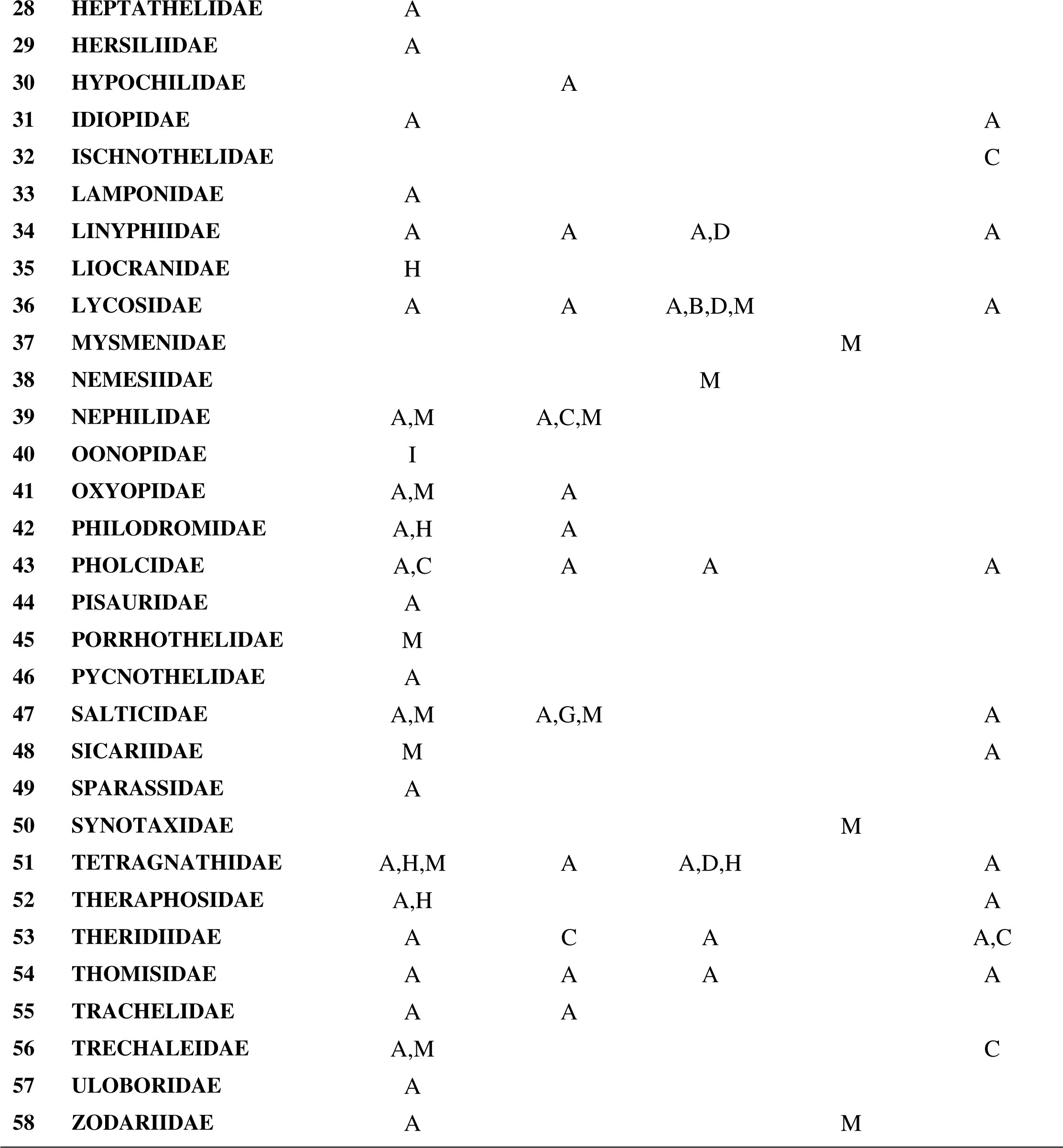
Diversity of spider families parasitized by fungal pathogens (based on Supplementary Table S1). <u>Types of fungi</u>: A = Pathogen in the order Hypocreales; B = Facultative pathogen in the order Entomophthorales; C = Facultative pathogen in the order Mucorales; D = Facultative pathogen in the class Hyphomycetes/order unknown; E = Hyperparasite overgrowing a hypocrealean pathogen; F = Hyperparasite overgrowing an unknown fungus; G = Order Capnodiales (saprobe or pathogen?); H = Saprobe in the order Eurotiales; I = Non-pathogenic in the order Basidiobolales; K = Order Trichosporonales (unknown type of infection); L = Phylum Deuteromycota (unknown type of infection); M = Unknown fungus.

Spider-pathogenic fungi belong for the most part to the phylum Ascomycota, with the order Hypocreales prevailing (Table 1; Figs. 2–6). The Hypocreales was represented by the families Bionectriaceae, Clavicipitaceae, Cordycipitaceae, Nectriaceae, and Ophiocordycipitaceae (Table 1). Hypocreales are classified as pathogenic with the exception of one spider-associated species in the family Nectriaceae (i.e., *Aphanocladium album*; see Humber et al. 2014) whose status is ambiguous (Koç & Défago 1983; Patil et al. 1994).

**Figure 3.**
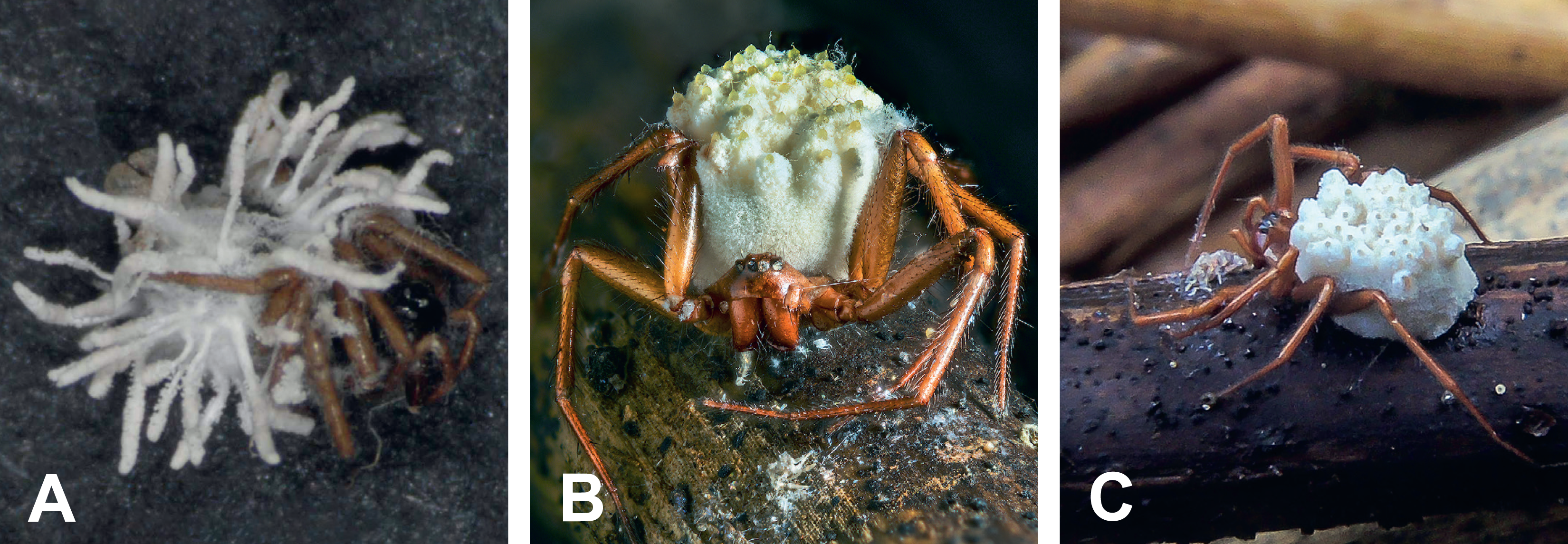
**A.** Linyphiid spider parasitized by *Gibellula pulchra* – vegetable plot in Denmark (Photo by Nicolai V. Meyling, University of Copenhagen). **B.** Linyphiid spider parasitized by *Torrubiella albolanata* – Denmark (Photo by Jens H. Petersen, “MycoKey”). **C.** Linyphiid spider parasitized by *Torrubiella albolanata* – Denmark (Photo by Thomas Læssøe, University of Copenhagen).

**Figure 4.**
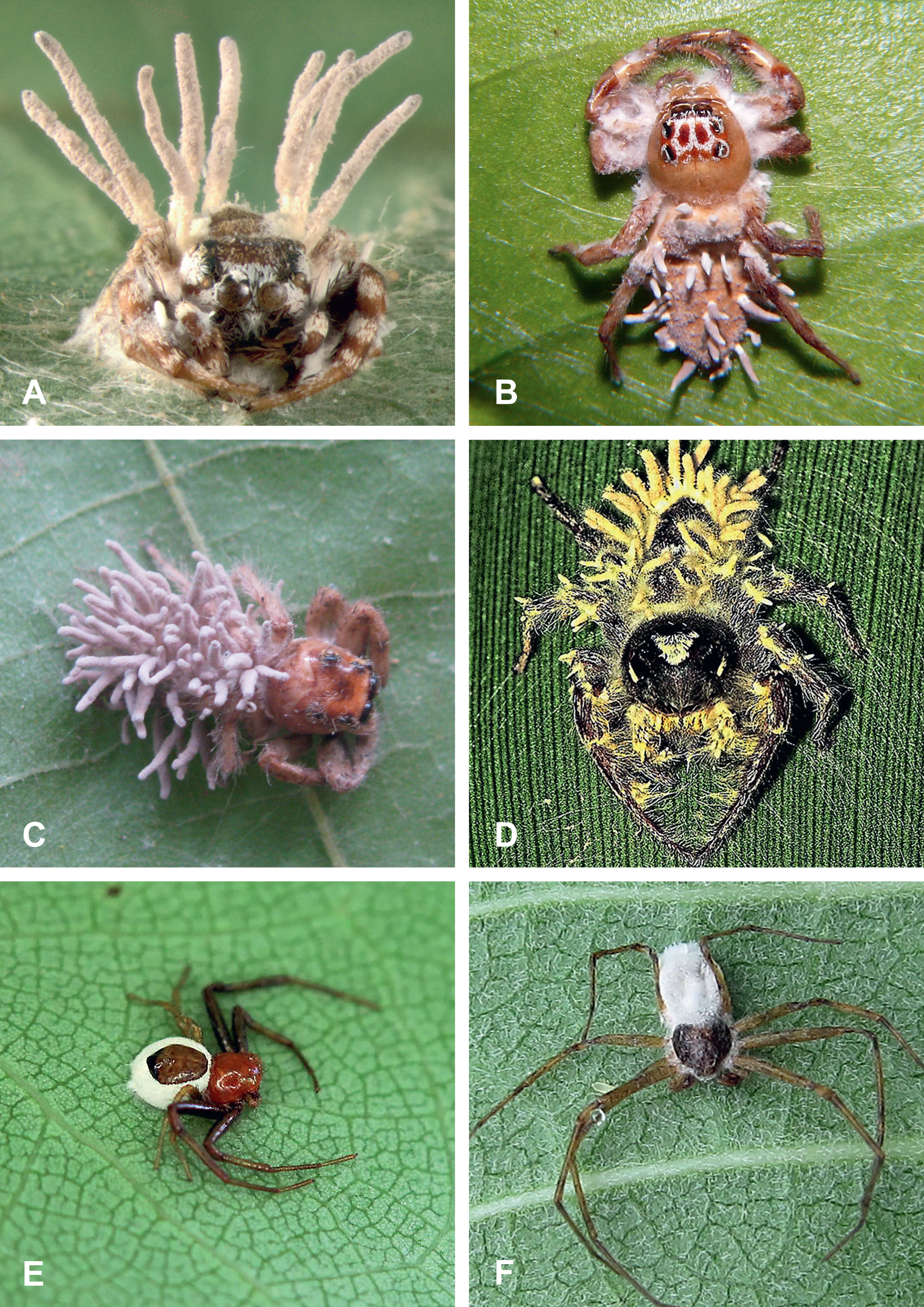
**A.** *Pelegrina proterva* (Walckenaer, 1837) (Salticidae) parasitized by *Gibellula* cf. *leiopus* – Jefferson County Park, Iowa, USA (Photo by Mary Jane Hatfield). **B.** *Mopsus mormon* Karsch, 1878 (Salticidae) parasitized by *Gibellula* sp. (a member of the *Gibellula leiopus* complex) – Airlie Beach, Queensland, Australia (Photo by Steve & Alison Pearson). **C.** Spider in the genus *Colonus* F. O. Pickard-Cambridge, 1901 (Salticidae) parasitized by *Gibellula* cf. *leiopus* – Athens/ Sandy Creek Nature Center, Georgia, USA (Photo by Carmen Champagne). **D.** *Phidippus putnami* (G. W. Peckham & E. G. Peckham, 1883) (Salticidae) parasitized by an immature *Gibellula* sp. – Bon Aqua, Tennessee, USA (Photo by Lisa Powers, “Froghaven Farm”). **E.** *Synema parvulum* (Hentz, 1847) (Thomisidae) parasitized by an immature hypocrealean fungus (most likely *Purpureocillium atypicola*) – Eno River State Park, North Carolina, USA (Photo by Tony DeSantis). **F.** Spider in the genus *Philodromus* Walckenaer, 1826 (Philodromidae) parasitized by an immature hypocrealean fungus (most likely *Purpureocillium atypicola*) –Kakiat County Park, New York, USA (Photo by Seth Ausubel).

**Figure 5.**
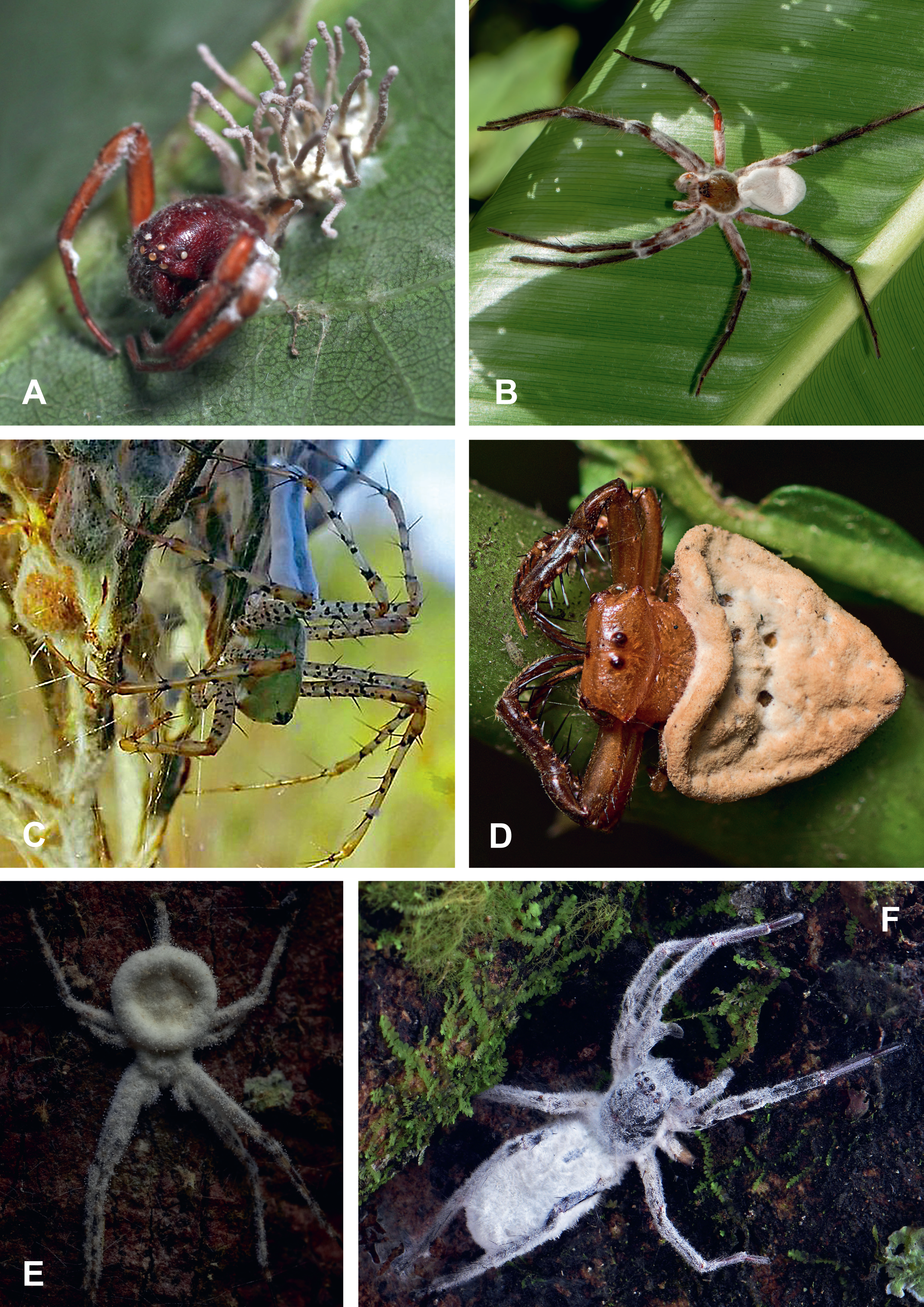
**A.** Spider in the genus *Trachelas* L. Koch, 1872 (Trachelidae) parasitized by *Gibellula leiopus* – near Donalds, South Carolina, USA (photo by Kim Fleming). **B.** Pisaurid spider parasitized by an immature hypocrealean fungus (most likely *Purpureocillium atypicola*) – Mindo, Ecuador (Photo by Gilles Arbour / “NatureWeb”). **C.** *Peucetia viridans* (Hentz, 1832) (Oxyopidae) parasitized by an immature hypocrealean fungus (most likely *Purpureocillium atypicola*) – Okeechobee County, Florida, USA (Photo by Jeff Hollenbeck). **D.** *Arkys lancearius* Walckenaer, 1837 (Arkyidae) parasitized by an immature hypocrealean fungus (most likely *Gibellula* sp.) – Brisbane, Queensland, Australia (Photo by Tony Eales). **E.** Spider in the family Hersiliidae parasitized by a *Gibellula* sp. – Tambopata Research Centre, Peruvian Amazon (Photo by Paul Bertner). **F.** Spider in the family Uloboridae parasitized by a hypocrealean fungus (Cordycipitaceae) – Mocoa, Columbia (Photo by Daniel Winkler).

**Figure 6.**
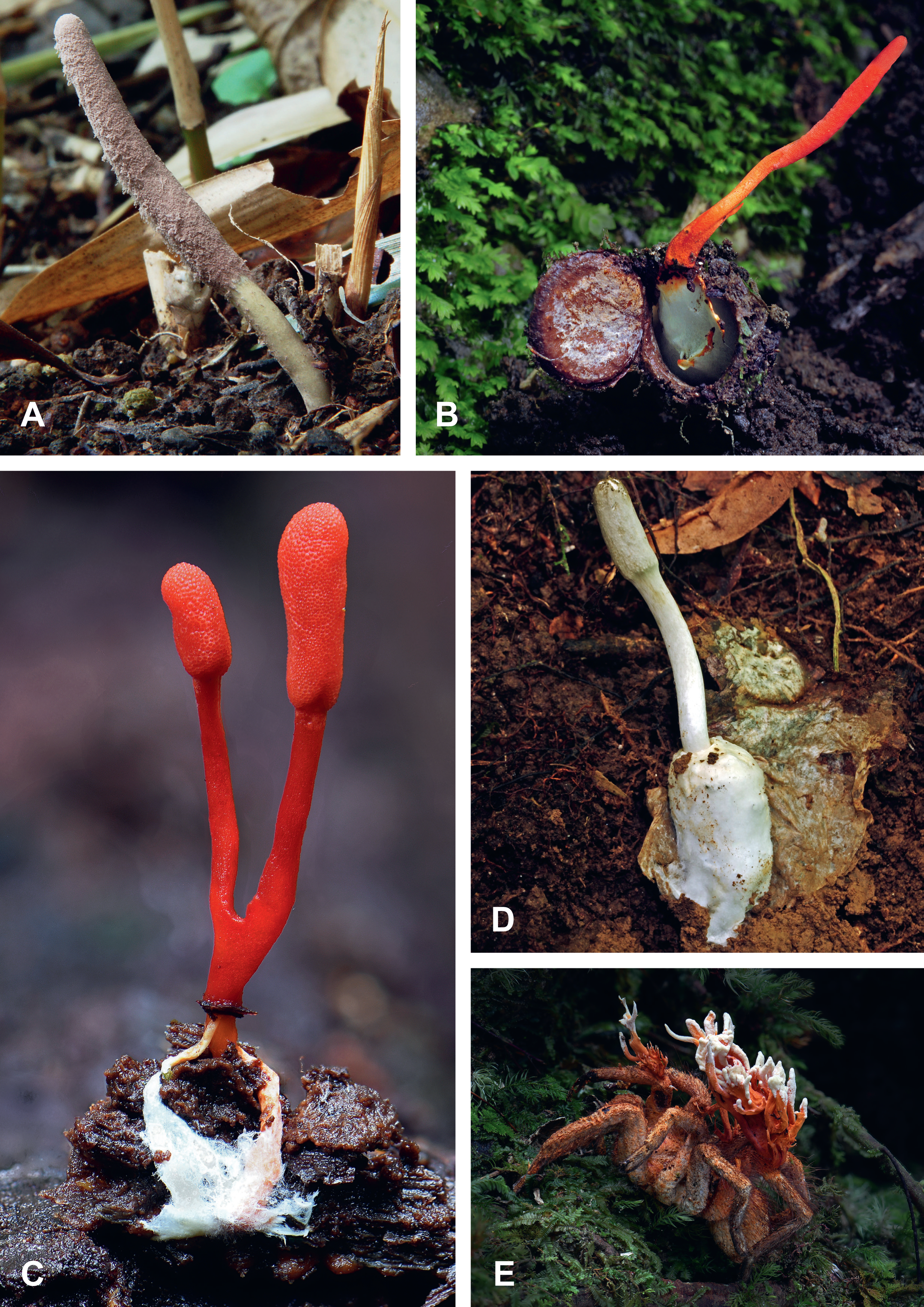
Mygalomorph spiders infected by fungi. **A.** Fruit body of *Purpureocillium atypicola* emerging from a unidentified mygalomorph spider carcass at Hongo, Bunkyo-ku, Tokyo, Japan (Photo by Sui-setz / “Wikipedia” CC BY-SA 3.0). **B.** Red fruit bodies (stromata) of *Cordyceps nidus* emerging from the underground burrow a dead trapdoor spider (family Idiopidae) – Chicaque Natural Park near Bogota, Colombia; the lid of the trapdoor has been pushed open by the emerging fruit bodies (Photo copyright: Daniel Winkler / “MushRoaming”, Seattle, USA). **C.** Red fruit bodies (stromata) of *Cordyceps nidus* emerging from a dead unidentified trapdoor spider – Isla Escondida, Colombia (Photo copyright: Daniel Winkler). **D.** Fruit body of *Cordyceps cylindrica* emerging from an dead unidentified trapdoor spider – Chalalan, Bolivia (Photo copyright: Daniel Winkler). **E.** *Cordyceps caloceroides* growing on a dead theraphosid (subfamily Theraphosinae) – Pitalito, Colombia (Photo copyright: Daniel Winkler).

**Figure 7.**
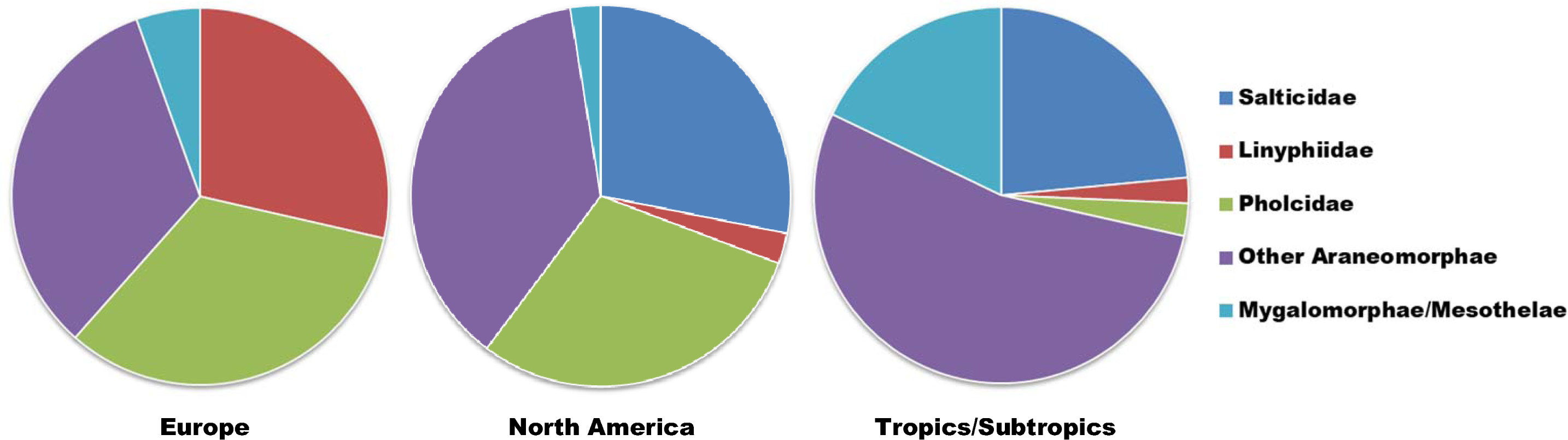
Percent composition of dominant fungus-infected spider groups in three different geographic regions: Europe (including the Russian Federation and the arctic island of Jan Mayen), North America, and Tropics/Subtropics (combined data for Africa, Asia, Central & South America, and Oceania). Pie charts generated based on Supplementary Table S1.

Based on our survey, Cordycipitaceae was the most prominent spider-pathogenic fungal family, accounting for ca. three-quarters of all recorded spider infections (Table 1). Within the Cordycipitaceae, the genus *Gibellula* (including its sexual morph *Torrubiella*) is a particularly species-rich and widespread group of specific spider-pathogenic fungi (see Chen et al. 2021; Mendes-Pereira et al. 2023). Up to the present, 50–60 *Gibellula* species have been listed in the global fungal databases “Index Fungorum” and “MycoBank”, some of which, however, have been declared to be synonyms (see Shrestha et al. 2019). In our review, we list 31 species of *Gibellula* and 28 species of *Torrubiella* as spider pathogens (Appendix 1) which is in line with Mendes-Pereira et al. (2023) as concerns the number of accepted *Gibellula* species. So far, spiders from at least 25 families have been observed to be infected by pathogens from the genus *Gibellula* (Appendix 2). Fungi in the genera *Akanthomyces* [= *Verticillium*], *Beauveria, Clathroconium, Clonostachys, Cordyceps, Engyodontium, Granulomanus, Hevansia, Hirsutella, Hymenostilbe, Jenniferia, Lecanicillium, Metarhizium, Neoaraneomyces*, *Ophiocordyceps*, *Parahevansia, Parengyodontium*, *Polystromomyces*, *Pseudogibellula*, *Pseudometarhizium*, *Purpureocillium*, and *Tolypocladium* are also considered to be spider pathogens (Appendix 1; see Edwards 1980; Evans & Samson 1987; Luangsa-ard et al. 2005; Evans 2013).

For the remaining fungal taxa (i.e., *Acrodontium*, *Aphanocladium*, *Apiotrichum*, *Aspergillus, Basidiobolus*, *Cladosporium, Conidiobolus, Cryptococcus* [= *Filobasidiella*]*, Mucor, Penicillium, Sporodiniella* etc.) their status as pathogens or saprobes is often still controversial or undefined (see Appendix 1; Malloch et al. 1978; Greif & Currah 2007; Yoder 2009; Bibbs et al. 2013; Evans 2013; Heneberg & Rezac 2013; Henriksen et al. 2018; Shrestha et al. 2019). Noordam et al. (1998: p. 346) commented on such cases, stating that “*their biology remains unknown; possibly they are essentially saprophytes, with a facultative parasitism on litter-inhabiting invertebrates*”. In line with this, Evans (2013) termed fungi of the genera *Conidiobolus* and *Mucor* associated with spider hosts as “opportunistic pathogens” (= “facultative pathogens” *sensu* Noordam et al. 1998). The category of unclassified fungi includes also fungal hyphae attached to extinct dead spiders found as inclusions in Baltic amber, although there is a strong suspicion that in such cases the fungi in question are most often saprobes (Wunderlich 2004). One of the fungi, growing on an extinct spider, has been identified to be *Arachnomycelium filiforme* (see Wunderlich 2004).

No fungi pathogenic to spiders are known from the myxomycetes (slime molds) (see Nentwig & Prillinger 1990) for the simple reason that these organisms feed on bacteria and other microorganisms (Keller et al. 2008). Because of that the myxomycetes were not included in Appendix 1. Nevertheless, we would like to briefly mention that associations between spiders and myxomycetes do occur. For example, a sporangium of the myxomycete *Licea poculiformis* was found on the leg of a dead spider, and in another instance a myxomycete was observed sporulating on a spider’s orb-web (Ing 1994; Keller et al. 2008).

### What infection mechanisms are used by spider-pathogenic fungi?

The infection mechanism differs between araneomorph above-ground spiders (Figs. 3–5) and mygalomorph ground-burrowing spiders (Fig. 6). The infection mechanism of araneomorph subterranean spiders (Fig. 2A–E) is similar to that of araneomorph above-ground spiders.

#### Infection mechanism in araneomorph above-ground spiders

Above-ground spider hosts were commonly found on elevated sites attached to the underside of leaves (Fig. 2F), where temperature and humidity conditions are optimal for fungal growth (Humber & Rombach 1987; Sanchez-Pena 1990; Samson & Evans 1992; Hywel-Jones 1996; Tzean et al. 1997; Andersen et al. 2009; Brescovit et al. 2019; Kuephadungphan et al. 2019, 2022; Mongkolsamrit et al. 2022). Discharged from the spider cadavers, the spores disperse rapidly over a wide area potentially infecting many spiders perched on lower leaves. Once the spores adhere to a spider’s cuticle, they germinate, and germ tubes penetrate the exoskeleton, intruding into the spider’s body cavity (Kuephadungphan et al. 2022). The fungus can make use of openings or weak spots in the exoskeleton (i.e., mouth, anus, booklung stigma, genital opening, intersegmental membranes at the leg joints, or directly through the thin abdominal cuticle) as entry points for the infection (see Yoder 2009; Martynenko et al. 2012; Kuephadungphan et al. 2022). In medium-to larger-sized araneomorphs, the abdomen appears to be the body part most vulnerable during the premature stages of a fungal infection (Figs. 3–5), whereas in small-sized araneomorphs fungal infections were most commonly first visible on the legs (Bishop 1990a; Noordam et al. 1998). We know from insect model systems that the pathogen penetrates the cuticle using lipases, proteases and chitinases (see St. Leger et al. 1998; Charnley 2003). The mechanisms used by pathogenic fungi to infect insects and araneomorph spiders apparently are very similar (also see Brescovit et al. 2019; Kuephadungphan et al. 2022). In the living arthropod host (insect, spider etc.) the fungus develops in the hemolymph. This process can last from a few days to months or years. This depends largely on the life-cycle of the host. In the hemolymph the fungus feeds on the nutrients that the host has present. As the yeast cell numbers increase the hemolymph becomes compromised leading to the death of the host. At this point there will be very little fungus material in the cephalothorax which is largely muscle tissue. The abdomen by contrast contains a large volume of organs bathed in hemolymph. Critical is the spider cuticle. The cuticle of the cephalothorax is heavily sclerotised with exocuticle – especially the carapace. The legs also are heavily sclerotised to give them rigidity. The abdomen, however, has a much thinner cuticle lacking an exocuticular layer. Similarly, although the leg segments are sclerotised the joints are non-sclerotised. Having killed the spider the fungus in the hemolymph rapidly switches to a mycelial form. This way it can consume the organs in the abdomen and begin to invade the muscle tissue of the cephalothorax and legs (Nigel Hywel-Jones, pers. obs.). This is why the external fungus is first visible on the soft (non-sclerotised) cuticle of the abdomen and leg joints. It has been hypothesized that – once the fungus invades the brain – the spider’s behavior is manipulated (“zombie spiders”) causing it to climb the plant to an elevated site where it will die on a leaf underside (Fig. 2F; Shrestha et al. 2019; Arruda 2020; Arruda et al. 2021; Kuephadungphan et al. 2022; Saltamachia 2022). Much of our perception of spider behavioral manipulation has been inferred from extensive research done on fungus-infected ants. The approach of comparing fungus-infected ants and spiders with each other is reasonable since it appears that the two arthropod groups show the same type of altered host behavior (see Samson et al. 1988; Andersen et al. 2009; de Bekker et al. 2014; Loreto & Hughes 2019). After the death of the spider, the fruiting bodies burst out the cadaver (Brescovit 2019). The behavior of pursuing elevated sites on the underside of leaves for spore release can be seen as an adaptation to optimize the fungal reproductive success (Fig. 2F; Jensen et al. 2001; Andersen et al. 2009; Hughes et al. 2016).

#### Infection mechanism in mygalomorph ground-burrowing spiders

Below-ground spider hosts are for the most part trapdoor spiders from various families, although other types of mygalomorphs get infected by the same group of fungi (Fig. 6). Fungal pathogens engaged in the infection of ground-burrowing spiders are several species in the sexually reproductive genus *Cordyceps sensu lato* (Fig. 6B–E), on the one hand, and the asexual morph *Purpureocillium atypicola* (Fig. 6A), on the other hand. Reports on mycosed trapdoor spiders originate predominantly from South America and Asia (Mains 1954; Kobayasi & Shimizu 1977; Coyle et al. 1990; Haupt 2002; Evans 2013; Hughes et al. 2016; Chirivi et al. 2017).

The infection of spiders by the sexual morphs of *Cordyceps* unfolds as follows: sexual propagules (ascospores) adhere to the spider’s cuticle, whereupon germ tubes are formed that penetrate the cuticle and enter the body cavity similar to the way described for araneomorph spiders. Again, yeast-like hyphal bodies develop in the hemolymph and reproduce, gradually replacing the host tissue which ultimately leads to the spider’s death. The fruit-body (stroma), up to 10 cm in length, emerges from the spider cadaver, grows along the burrow, and pushes through the trapdoor to facilitate the aerial dispersal of the forcibly-ejected ascospores through specialised pores at the tip of the spore-containing sac or ascus (Figs. 6B–D; Kobayasi & Shimizu 1977; Evans 2013; Hughes et al. 2016; Chirivi et al. 2017).

The infection of spiders by the asexual morph *Purpureocillium atypicola* plays out similar to that of *Cordyceps*, except that in this latter case fungal reproduction takes place by means of asexual spores (conidia) produced at the upper end of the fruit body (synnema; Fig. 6A). The conidia are dispersed aerially by wind currents (Coyle et al. 1990; Haupt 2002).

Evans (2013) noted that in the case of trapdoor spiders the question, how the aerially dispersed spores (ascospores and conidia) reach their target (i.e., the spider cuticle), remains an unresolved mystery given that trapdoor spiders live in their burrows beneath the soil surface with little exposure to airborne spores. But the hidden, sedentary life only applies to the female trapdoor spiders, whereas the males likely come into contact with airborne spores while wandering around on the soil surface in search of female burrows during the mating season (see Schwendinger 1991; Bond & Stockman 2008). [It must be added that male and female spiders alike get infected by fungal pathogens (Noordam et al. 1998; Gonzaga et al. 2006; Brescovit et al. 2019)]. This suggests possible sexually transmitted fungal infections (i.e., spores being transferred from an infected male to a healthy female during mating encounters). The possibility of spider-to-spider transmission of fungal pathogens had already been suggested by Henschel (1998). Alternatively, there is a possibility that in some trapdoor spiders known to disperse as tiny juveniles by ballooning (see Coyle et al. 1985; Buzatto et al. 2021) those could get exposed to fungal spores after leaving the mother’s burrows similar to the situation observed in immature araneomorphs (see Bishop 1990a). For comparison, in cicadas, another arthropod group parasitized by fungi from the genus *Cordyceps*, which develop in burrows up to two meters deep, the infection is also picked up by the immature stage before burrowing into the soil (Nigel Hywel-Jones, pers. obs.). A further possibility is that ants with fungal spores attached to their cuticle act as carriers of spores while wandering from above-ground to below-ground areas (see Bibbs et al. 2013). Still another possibility is that spiders could be exposed to fungi from their arthropod food sources (Bibbs et al. 2013).

**Diversity of spider families parasitized by pathogenic fungi**.—Our survey revealed that more than 40 out of the currently accepted 135 spider families (ca. 30%) contain species which are attacked by fungal pathogens (Fig. 1; Table 2; Appendix 2). The majority (roughly 90%) of the reported fungus-infected spiders belong to the suborder Araneomorphae (Figs. 2– 5; see Supplementary Table S1). The list of araneomorph families known to be fungus-infected under natural conditions includes the Agelenidae, Amaurobiidae, Anyphaenidae, Araneidae, Arkyidae, Cheiracanthiidae, Clubionidae, Corinnidae, Ctenidae, Cybaeidae, Cycloctenidae, Deinopidae, Desidae, Dysderidae, Eresidae, Hahniidae, Hersiliidae, Hypochilidae, Lamponidae, Linyphiidae, Liocranidae, Lycosidae, Nephilidae, Oonopidae, Oxyopidae, Philodromidae, Pholcidae, Pisauridae, Salticidae, Sicariidae, Sparassidae, Tetragnathidae, Theridiidae, Thomisidae, Trachelidae, Trechaleidae, Uloboridae, and Zodariidae (Fig. 1; Table 2).

Only roughly 10% of the documented spider mycoses are attributed to the Mygalomorphae or Mesothelae (Fig. 1; Table 2; Supplementary Table S1). The list of fungus-infected mygalomorph/mesothele families includes the Actinopodidae, Atracidae, Atypidae, Barychelidae, Ctenizidae, Dipluridae, Euctenizidae, Halonoproctidae, Heptathelidae, Idiopidae, Nemesiidae, Porrhothelidae, Pycnothelidae, and Theraphosidae (Fig. 1; Table 2).

Apart from these naturally occurring fungal infections, laboratory infections by fungi have been reported for the following spider families not mentioned so far: Antrodiaetidae, Dictynidae, and Ischnothelidae (Table 2; Appendix 2). It can be expected that these latter spider families are parasitized by fungi under natural conditions as well. Furthermore, it is worth mentioning that fungal hyphae attached to spider cadavers encased in samples of Baltic amber were reported in the literature, whereby this refers to the spider families Corinnidae, ‘Gnaphosidae’, Hahniidae, Mysmenidae, Synotaxidae, and Zodariidae (Wunderlich 2004). Although the fossil species of “amber spiders” in question are considered to be extinct (Wunderlich 2004), the six families, to which they belong, are still extant today (see World Spider Catalog 2023).

Jumping spiders (Salticidae), cellar spiders (Pholcidae), and sheet-web spiders (Linyphiidae) are the spider families most frequently reported to be infected by fungal pathogens, these three families combined being accountable for >40% of all documented spider mycoses (Fig. 1). Each of these families is typically associated with a particular environment. Salticids (Figs. 4A–D) were the dominant fungus-infected above-ground spiders of the Tropics/Subtropics and of North America (Evans & Samson 1987), whereas linyphiids (Fig. 3) prevailed among the fungus-infected above-ground spiders in Europe (Bristowe 1948, 1958; Duffey 1997; Noordam et al. 1998; Meyling et al. 2011). Fungus-infected pholcids (Figs. 2A–E) are very common in subterranean spaces such as basements, caves and tunnels in Europe and North America (Martynenko et al. 2012; Kathie T. Hodge, pers. comm.).

### Geographic differences in the percent composition of dominant spider families parasitized by pathogenic fungi

Fungal infections of spiders have been observed in the geographic belt between latitude 78°N and 52°S (Bristowe 1948; O’Donnell et al. 1977). That is, on all continents except Antarctica, and with spider mycoses occurring in very different climatic regions. Spider-pathogenic fungi have excellent long-distance dispersal capabilities taking advantage of their spider hosts’ powerful means of dispersal (Bishop 1990a). On board of parasitized immature spiders such fungi can theoretically “balloon” over distances of 100– 200 kilometres (see Decae 1987; Bishop 1990a). There seems to be an exception to this in the case of fungus-infected subterranean pholcids (*Pholcus* species) in Europe and North America; most species from this particular family apparently do not disperse via ballooning (Schäfer et al. 2001; Simonneau et al. 2016). In Fig. 7, the parasitization of spiders by pathogenic fungi in different geographic regions is compared.

Fig. 7 shows that intercontinental differences in the frequency with which different fungus-infected spider groups were reported exist. The following points are notable:

1. Salticidae is overall the above-ground spider family most frequently reported as hosts of fungal pathogens in the Tropics/Subtropics and North America (Fig. 7). The distinctive shape of the cephalothorax makes these spiders identifiable even when completely covered by mycelium. Statistically, the percentage of fungus-infected salticids in the Tropics/Subtropics is not significantly different from that in North America (23% vs. 28%; Chi-square test, χ*^2^*= 0.730, *df* = 1, *P* > 0.05). The predominance of the salticids among the fungus-infected spiders may be explained by the fact that salticids: (i) are among the most common spiders in warmer areas, thus, making available a huge number of potential hosts for fungal attacks (Nelson et al. 2004); and (ii) that these spiders are highly exposed to the attacks by airborne fungal spores while searching the plant surface for prey or potential mates during daylight hours (Evans & Samson 1987). Average annual temperatures in the cold/temperate zones of Europe are significantly lower compared to North America and the Tropics/Subtropics which explains why salticids as thermophilic animals are less common in Europe (see Nyffeler & Sunderland 2003). The much lower availability of salticids as potential hosts for pathogenic fungi might explain why this spider family is apparently less targeted by fungi in this part of the globe. So far, published reports of fungus-infected salticids from Europe are lacking (Fig. 7).
2. Linyphiidae are one of the most frequently reported spider groups parasitized by fungal pathogens in Europe but not in the Tropics/Subtropics and North America (Fig. 7). Statistically, the percentage of linyphiids infected by fungi in Europe is significantly higher than that in the Tropics/Subtropics (27% vs. 2%; Chi-square test, χ*^2^* = 40.130, *df* = 1, *P* < 0.0001) or in North America (27% vs. 3%; Chi-square test, χ*^2^* = 18.997, *df* = 1, *P* < 0.0001). This can be explained by the fact that linyphiids, which are well-adapted to moderate and colder temperate climates, are among the most common, diverse and widespread above-ground spiders across large parts of Europe (Bristowe 1958; Nyffeler & Sunderland 2003). As opposed to this, only relatively few reports on fungus-infected linyphiids are known from the Tropics/Subtropics and North America, two geographic regions where linyphiids are less common (Nyffeler & Sunderland 2003).
3. Pholcidae are among the spider groups most frequently reported to be attacked by fungal pathogens in North America and Europe (Fig. 7). Statistically, the percentage of fungus-infected pholcids in North America does not significantly differ from that in Europe (30% vs. 36%; Chi-square test, χ*^2^* = 0.917, *df* = 1, *P* > 0.05). The reported infected pholcids have a subterranean life style, inhabiting caves, tunnels, and basements in buildings (Eiseman et al. 2010; Martynenko et al. 2012; Humber et al. 2014). Such subterranean spaces maintain a microclimate characterized by high relative humidity (98–100% RH) and moderate temperatures which are optimal for fungal growth (see Yoder 2009; Martynenko et al. 2012). The spiders most frequently reported in this context are cellar spiders in the genus *Pholcus* Walckenaer, 1805 (i.e., *Pholcus phalangioides* (Fuesslin, 1775)) (Fig. 2A–E; Keller 2007; Martynenko et al. 2012; Jent 2013; Humber et al. 2014). In North America and Europe, fungi isolated from mycosed pholcids belonged exclusively to the family Cordycipitaceae (Appendix 1–2). The following fungi were isolated from subterranean pholcids: the asexual morphs *Engyodontium aranearum* [= *Lecanicillium tenuipes*]*, E. rectidentatum*, and *Parengyodontium album* [= *Beauveria alba = Engyodontium album*] as well as the sexual morph *Torrubiella pulvinata* (Cokendolpher 1993; Keller 2007; Eiseman et al. 2010; Martynenko et al. 2012; Jent 2013; Humber et al. 2014; Dubiel 2015; Kubátová 2017; Kathie T. Hodge, pers. comm.). There are few reports of fungal infections on subterranean spiders from tropical/subtropical regions. In a cave in Cuba a pholcid in the genus *Modisimus* Simon, 1893 was infected by a *Mucor* sp. (Mucoraceae; Mercado et al. 1988). Statistically, the percentage of pholcids infected by fungi in the Tropics/Subtropics is significantly lower than that in North America (3% vs. 30%; Chi-square test, χ*^2^* = 40.048, *df* = 1, *P* < 0.0001) or Europe (3% vs. 36%; Chi-square test, χ*^2^* = 57.098, *df* = 1, *P* < 0.0001). We cannot rule out that the low percentage of subterranean pholcids in the Tropics/Subtropics is biased in the sense that fungus-infected pholcids from tropical/subtropical subterranean basements and caves may have been less frequently investigated so far. Furthermore, the situation may be different with regard to free-living pholcids in the Tropics/Subtropics. For example, in one study from Brazil, plant-dwelling pholcids (genus *Metagonia* Simon, 1893) were found to be the spider family most frequently attacked by fungi from the genus *Gibellula* (27% of all fungus-infected spider specimens; Costa 2014). Such a plant-dwelling pholcid from Brazil (*Metagonia taruma* Huber, 2000) is depicted in Fig. 2F.
4. Another notable point is that in the Tropics/Subtropics both araneomorph and mygalomorph spiders are routinely attacked by fungal pathogens, whereas reports of fungus-infected mygalomorphs from Europe and North America are very scarce (≤5% of all reports; Fig. 7). For example, in Southwestern Europe only one mycosed trapdoor spider per ca. 60–70 burrows was found (Vera Opatova, pers. comm.). Statistically, the percentage of mygalomorphs infected by fungi in the Tropics/Subtropics is significantly higher than that in North America (18% vs. 3%; Chi-square test, χ*^2^*= 11.315, *df* = 1, *P* < 0.001) or Europe (18% vs. 5%; Chi-square test, χ*^2^* = 8.937, *df* = 1, *P* < 0.005). Most reports of fungal attacks on mygalomorphs refer to trapdoor spiders in the families Actinopodidae, Barychelidae, Halonoproctidae, Idiopidae, and Pycnothelidae (Coyle et al. 1990; Schwendinger 1996; Haupt 2002; Chirivi et al. 2017; Manfrino et al. 2017; Pérez-Miles & Perafan 2017; and others).

### Are spider-pathogenic fungi host-specific?

The host specificity of spider-pathogenic fungi varies considerably between the different pathogen species. Evans (2013: p. 109) suggested that many spider-pathogenic fungi are highly specific and “*that this specificity must operate at the exoskeleton level*”. Some fungal pathogens are host specific to the extent that they have been found so far from only one spider host (Kuephadungphan et al. 2022). Spider-pathogenic fungi of this type include a newly discovered fungus with the epithet *Gibellula ‘bang-bangus’*, *G. brevistipitata*, *G. cebrennini*, *G. dimorpha*, *G. fusiformispora*, *G. longicaudata*, *G. longispora*, *G. nigelii*, *G. parvula, G. pilosa*, *G. pigmentosinum*, *G. scorpioides*, *G. solita*, *G. unica*, *Jenniferia cinerea*, *J. griseocinerea*, *J. thomisidarum*, and others (see CABI 2022; Kuephadungphan et al. 2022; Mongkolsamrit et al. 2022). Thus, based on current knowledge, these fungi appear to have a very narrow host range. However, due to the fact that such assessments are often based on a very low number of observations, it can not be ruled out that in some of these cases a broader host range will come to light in the future, once a larger number of host records has been made available.

Others are spider-specific fungal pathogens with a wide host range across many spider families. *Purpureocillium atypicola* is perhaps the most prominent example of a spider-specific fungal pathogen with a wide host range. Spiders from ten families could be infected with *P. atypicola* under laboratory conditions (Greenstone et al. 1987). The fact that this fungus is utilizing a wide range of hosts under laboratory conditions has been verified by our survey according to which spiders from at least 17 families are infected by *P. atypicola* under natural conditions (Appendix 2). Other examples of fungi which infect a range of hosts from different spider families are *Gibellula leiopus* and *G. pulchra*, two species with a broad geographic distribution (Samson & Evans 1973; Strongman 1991; Bałazy 1970; Kubátová 2004; Selçuk et al. 2004; Zare & Zanganeh 2008; Dubiel 2015; Savić et al. 2016; Shrestha et al. 2019). Based on our survey, these two very common fungal pathogens utilize seven different spider families as hosts in the case of *G. leiopus* and eight families in the case of *G. pulchra* (Appendix 2). *Gibellula leiopus* found on very different spider hosts may actually be a *G. leiopus* species complex that needs to be disentangled by future research (Harry Evans, pers. comm.; Nigel Hywel-Jones, pers. obs.). It seems that exclusively araneomorph spiders are attacked by *Gibellula* species (Shrestha et al. 2019; Kuephadungphan et al. 2022). By way of contrast, mygalomorph spiders, whose cephalothorax is covered with a strong exoskeletal cuticle, apparently remain unaffected from attacks by *Gibellula* species (Shrestha et al. 2019). Nevertheless, attacks by fungi from the genus *Gibellula* are apparently not strictly limited to araneomorph spiders, because Brazilian researchers recently reported on a harvestman of the family Stygnidae being infected by a *Gibellula* species (Villanueva-Bonilla et al. 2021).

Apart from these spider-specific fungal pathogens, there are other pathogens with lower host specificity, which infect spiders and other arthropods alike. An example of a fungal taxon with an extremely wide host range is *Beauveria bassiana* known to attack over 700 species in 15 insect orders in addition to spiders and mites (Sosnowska et al. 2004; Cummings 2009; Meyling et al. 2011).

## DISCUSSION

### How many species of spider-pathogenic fungi exist worldwide?

Fungi in general and spider-pathogenic fungi in particular remained understudied (Boomsma et al. 2014). Only about 5% of all extant fungal species have been described so far (see Hawksworth & Rossman 1997; Bhalerao et al. 2019). If this figure is used to estimate by extrapolation the number of still undiscovered spider-pathogenic fungi, then it follows that >1,000 unnamed species might exist. If we are only talking about the genus *Gibellula*, it was suggested that this genus alone probably contains hundreds of unnamed species (Harry Evans, pers. comm.). In fact, new species of spider-pathogenic fungi are being discovered all the time (e.g., Han et. al. 2013; Shrestha et al. 2019; Chen et al. 2017, 2018, 2021, 2022a,b; CABI 2022; Kuephadungphan et al. 2022; Mendes-Pereira et al. 2022; Mongkolsamrit et al. 2022; Tan et al. 2022; Zhou et al. 2022; Chen et al. 2023; Mongkolsamrit et al. 2023; Wang et al. 2023b).

Furthermore, because the biology of a large number of fungal species is still unexplored, their status as pathogens or saprobes is currently unknown or controversial (see Noordam et al. 1998; Evans 2013). So it could be that some species currently classified as saprobes are in fact facultative pathogens which would further increase the true number of existing species of spider-pathogenic fungi.

### Low-investment vs. high-investment reproductive strategy in hypocrealean fungi

A comparison between fungus-infected above-ground spiders (araneomorphs) with fungus-infected ground-burrowers (mygalomorphs) shows that the fungi attacking these two types of spiders differ in their reproductive patterns in terms of energy expenditure. The majority of fungus-infected above-ground spiders are small in size, often measuring only ca. 1–5 mm in body length (Mains 1939; Bristowe 1948, 1958; Bishop 1990a,b; Noordam et al. 1998; Meyling et al. 2011; Brescovit et al. 2019). Spiders of that size have a body mass of ca. 1–5 mg according to Nyffeler & Sunderland (2003). For comparison, the ground-burrowing trapdoor and purse-web spiders are much larger (ca. 10–35 mm in body length) (Coyle et al. 1990; Bellmann 1997; Ono 2001; Ríoríos-Tamayo & Goloboff 2018) with a body mass of ca. 500–2,500 mg per spider (Anderson 1987; Bradley 1996; Hardy 2018). Ground-burrowing theraphosids can reach a body mass of up to 30,000 mg (Baerg 1963). The amount of energy which can be extracted by a fungal parasite from a large burrowing spider host is roughly two to four orders of magnitude larger than that extracted from a small above-ground host. Thus, fungi parasitizing above-ground spiders (araneomorphs) exhibit a low-investment reproductive strategy as opposed to the fungi found on the ground-burrowers (mygalomorphs). The fungi parasitizing ground-burrowers invest a much higher amount of energy into their reproductive effort (also see Evans & Samson 1987). Subterranean pholcids such as *Pholcus phalangioides* are also relatively small araneomorphs, with a body mass of ca. 10–30 mg (Schmitz 2015). The fungi which attack pholcids (i.e., *Engyodontium aranearum*, *E. rectidentatum*, *Parengyodontium album*, and *Torrubiella pulvinata*) also exhibit a low-investment reproductive strategy (Fig. 2A–E).

The low-investment strategy of the fungi parasitizing small above-ground spiders – such as *Gibellula* species – works in such a way that a spider is manipulated to climb a plant, whereupon spores are released from tiny fruit bodies growing on the dead spider body (Fig. 2F). This procedure requires little energy, taking into account that small spiders are excellent climbers which due to their low body mass have to exert little resistance against the force of gravity and that little energy needs to be invested to produce the tiny fruit bodies.

For the ground-burrowing spiders, the situation is completely different. Fungi parasitizing mygalomorphs – usually found in the genera *Cordyceps* and *Purpureocillium* – die in their underground burrows and then use phototrophic fruit bodies to push above ground (Fig. 6A– E; Cummings 2009; Hughes et al. 2016; Rowley et al. 2022). These types of fungi must produce stout, fleshy fruit bodies strong enough to push through the soil (i.e., theraphosids) or to open trapdoors (i.e., trapdoor spiders) and which are long enough to emerge several centimeters above the ground surface to facilitate aerial dispersal of the released spores (Fig. 6 A–E). The production of such stout fruiting bodies is costly in energy. Only heavy built spider hosts – usually mygalomorphs – provide a sufficiently large amount of energy to facilitate the costly production of this type of reproductive structure (Fig. 6E).

### In which habitats do spider mycoses occur most frequently?

High rates of fungus infection were documented for above-ground spiders in tropical/subtropical forested areas (including national parks and wildlife sanctuaries) in Brazil, China, Ghana, Indonesia, and Thailand (Tables 3–4; Nentwig 1985a; Evans & Samson 1987; Humber & Rombach 1987; Samson & Evans 1974, 1982, 1992; Rong & Botha 1993; Tzean et al. 1997; Selçuk et al. 2004; Aung et al. 2006; Shrestha et al. 2019; Kuephadungphan et al. 2019, 2022; Mongkolsamrit et al. 2022). As Aung et al. (2008) point out, fungal pathogens show their highest dominance in humid tropical forests. Cacao farms in Ecuador and Ghana are further tropical habitats in which high rates of fungus infection were observed (Table 3–4; Evans & Samson 1987). Citrus, coffee, guava, rice, and sugar-cane plantations are other tropical/subtropical habitats in which spider-pathogenic fungi were found (Table 3). According to several field assessments, infection rates by hypocrealean fungi of ca. 10–35% were observed for tropical spider populations, although there were also exceptions to this with infection levels of ≤1% (Table 4).

**Table 3.** Diversity of habitat types from which spider mycoses had been reported (based on literature).

| Habitat type | Tropics / Subtropics | North America | Europe | Source |
| --- | --- | --- | --- | --- |
| Tropical & subtropical forests (including National Parks and Wildlife Sanctuaries) | ● |  |  | Samson & Evans 1973; Evans 1974; Nentwig 1985a; Selçuk et al. 2004; Brescovit et al. 2019; Arruda 2020; Saltamachia 2022; Zhou et al. 2022; Vandegrift et al. 2023 |
| Atlantic forest | ● |  |  | Gonzaga et al. 2006; Costa 2014; Mendes-Pereira et al. 2022 |
| Scalesia forest | ● |  |  | Evans & Samson 1982 |
| Southern deciduous forest |  | ● |  | Bishop 1990a |
| Temperate forests |  | ● | ● | Mains 1939; Bałazy 1970; Kubátová 2004; Tkaczuk et al. 2011; Robb Bennett, pers. comm. |
| Nature Parks, Wildlife Preserves (outside the Tropics & Subtropics) |  | ● |  | https://bugguide.net/node/view/492026/bgimage; https://bugguide.net/node/view/1079720/bgimage |
| Tropical bracken vegetation and moorland | ● |  |  | Evans & Samson 1982 |
| Cacao plantation | ● |  |  | Samson & Evans 1973; Evans & Samson 1987 |
| Citrus plantation | ● | ● |  | Muma 1975; Sanchez-Pena 1990 |
| Coffee plantation | ● |  |  | Wolcott 1948 |
| Guava plantation | ● |  |  | Johnston 1915 |
| Rice field | ● |  |  | Heinrichs 1994; Hywel-Jones 1996 |
| Sugar-cane plantation | ● |  |  | Williams 1921; Evans & Samson 1987 |
| Vegetable field | ● |  | ● | Meyling et al. 2011; Chen et al. 2017 |
| Meadows |  |  | ● | Ellis 1956; Sosnowska et al. 2004 |
| Rush communities (Juncaceae) |  |  | ● | Sosnowska et al. 2004 |
| Fallows covered by *Festuca pratensis* |  |  | ● | Ryszkowski & Wicherek 1997 |
| Ravine bank | ● |  |  | Coyle et al. 1990 |
| Subterranean caves | ● | ● | ● | Mercado et al. 1988; Yoder et al. 2009 |
| Subterranean tunnels |  |  | ● | Martynenko et al. 2012 |
| Subterranean basements |  | ● | ● | Humber et al. 2014; CABI 2022 |
| Gardens | ● |  |  | Kobayasi 1941 |
| Roadside | ● |  |  | Kobayasi 1941 |
| Fens and swampy areas |  |  | ● | Bristowe 1948, 1958; Ellis 1956; McLean 1993; Duffey 1997 |
| Filterbed |  |  | ● | Duffey 1997 |
| Tree-covered dunes |  |  | ● | Noordam et al. 1998 |
| Arctic island, between stones |  |  | ● | Bristowe 1948, 1958 |
| Aquatic environments – Diving bell spider *Argyroneta aquatica* |  |  | ● | Noordam et al. 1998 |

**Table 4.** Infection rate of spider populations (in %) caused by fungal pathogens based on literature data. N/A = information not available.

| Infected spider taxon | Fungal pathogen | Habitat type | Infection rate (%) | Reference |
| --- | --- | --- | --- | --- |
| **TROPICS / SUBTROPICS** |  |  |  |  |
| *Actinopus* Perty, 1833<br>(Actinopidae) | *Purpureocillium atypicola* | Ravine bank – Argentina | 34.0 [*n* = 50] | Coyle et al. 1990 |
| *Latouchia* Pocock, 1901<br>(Halonoproctidae) | *Purpureocillium atypicola* | N/A – Japan | 18.2 [*n* = 22] | Haupt 2002 |
| *Prothemenops siamensis*<br>Schwendinger, 1991 (Idiopidae) | *Purpureocillium atypicola* | N/A – Thailand | High infection rate | Schwendinger 1996 |
| *Helvibis longicauda* Keyserling, 1891 (Theridiidae) | *Gibellula pulchra* | Atlantic Forest – Brazil | ♂♂ = 19.4 [*n* = 36],<br>♀♀ = 9.4 [*n* = 149],<br>Overall = 11.6 [*n* = 185] | Gonzaga et al. 2006;<br>Cardoso et al. 2018 |
| *Macrophyes pacoti* Brescovit, Oliveira, J. C. M. S. M. Sobczak & J. B. Sobczak, 2019 (Anyphaenidae) | *Gibellula* sp. | Tropical forest – Brazil | 27.2 [*n* = 2,565] | Arruda 2020 |
| Jumping spiders (Salticidae) and others | *Gibellula* spp. + others | Tropical forests and cacao farms – Ghana | 80 spiders killed by fungi in a cacao farm and 120 spiders in a tropical forest | Samson & Evans 1973; Evans 1974; Evans & Samson 1987 |
| Orb-weaving spiders (Araneidae, Nephilidae) | *Purpureocillium atypicola* | Tropical forest – Panama | ≤1.0 [*n* >1800] | Nentwig 1985a,b |
| *Atypena formosana* (Oi, 1977) (Linyphiidae) | *Gibellula leiopus* / *Gibellula pulchra* | Rice fields – Philippines | 1.0 [*n* = N/A] | Heinrichs 1994 |
| **TEMPERATE CLIMATE** |  |  |  |  |
| Ballooning spiders (Araneidae, Linyphiidae, Salticidae, Thomisidae) | *Gibellula* sp. or *Torrubiella* sp. | Southern deciduous forest – USA, Tennessee | 5.0–20.0 [*n* = ≈7 00] | Bishop 1990a,b |
| Small unidentified spiders on leaves | *Cordyceps thaxteri* | Temperate forest – USA, North Carolina | ≈70 spiders killed by the fungus in one spot | Mains 1939 |
| *Centromerus sylvaticus* (Blackwall, 1841) (Linyphiidae) | Unidentified Hyphomycetes | Dune habitat – Netherlands | 20.0–50.0 [*n* = 2,615] | Noordam et al. 1998 |
| Sheet-web spiders (Linyphiidae) | *Gibellula* spp. | Agricultural fields – Denmark | 50.0–85.0 [*n* = 40] | Meyling et al. 2011 |
| Sheet-web spiders (Hahniidae, Linyphiidae) | *Gibellula araneorum* / *Torrubiella albolanata* | Fens and swampy areas – England | Countless spiders killed by fungi | Bristowe 1958; Duffey 1997 |
| Unidentified spiders | *Gibellula pulchra* | Fallows covered by *Festuca pratensis* – France | >70.0 [*n* = high] | Ryszkowski & Wicherek 1997 |
| *Pholcus* sp. (Pholcidae) | *Parengyodontium album* [= *Beauveria alba* = *Engyodontium album*] | Subterranean tunnels, Ukraine | 70.0–75.0 [*n* = high] | Martynenko et al. 2012 |
| **ARCTIC CLIMATE** |  |  |  |  |
| Sheet-web spiders (Linyphiidae) | *Cordyceps* sp. | Arctic Island – Norway | Thousands of spiders killed by fungi | Bristowe 1948, 1958 |

The reports of fungus-infected above-ground spiders in North America refer in particular to temperate forests. Fairly high infection rates were observed in the mountainous area of western North Carolina and eastern Tennessee and in a mixed hardwood forest area in northwestern South Carolina (Mains 1939; Bishop 1990a; Kim Fleming, pers. comm.). Mains (1939) found ca. 70 small spiders parasitized by *Cordyceps thaxteri* at one location. In Bishop’s (1990a) study from Tennessee, 5–20% of immature spiders (i.e., Araneidae, Linyphiidae, Salticidae, and Thomisidae) collected from a forest-meteorology tower were infected by hypocrealean fungi, with a maximum infection rate occurring in autumn (Table 4). A huge number of spiders parasitized by fungi from the *Gibellula leiopus* complex was discovered in a mixed hardwood forest area in northwestern South Carolina (Kim Fleming, pers. comm.). The fungus-infected spiders in this latter study includes among others salticids, thomisids, and trachelids (Fig. 5A). It is noteworthy that about half a dozen cases of fungal-infected trachelids (e.g., *Trachelas tranquillus* (Hentz, 1847)) were reported from wooded areas in North America (Table 2; Appendix 2). The fungal pathogens attacking *Trachelas* spiders were usually *Gibellula* species (Fig. 5A; Appendix 2). *Trachelas* often hides very deep in cracks of dead wood which seems to favor fungal infestation (Tobias Bauer, pers. comm.). It is also worth mentioning that several fungus-infected *Cybaeus reticulatus* Simon, 1886 (Cybaeidae) were found from under rotting wood in old rainforest woods on Queen Charlotte Islands, British Columbia, all of which had fruiting bodies growing out of them (Robb Bennett, pers. comm.). Furthermore, high infection rates of pholcid populations by hypocrealean fungi were observed in basements of old houses and in caves at various locations in USA (Eiseman et al. 2010; Jent 2013; Humber et al. 2014; Kathie T. Hodge, pers. comm.).

The reports from Europe refer to temperate forests, tree-covered dunes, meadows, vegetable fields, fens, and swampy areas as regards above-ground spiders (Table 3). The majority of European above-ground spider hosts infected by hypocrealean fungi were small sheet-weavers (Fig. 3; Bristowe 1958; Duffey 1997; Noordam et al. 1998; Meyling et al. 2011). The fungal pathogens engaged in these infections belonged to the genera *Gibellula* and *Torrubiella* (Bristowe 1958; Duffey 1997) or remained unidentified (Noordam et al. 1998). High infection rates of spider populations by fungi occurred in particular in damp places such as swampy areas in eastern England (Ellis 1956; Bristowe 1958; Duffey 1997) and Wales (McNeil 2012; Harry Evans, pers. comm.). High infection rates of 20–85% have been reported for European above-ground spider populations dominated by sheet-weavers (Table 4).

As in North America, in European subterranean spaces a high percentage of pholcids are infected by hypocrealean fungi (Keller 2007; Martynenko et al. 2012). Martynenko et al. (2012) reported that the infection rates of *Pholcus* populations due to *Parengyodontium album* [= *Beauveria alba* = *Engyodontium album*] in tunnel systems in the Ukraine was ca. 70–75% (Table 4). These tunnels, characterized by high relative humidity, are optimal habitats for spiders to be infected by fungal pathogens.

Fungal infestations of spiders occur even in the cold, Arctic climate of the Island of Jan Mayen, Norway (70°59′N / 8°32′W) where huge numbers of small linyphids were killed by a hypocrealean fungus in the genus *Cordyceps* not further identified (Bristowe (1948, 1958). Bristowe (1958: p. 66) noted “*….when I visited the arctic island of Jan Mayen in 1921, I noticed thousands of the dead bodies of each of the four species comprising its spider fauna. They were covered with a white fungus identified as* Cordyceps*…Now I believe it kills the spiders and is almost the only enemy these spiders have in this cold damp island.*” The only spider-pathogenic species in the genus *Cordyceps* so far known from outside of the Tropics/Subtropics apparently is *C. thaxteri* (see Mains 1939; Savić et al. 2016). However, in non-mycological literature of that time, the name ‘Cordyceps’ was often used to generally describe any fungus infecting an insect or, in this case, a spider.

### Possible ecological implications of spider mycoses for the spiders

In the Tropics/Subtropics and in North America/Europe, above-ground spiders perform important ecosystem services by exerting top-down control of herbivorous insects including numerous crop and forest pests (Nyffeler & Sunderland 2003; Nyffeler & Birkhofer 2017). Samson & Evans (1973) suggested that heavy infestation of above-ground spider populations by hypocrealean pathogens might interfere with the natural suppression of herbivorous insects. Heavy hypocrealean infestation of spiders was reported especially from humid forested areas, sugar-cane and cacao plantations in tropical/subtropical regions, on the one hand, and from forested and swampy areas in temperate climates, on the other hand (Williams 1921; Mains 1939; Bristowe 1958; Samson & Evans 1973; Bishop 1990a; Duffey 1997; Meyling et al. 2011). As pointed out by Samson & Evans (1973), much more data on variables such as infection rates, spider densities, and prey densities must be acquired before final conclusions on potentially disruptive effects of fungal pathogens on spider herbivore suppression can be drawn.

Fungal pathogens such as *Purpureocillium atypicola* and some *Cordyceps* species (Fig. 6) may play a significant role in the regulation of mygalomorph spider populations in tropical/subtropical regions (Coyle et al. 1990; Schwendinger 1996; Haupt 2002). Mygalomorph spiders, for their part, serve as food for the larvae of spider-hunting wasps (Pompilidae). In turn, the adults of such wasps act as pollinators of a variety of different plants (Ollerton et al. 2003; Johnson 2005; Shuttleworth & Johnson 2006, 2009, 2012; Phillips et al. 2021). Whether locally heavy fungal infestation of mygalomorph spider populations indirectly affects the pollination success of spider hunting wasp-pollinated plants is still unknown.

What about the pholcid spiders dwelling in subterranean spaces? Due to their high abundance pholcids are prominent members of subterranean food webs at times heavily attacked by fungal pathogens (Souza-Silva et al. 2011; Martynenko et al. 2012). But the ecological role of subterranean pholcid populations is currently unexplored and the implications of fungal infestations cannot yet be determined.

Finally, we would like to add that fungi may have an adverse effect on spider survival in still another way. The fact is that airborne fungal spores from many different fungal families (e.g., Botryosphaeriaceae, Davidiellaceae, Helotiaceae, Massarinaceae, Microascaceae, Nectriaceae, Phragmidiaceae, Pleosporaceae, Trichocomaceae, Trichosphaeriaceae, and Venturiaceae) are blown by wind into the webs of orb-weaving spiders (Smith & Mommsen 1984; Nyffeler et al. 2016, 2023). The trapped spores are ingested by the spiders along with silk material during the recycling process of old webs prior to the construction of new webs, and the consumption of spores may have a detrimental effect on spider survival because of the presence of noxious secondary compounds in the spores (see Smith & Mommsen 1984; Nyffeler et al. 2023).

## CONCLUDING REMARKS

Our review reveals that spiders from at least 40 families are parasitized by fungal pathogens. This is a much higher number of fungus-infected spider families compared to the <10–20 families reported from previous reviews (see Evans 2013; Costa 2014; Humber et al. 2014; Shrestha et al. 2019; Durkin et al. 2021; Kuephadungphan et al. 2022). The fact that: 1) such a high number of spider taxa are parasitized by pathogenic fungi; and that 2) high infection rates have been reported in numerous literature reports (Table 4; Mains 1939; Bristowe 1958; Samson & Evans 1973; Evans & Samson 1987; Bishop 1990a; Coyle et al. 1990; Schwendinger 1996; Duffey 1997; Noordam et al. 1998; Haupt 2002; Gonzaga et al. 2006; Meyling et al. 2011; Martynenko et al. 2012; Arruda 2020; and others) leads to the conclusion that pathogenic fungi might be among the spiders’ major natural enemies. This conclusion is supported by the widespread occurrence of spider-pathogenic fungi in diverse habitat types over a large area of the globe (from latitude 78°N to 52°S). Our conclusion that fungal infection is an important mortality factor for spiders is at odds with the way spider-pathogenic fungi have previously been covered in most text books on spider biology (see above). With this paper, we wish to raise awareness among arachnologists, mycologists, and ecologists that many ecologically significant trophic links between spider-pathogenic fungi and their spider hosts exist and that it would be worthwhile for them to pursue future arachnological-mycological interdisciplinary research in this very fascinating, yet unexplored frontier between arachnology and mycology.

## ACKNOWLEDGEMENTS

Thanks to Tobias Bauer (State Museum of Natural History Karlsruhe, Germany), Robb Bennett (Royal British Columbia Museum, Canada), Antonio Brescovit (Instituto Butantan, São Paulo, Brazil), Bruce Cutler (University of Kansas, USA), Ansie Dippenaar-Schoeman (University of Venda, South Africa), G.B. Edwards (Curator Emeritus, Florida State Collection of Arthropods, Gainesville, USA), Charles Haddad (University of the Free State, Bloemfontein, South Africa), David Hill (Peckham Society, USA), Hubert Höfer (Staatliches Museum für Naturkunde Karlsruhe, Germany), Matjaz Kuntner (National Institute of Biology, Ljubljana, Slovenia), Yuri Marusik (Institute for Biological Problems of the North, Magadan, Russia), Geoff Oxford (York University, UK), Martín Ramírez (Museo Argentino de Ciencias Naturales, Argentina), Robert Raven (Queensland Museum, Australia), Darrell Ubick (California Academy of Sciences, San Francisco, USA), Rick West (Sooke, British Columbia, Canada), Jörg Wunderlich (Hirschberg, Germany), and Alireza Zamani (University of Turku, Finland) who identified for us fungus-infected spiders based on photographs. We also wish to thank Dmitri Logunov (Manchester Museum, the University of Manchester, UK) for taxonomic information. Furthermore, we wish to thank João Araújo (The New York Botanical Garden, USA), Harry C. Evans (CAB International, UK / Universidade Federal de Viçosa, Brazil), Kathie T. Hodge (Cornell University, USA), Richard A. Humber (USDA / Cornell University, USA), Alena Kubátová (Charles University, Prague, Czech Republic), Jennifer Luangsa-ard (National Center for Genetic Engineering and Biotechnology, Thailand), Bhushan Shrestha (Madan Bhandari University of Science and Technology, Nepal), Joseph Spatafora (Oregon State University, USA), and Daniel Winkler (MushRoaming, Seattle, USA) for assistance in the identification of fungi based on photographs. Likewise, we wish to thank Fernando Pérez-Miles (Universidad de la República de Uruguay, Uruguay), Hirotsugu Ono (National Museum of Nature and Science, Japan), Arthur E. Decae (Natural History Museum Rotterdam, Netherlands), and Vera Opatova (Charles University, Prague, Czech Republic) for information on fungus-infected trapdoor spiders. Leonardo Sousa Carvalho (Universidade Federal do Piauí, Brazil) provided us with informing on a fungus-infected pholcid in Brazil. Jason Dunlop (Museum für Naturkunde, Humboldt University Berlin, Germany) and Harry C. Evans helped us to get access to literature. Thanks to Harry C. Evans for his comments on an earlier draft of this paper. We also acknowledge two anonymous reviewers for their valuable comments and the editors Michael Rix (Queensland Museum, Australia), Deborah Smith (University of Kansas, USA), and Rick Vetter (University of California, Riverside, USA) for their support. Furthermore, we express our gratitude to Andy Henzi (Breitenbach, Switzerland) for his help with the layout of the photographs. Last but not least, the following photographers are greatly acknowledged for granting permission to use their photographs: Gilles Arbour (“NatureWeb”, Quebec, Canada), Seth Ausubel (Tubac, Arizona, USA), Paul Bertner (“Rainforests Photography”, British Columbia, Canada), Leonardo Sousa Carvalho (Piauí, Brazil), Carmen Champagne (Athens, Georgia, USA), Jeffrey J. Cook (Wayne, New Jersey, USA), Tony DeSantis (Scranton, Pennsylvania, USA), Tony Eales (Brisbane, Queensland, Australia), Kim Fleming (Donalds, South Carolina, USA), Mary Jane Hatfield (Iowa, USA), Jeff Hollenbeck (Daytona Beach, Florida, USA), Thomas Læssøe (Copenhagen, Denmark), Lynn Lunger (Pennsylvania, USA), J. Mass (Uxbridge, Ontario, Canada), Nicolai V. Meyling (Copenhagen, Denmark), Steve & Alison Pearson (Airlie Beach, Queensland, Australia), Jens H. Petersen (“MycoKey”, Ebeltoft, Denmark), Lisa Powers (“Froghaven Farm”, Bon Aqua, Tennessee, USA), Carlos M. Silva (Gemunde, Portugal), Daniel Winkler (“MushRoaming”, Seattle, Washington, USA), and Kim Wood (Cottage Grove, Wisconsin, USA).

---

**Appendix 1.**
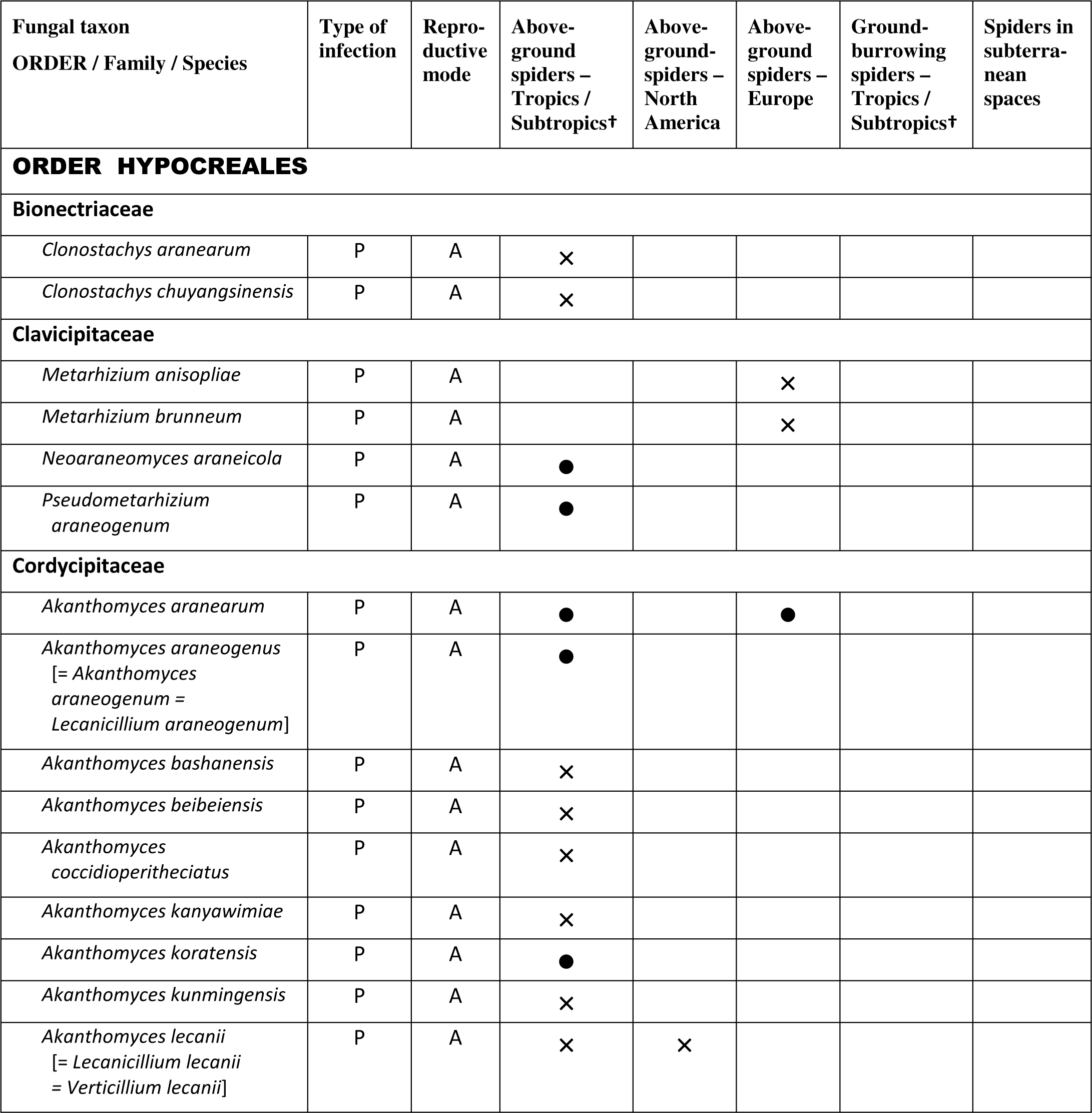
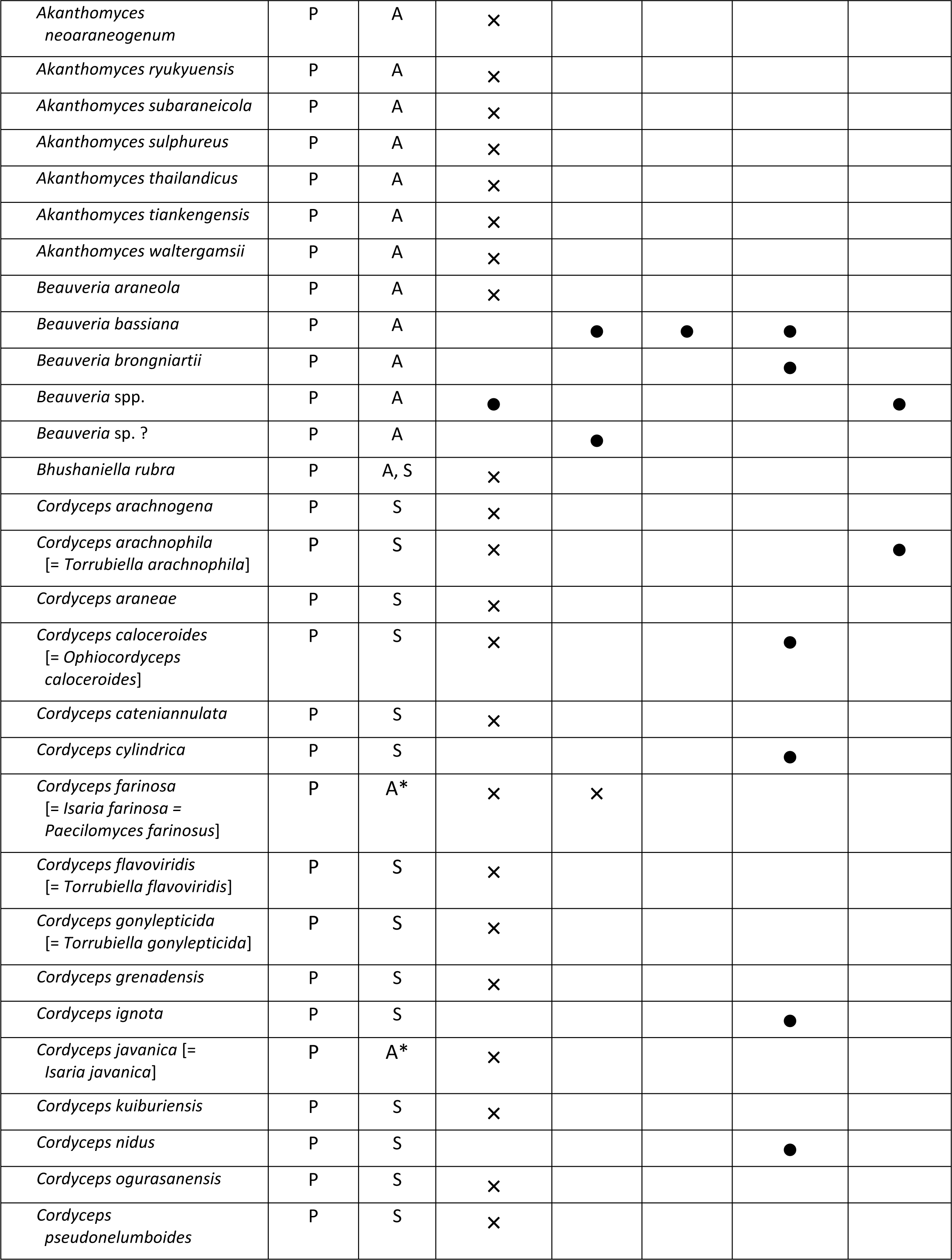
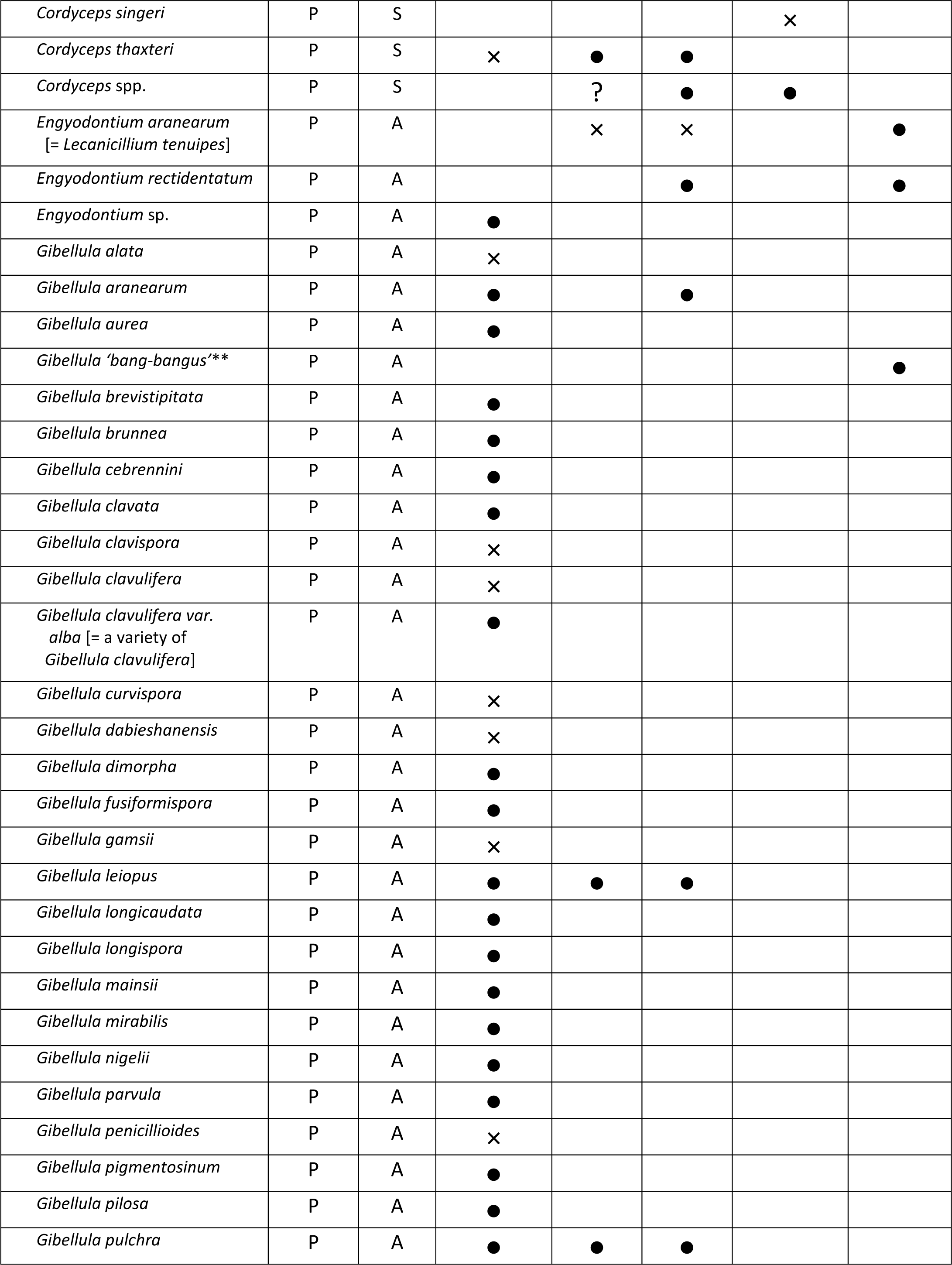
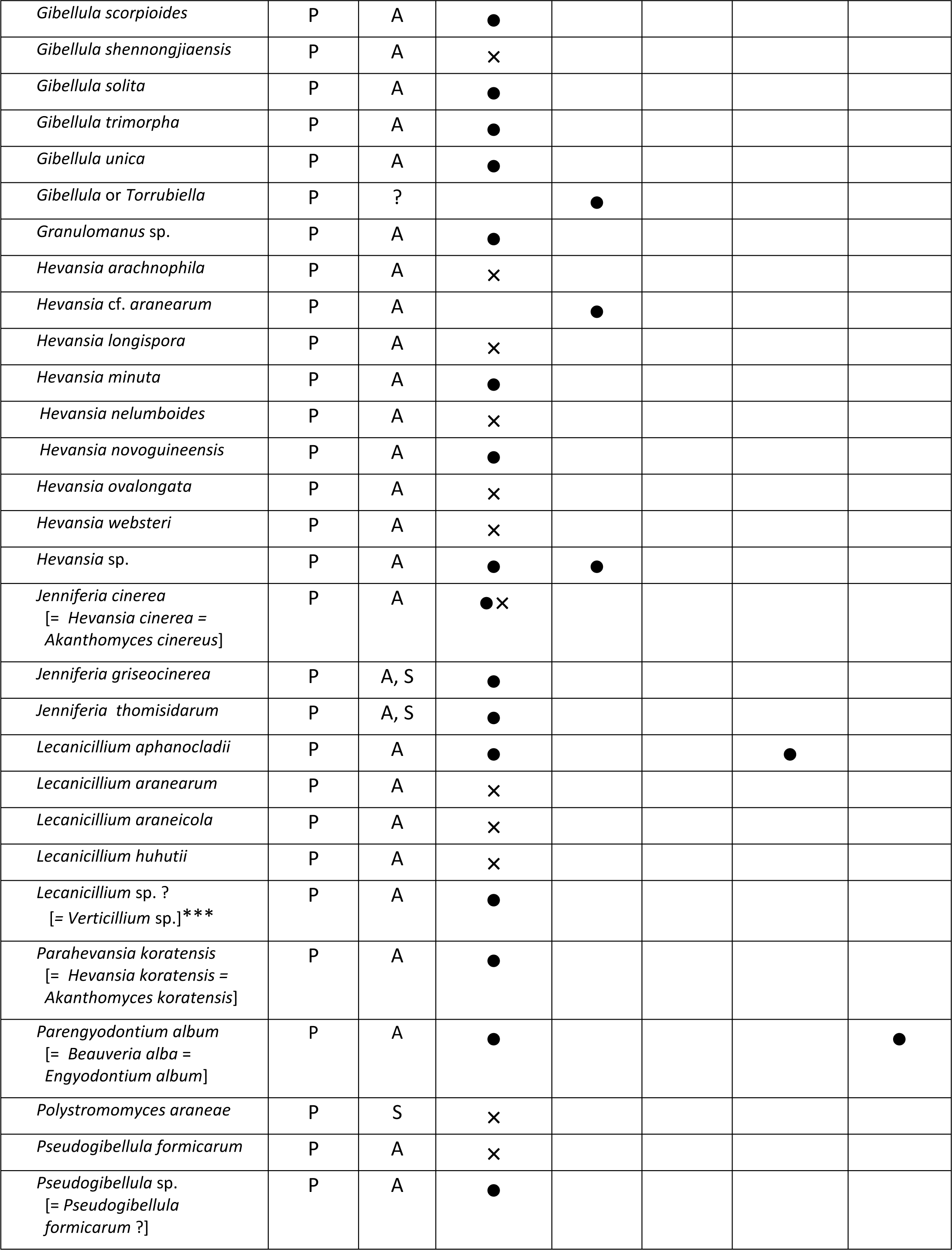
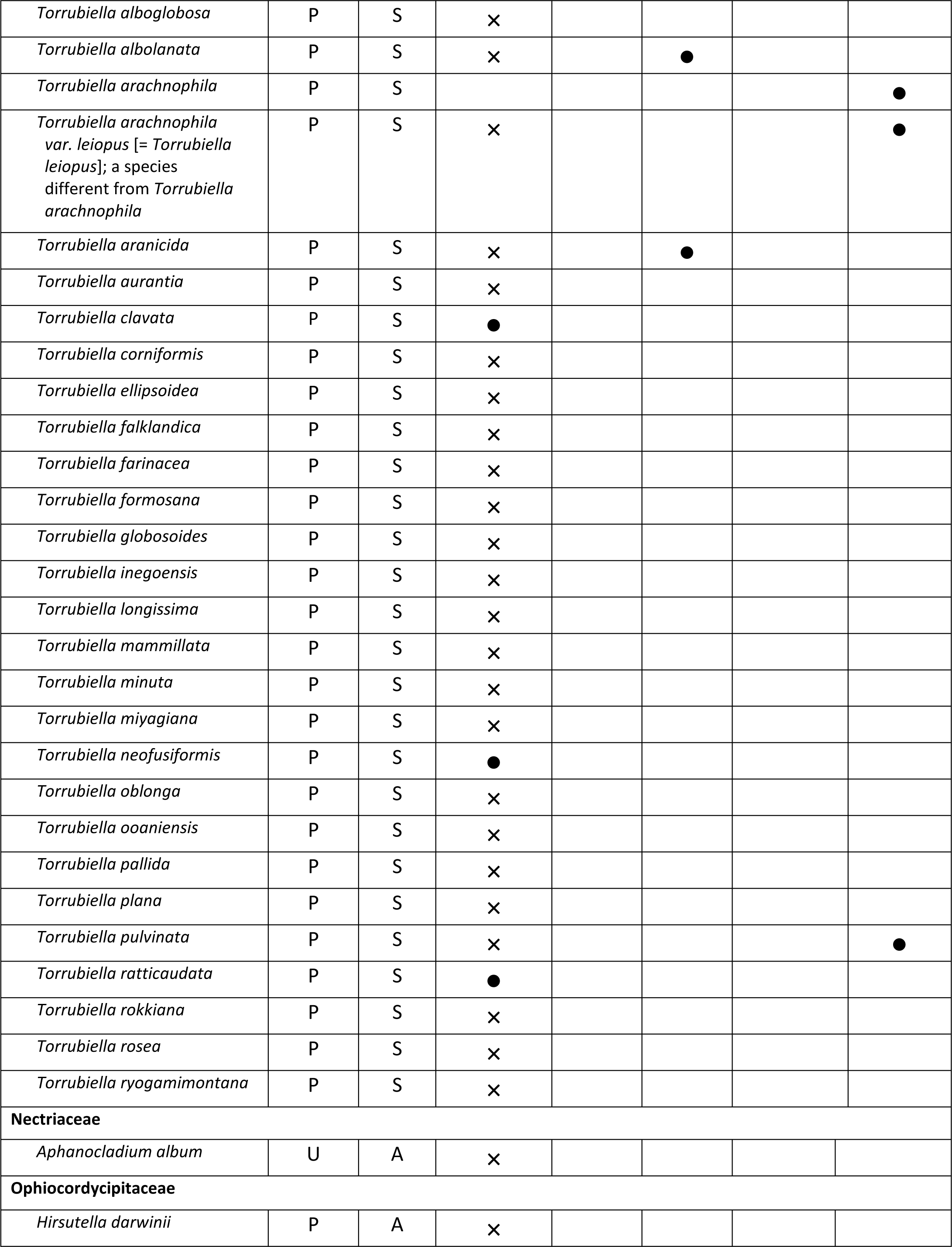
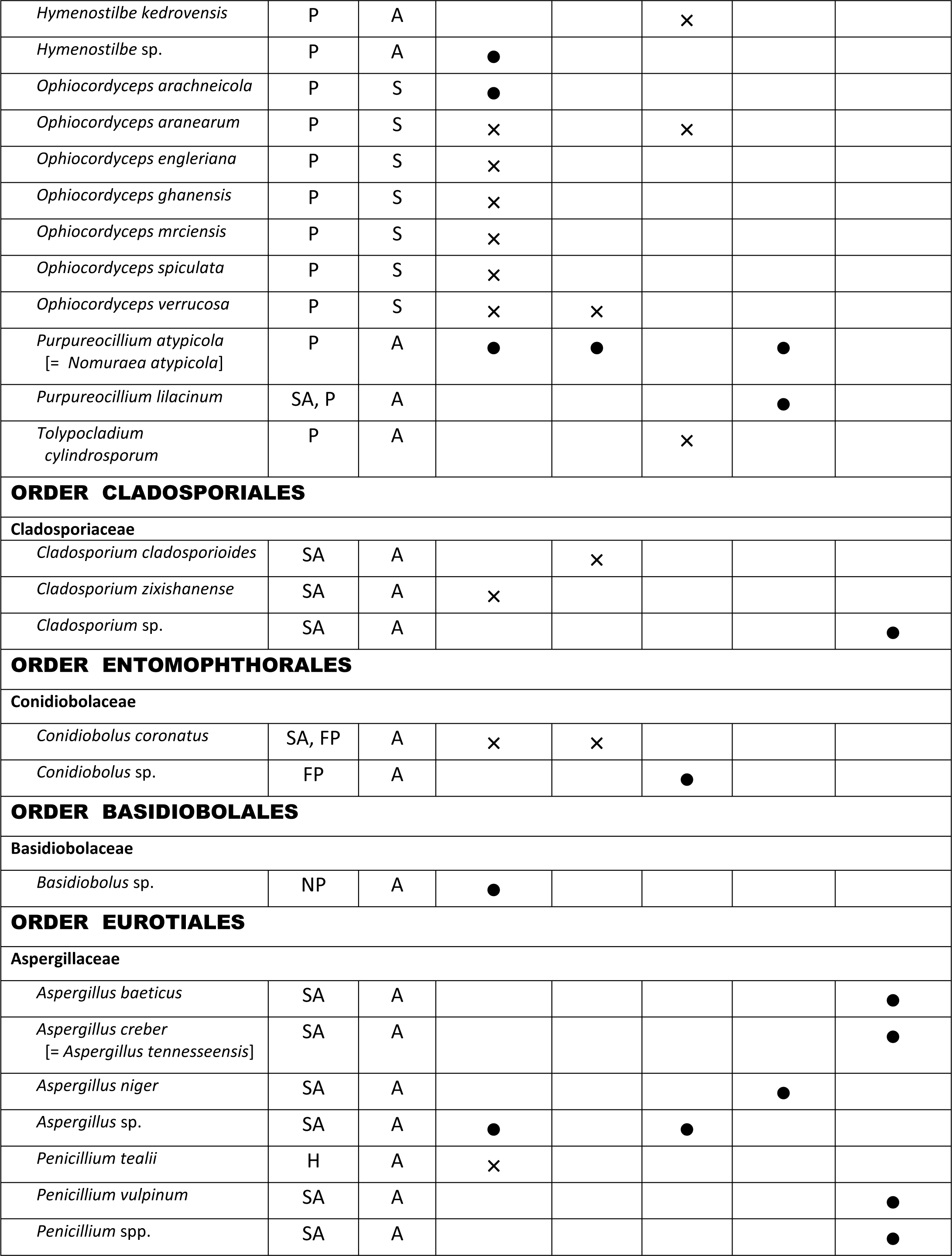
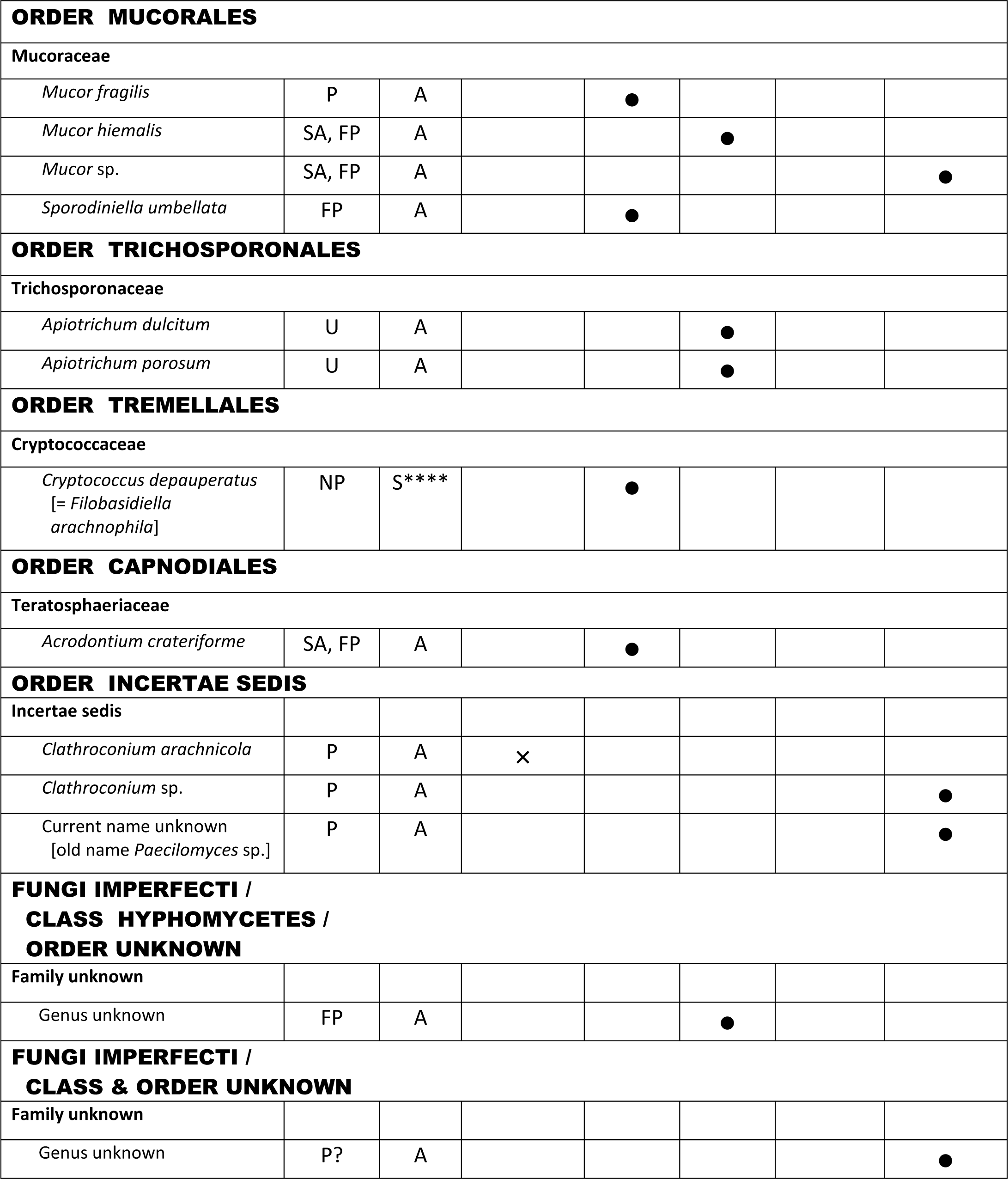
Fungal pathogens infecting different groups of spider hosts (references in Supplemental Materials). Current names based on MycoBank (2023). †Africa, Asia, Central & South America, Oceania. – <u>Type of infection</u>: P = Pathogen; FP = Facultative pathogen; SA = Saprobe; H = Hyperparasite of a *Gibellula* sp.; NP = Non-pathogenic; U = Unknown. – <u>Reproductive mode</u>: A = asexual; S = Sexual. – <u>Host identity</u>: ● = identified host; × = unidentified host. *Asexual morphs [= *Isaria*] found on dead spiders. **Newly discovered fungus with the epithet *Gibellula* ‘*bang-bangus*’ according to CABI (2022); not yet included in the MycoBank data base. ***A species described as “*Verticillium* sp.” (see Austin 1984) was tentatively assigned by us to *Lecanicillium*. ****Sexual morph [= *Filobasidiella arachnophila*] found on a dead spider.

**Appendix 2.**
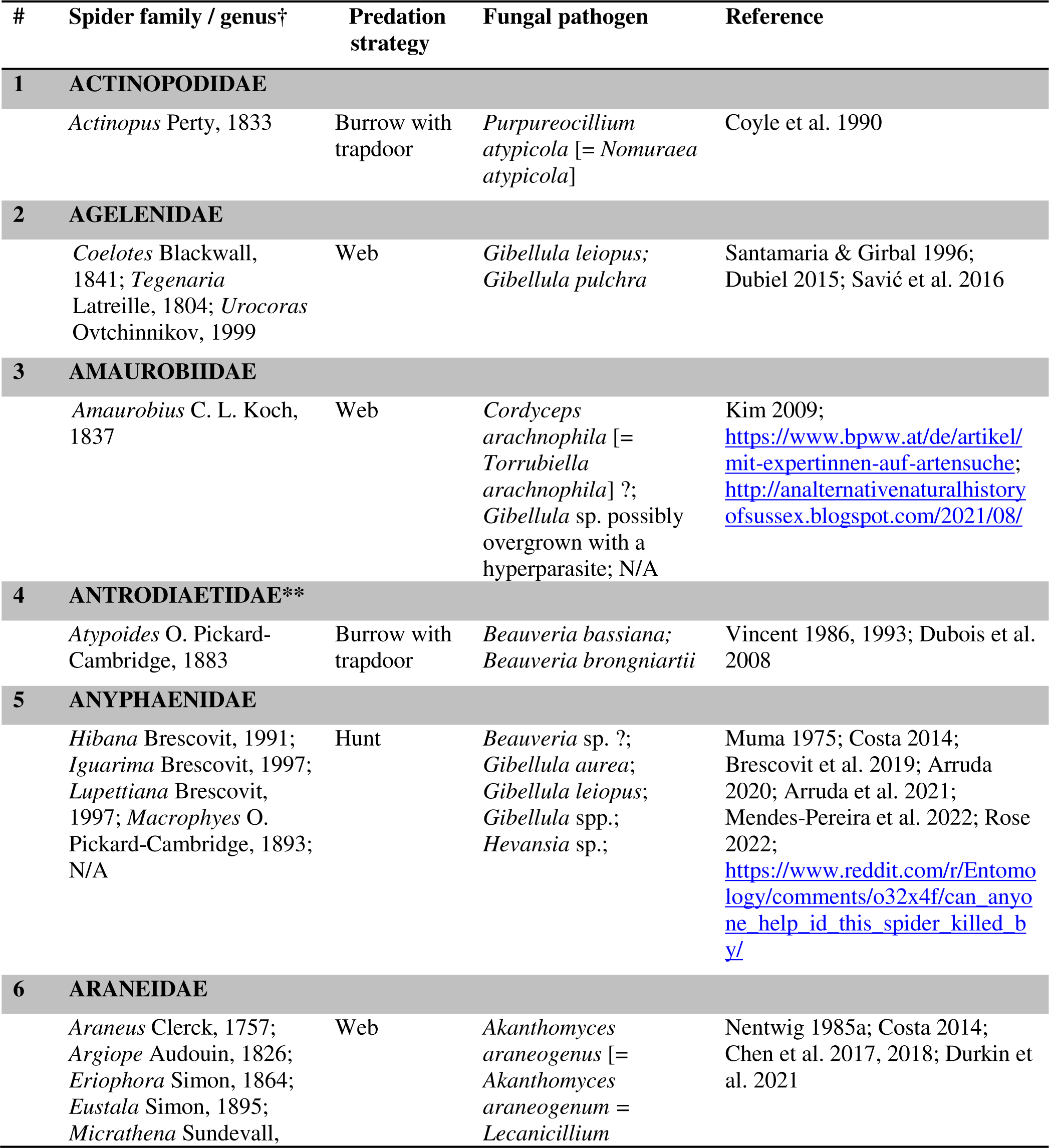
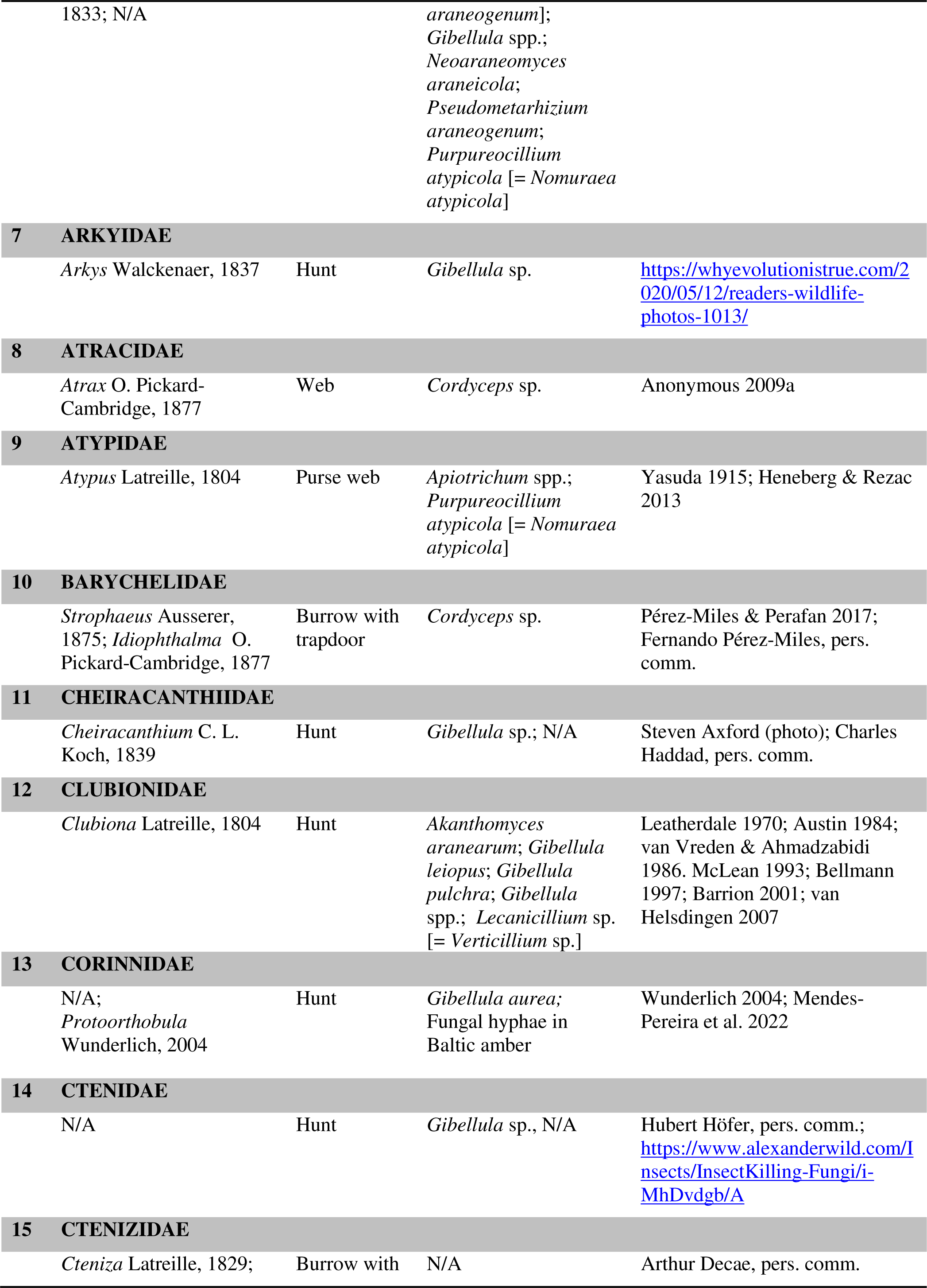
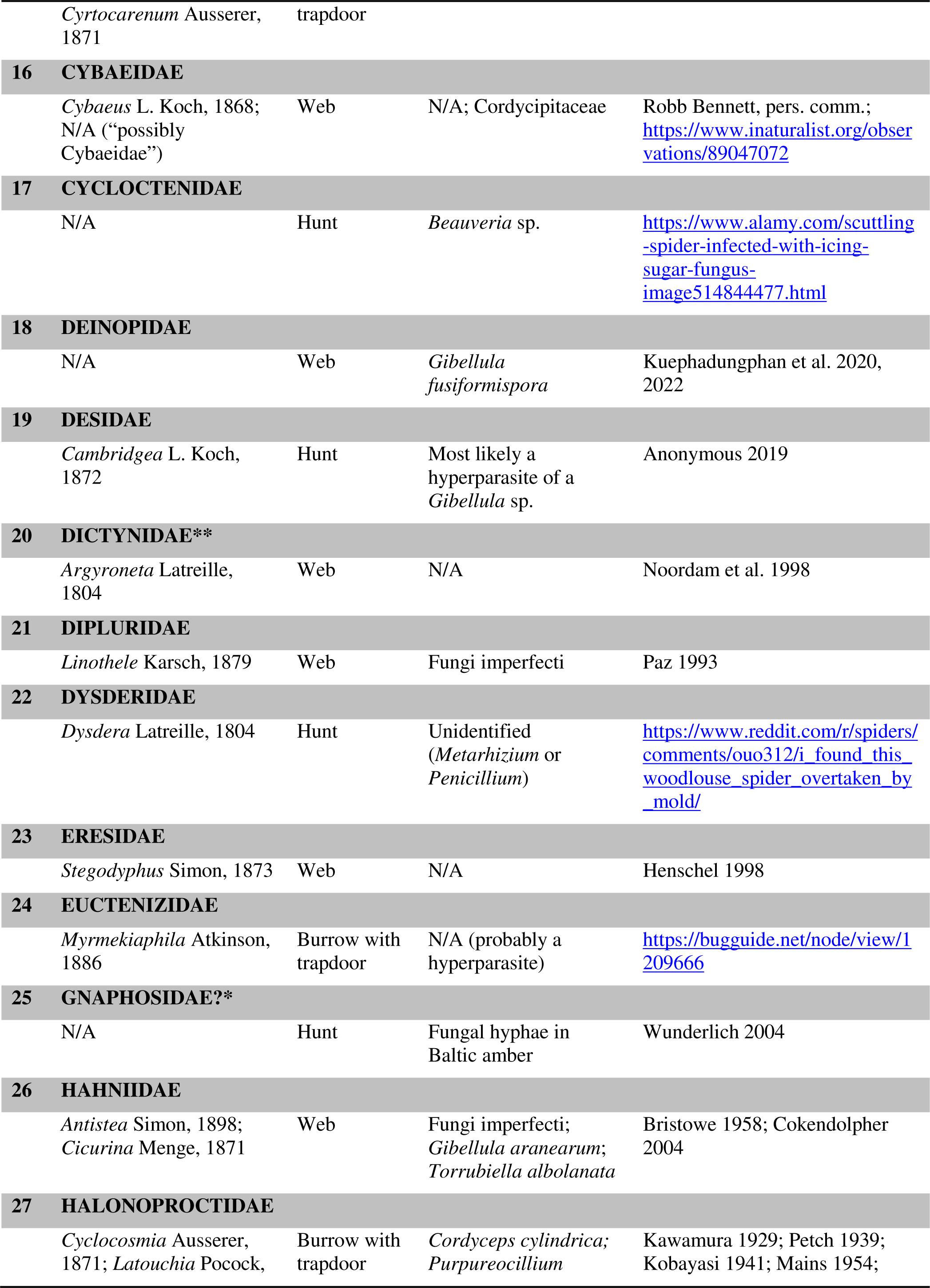
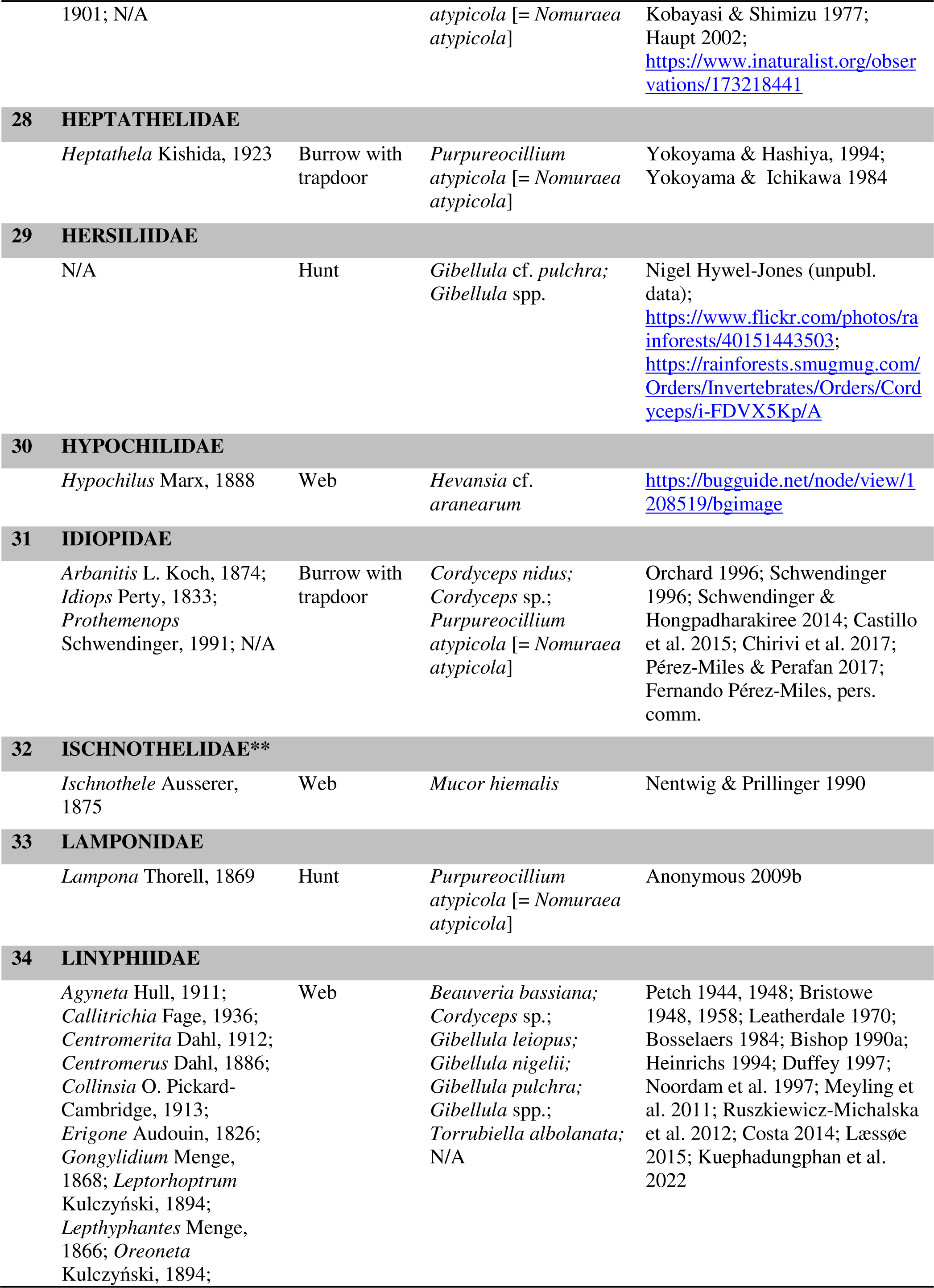
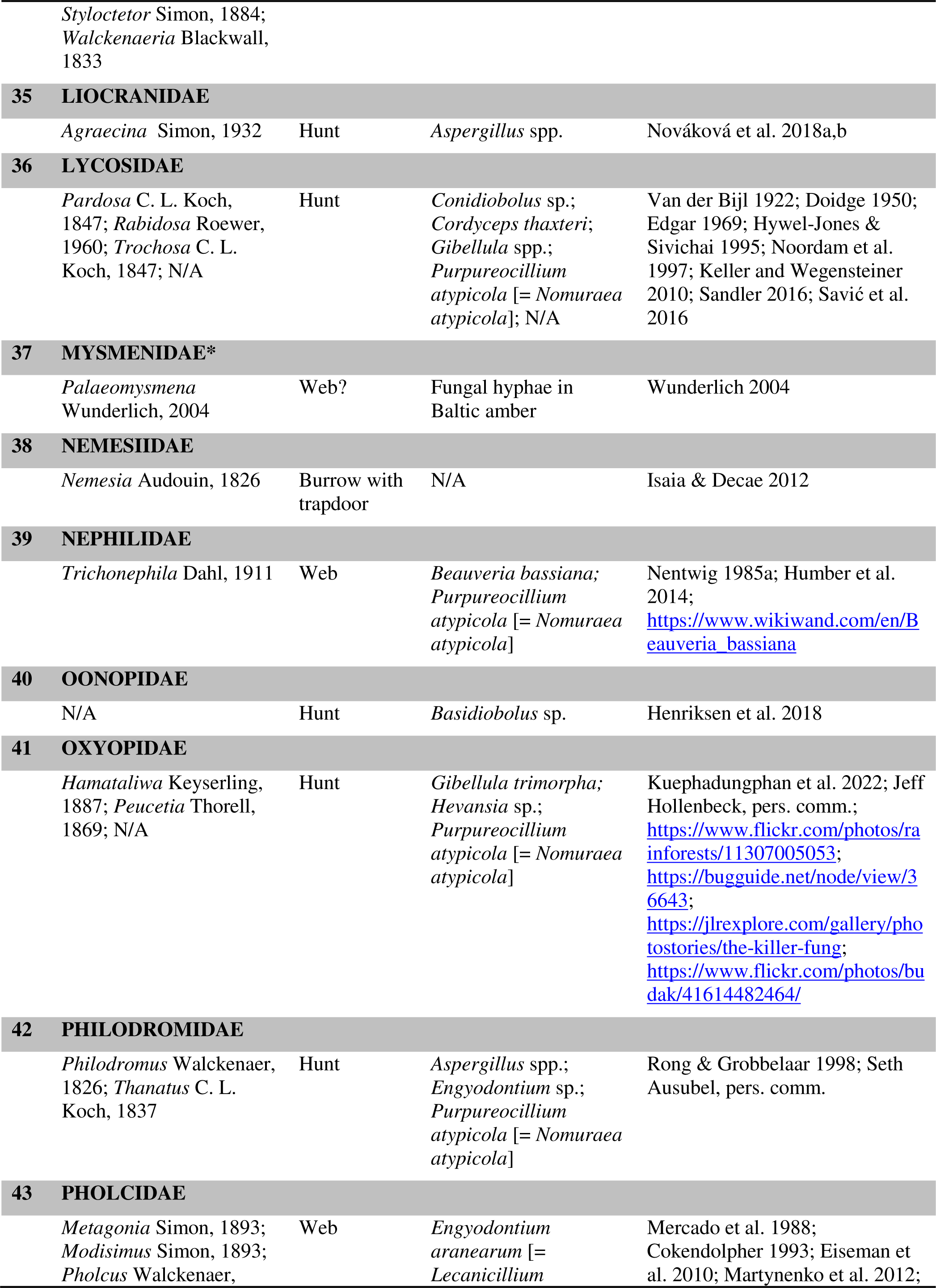
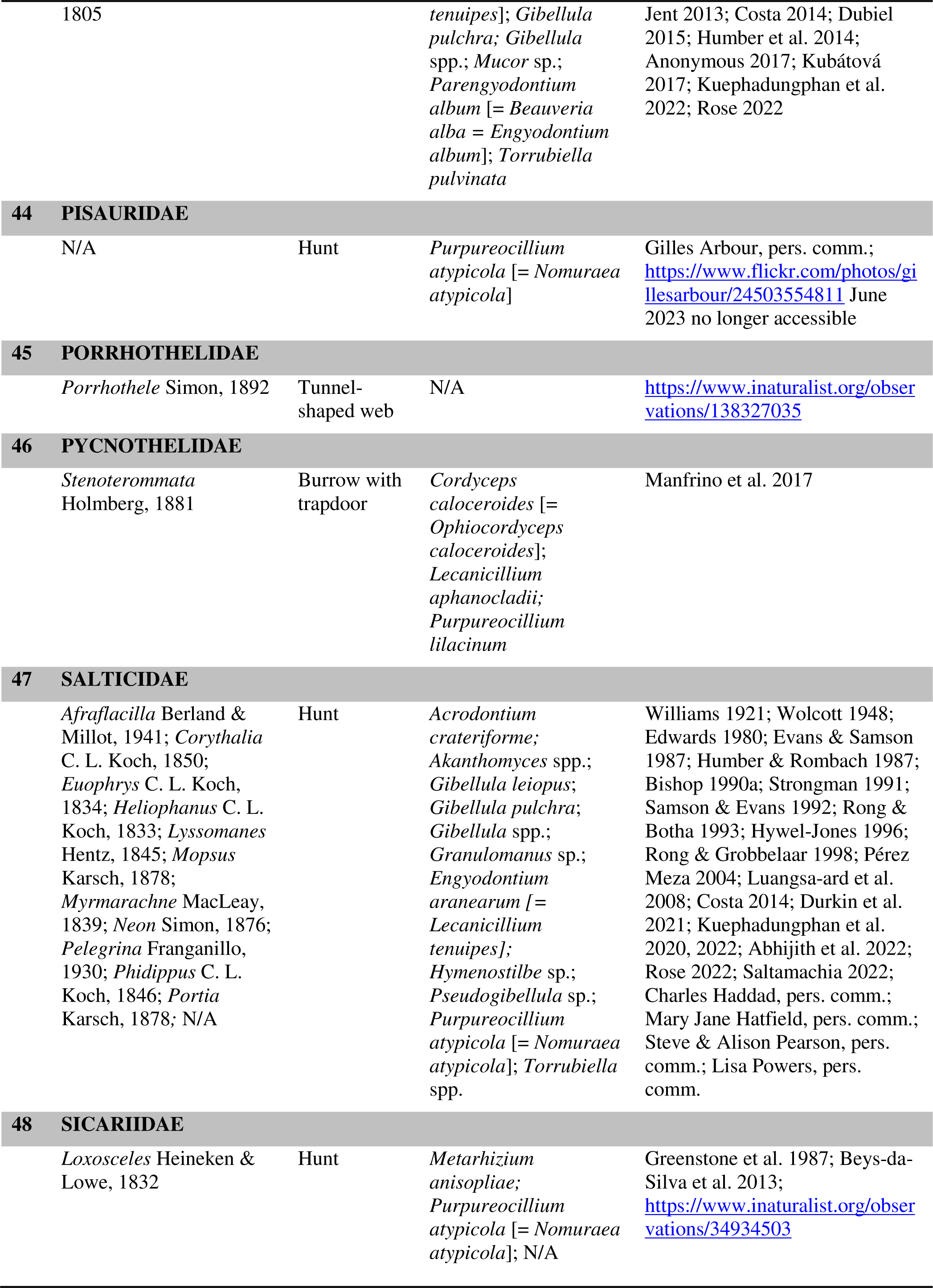
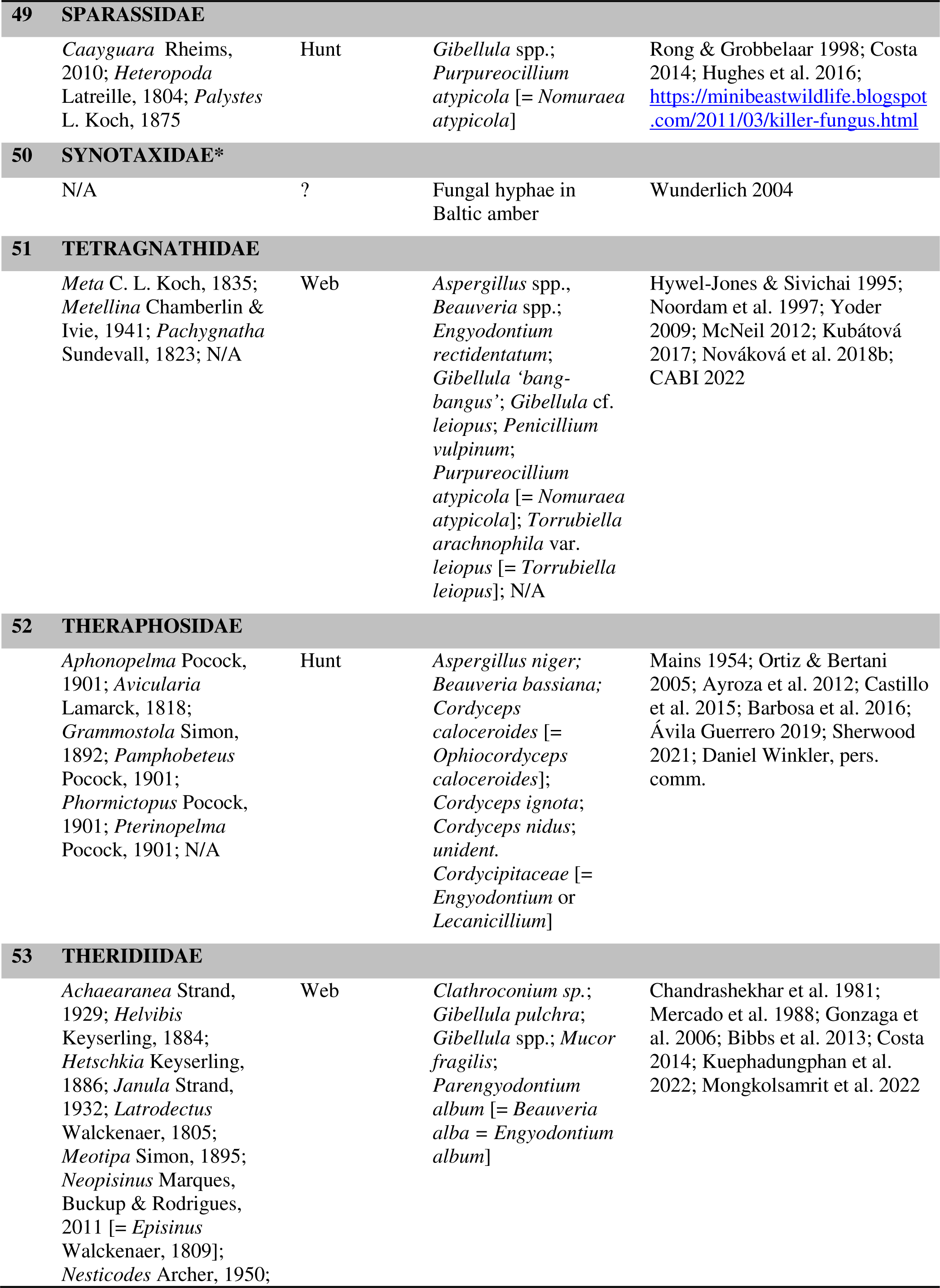
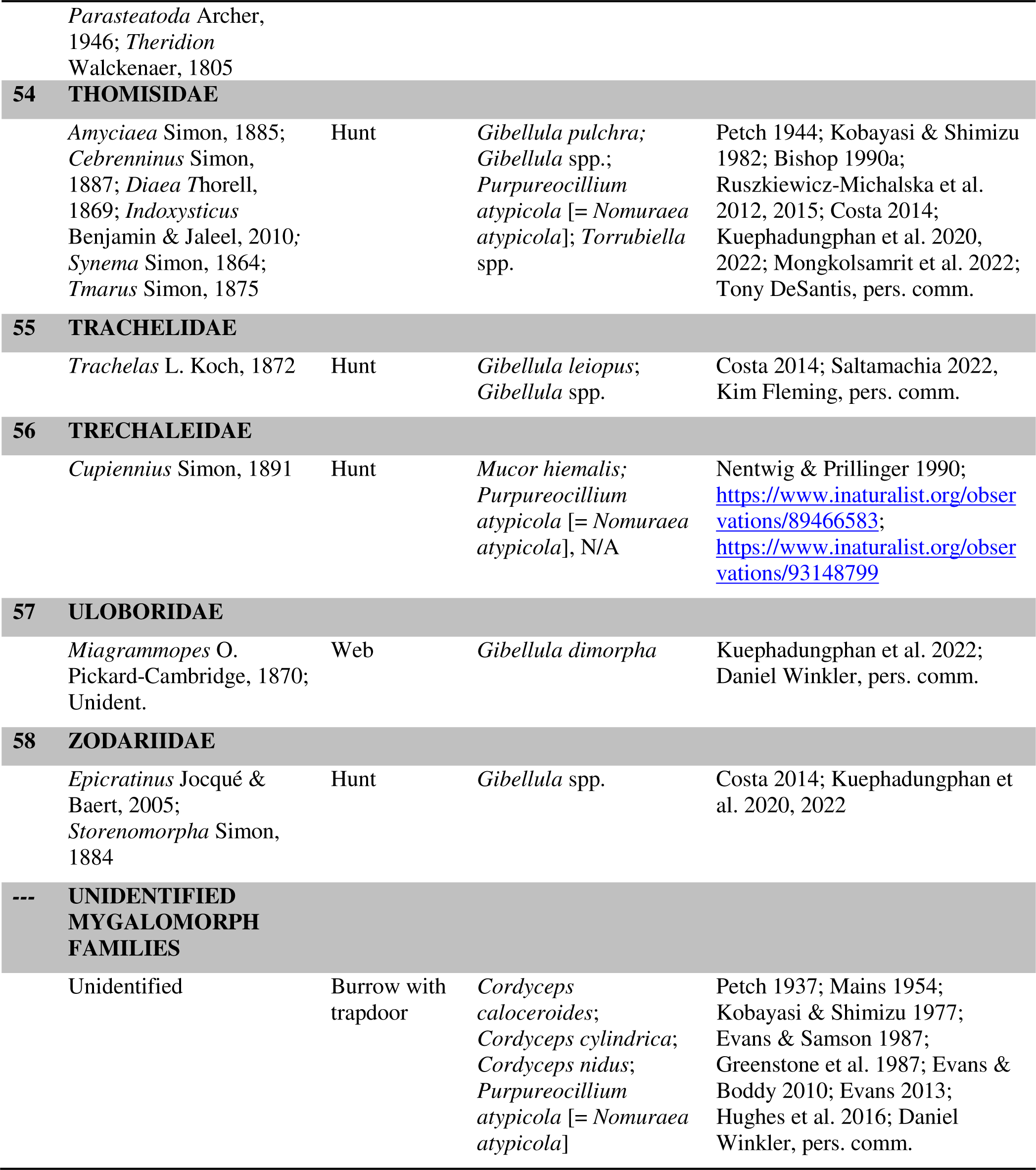
Spiders from various taxa infected by fungal pathogens (based on literature and social media data). †Spider species – as far as identified – have been listed in Supplementary Table S1. *Indicates fungal hyphae on dead spiders encased in samples of Baltic amber. **Indicates fungal infections exclusively under laboratory conditions; all other spider taxa were observed to be infected in the field.

