## Supplementary Table S1 for "Diversity of spider families parasitized by fungal pathogens: a global review"

**Table S1 –** List of spider taxa infected by fungi (based on literature and internet information). *Highly likely a species in the family Cordycipitaceae due to the fact that temperate zone pholcids are infected almost exclusively by the fungi *Engyodontium aranearum* [= *Lecanicillium tenuipes*], *Parengyodontium album* [= *Beauveria alba* = *Engyodontium album*], and *Torrubiella pulvinata*, all of which belong to the Cordycipitaceae (see MycoBank 2023). F = Field observation, L = Laboratory observation.
**Data used to create Fig. 1; the associated references are cited here but not in the Literature Cited section of the paper.

| Spider family / species | Fungus species | Fungus family or higher taxon | Country | Reference |
|---|---|---|---|---|
| **ACTINOPODIDAE** | | | | |
| Actinopus sp. | Purpureocillium atypicola [= Nomuraea atypicola] | Ophiocordycipitaceae | Argentina (F) | 1 |
| **AGELENIDAE** | | | | |
| Agelenopsis sp. | Purpureocillium atypicola [= Nomuraea atypicola] | Ophiocordycipitaceae | USA (L) | 2 |
| Coelotes terrestris (Wider, 1834) | Gibellula leiopus | Cordycipitaceae | Poland (F) | 3 |
| Tegenaria sp. ? | Gibellula pulchra | Cordycipitaceae | Spain (F) | 4 |
| Urocoras longispina (Kulczyński, 1897) | Gibellula leiopus | Cordycipitaceae | Republic of Serbia (F) | 5 |
| **AMAUROBIIDAE** | | | | |
| Amaurobius ferox (Walckenaer, 1830) | Gibellula sp. overgrown with a hyperparasite | Cordycipitaceae | England (F) | 6 |
| Amaurobius ferox (Walckenaer, 1830) | Cordyceps arachnophila [= Torrubiella arachnophila] ? | Cordycipitaceae | Austria (F) | 7 |
| Amaurobius sp. | N/A | N/A | Korea (F) | 8 |
| Amaurobius sp. | N/A | N/A | Korea (L) | 9 |
| **ANTRODIAETIDAE** | | | | |
| Atypoides riversi O. Pickard-Cambridge, 1883 | Beauveria brongniartii | Cordycipitaceae | N/A (L?) | 10 |
| Atypoides riversi O. Pickard-Cambridge, 1883 | Beauveria bassiana | Cordycipitaceae | USA (L) | 11 |
| **ANYPHAENIDAE** | | | | |
| Anyphaena sp. | Purpureocillium atypicola [= Nomuraea atypicola] | Ophiocordycipitaceae | USA (L) | 12 |
| Hibana [= Aysha] gracilis (Hentz, 1847) | Beauveria sp. ? | Cordycipitaceae ? | USA, FL (F) | 13 |
| Iguarima censoria (Keyserling, 1891) | Gibellula sp. | Cordycipitaceae | Brazil (F) | 14 |

| | | | | |
|---|---|---|---|---|
| Iguarima censoria (Keyserling, 1891) | Gibellula sp. | Cordycipitaceae | Brazil (F) | 15 |
| Lupettiana mordax (O. Pickard-Cambridge, 1896) | Hevansia sp. | Cordycipitaceae | USA, Miss (F) | 16 |
| Macrophyes pacoti Brescovit, Oliveira, J. C. M. S. M. Sobczak & J. B. Sobczak, 2019 | Gibellula aurea | Cordycipitaceae | Brazil (F) | 17 |
| Macrophyes pacoti Brescovit, Oliveira, J. C. M. S. M. Sobczak & J. B. Sobczak, 2019 | Gibellula sp. | Cordycipitaceae | Brazil (F) | 18 |
| Macrophyes pacoti Brescovit, Oliveira, J. C. M. S. M. Sobczak & J. B. Sobczak, 2019 | Gibellula sp. | Cordycipitaceae | Brazil (F) | 19 |
| Macrophyes pacoti Brescovit, Oliveira, J. C. M. S. M. Sobczak & J. B. Sobczak, 2019 | Gibellula sp. | Cordycipitaceae | Brazil (F) | 20 |
| N/A | Gibellula leiopus | Cordycipitaceae | Brazil (F) | 21 |
| N/A | Gibellula sp. | Cordycipitaceae | Brazil (F) | 22 |
| N/A | N/A | N/A | USA (F) | 23 |
| **ARANEIDAE** | | | | |
| Acanthepeira stellata (Walckenaer, 1805) | Purpureocillium atypicola [= Nomuraea atypicola] | Ophiocordycipitaceae | USA (L) | 24 |
| Araneus ventricosus (L. Koch, 1878) | Ophiocordyceps arachneicola | Ophiocordycipitaceae | Japan (F) | 25 |
| Araneus sp. | Akanthomyces araneogenus [= Akanthomyces araneogenum = Lecanicillium araneogenum] | Cordycipitaceae | China (F) | 26 |
| Araneus sp. | Akanthomyces araneogenus [= Akanthomyces araneogenum = Lecanicillium araneogenum] | Cordycipitaceae | China (F) | 27 |
| Argiope argentata (Fabricius, 1775) | Purpureocillium atypicola [= Nomuraea atypicola] | Ophiocordycipitaceae | Panama (F) | 28 |
| Argiope aurantia Lucas, 1833 | Purpureocillium atypicola [= Nomuraea atypicola] | Ophiocordycipitaceae | USA (L) | 29 |
| Argiope aurantia Lucas, 1833 | N/A | N/A | USA (F) | 30 |
| Argiope submaronica Strand, 1916 [= Argiope savignyi] | Purpureocillium atypicola [= Nomuraea atypicola] | Ophiocordycipitaceae | Panama (F) | 31 |
| Eriophora fuliginea (C. L. Koch, 1838) | Gibellula sp. | Cordycipitaceae | Panama (F) | 32 |
| Eustala sp. | Gibellula spp. | Cordycipitaceae | Brazil (F) | 33 |
| Micrathena sp. | N/A | Order Hypocreales | Brazil (F) | 34 |
| Neoscona sp. | Purpureocillium atypicola [= Nomuraea atypicola] | Ophiocordycipitaceae | USA (L) | 35 |
| N/A | Gibellula spp. | Cordycipitaceae | Brazil (F) | 36 |
| N/A | Gibellula or Torrubiella | Cordycipitaceae | USA, TN (F) | 37 |
| N/A | Purpureocillium | Ophiocordycipitaceae | Solomon Islands (F) | 38 |

| | | | | |
|---|---|---|---|---|
| | atypicola [= Nomuraea atypicola] | | | |
| **ARKYIDAE** | | | | |
| Arkys lancearius Walckenaer, 1837 | Gibellula sp. | Cordycipitaceae | Australia (F) | 39 |
| **ATRACIDAE** | | | | |
| Atrax robustus O. Pickard-Cambridge, 1877 | Cordyceps sp. | Cordycipitaceae | Australia (F) | 40 |
| **ATYPIDAE** | | | | |
| Atypus affinis Eichwald, 1830 | Apiotrichum porosum | Trichosporonaceae | Czech Republic (F) | 41 |
| Atypus karschi Dönitz, 1887 | Purpureocillium atypicola [= Nomuraea atypicola] | Ophiocordycipitaceae | Japan (F) | 42 |
| Atypus piceus (Sulzer, 1776) | Apiotrichum dulcitum | Trichosporonaceae | Czech Republic (F) | 43 |
| **BARYCHELIDAE** | | | | |
| Idiophthalma sp. | Cordyceps sp. | Cordycipitaceae | Uruguay (F) | 44 |
| Strophaeus sp. | Cordyceps sp. | Cordycipitaceae | Uruguay (F) | 45 |
| **CHEIRACANTHIIDAE** | | | | |
| Cheiracanthium furculatum Karsch, 1879 | N/A | N/A | South Africa (F) | 46 |
| Cheiracanthium gracile L. Koch, 1873 | Gibellula sp. | Cordycipitaceae | Australia (F) | 47 |
| **CLUBIONIDAE** | | | | |
| Clubiona cycladata Simon, 1909 | Lecanicillium sp.? [= In the original paper mentioned as Verticillium sp.] | Cordycipitaceae | Australia (F) | 48 |
| Clubiona robusta L. Koch, 1873 | Lecanicillium sp.? [= In the original paper mentioned as Verticillium sp.] | Cordycipitaceae | Australia (F) | 49 |
| Clubiona terrestris Westring, 1851 | Gibellula pulchra | Cordycipitaceae | Netherland (F) | 50 |
| Clubiona sp. | Akanthomyces aranearum | Cordycipitaceae | England (F) | 51 |
| Clubiona sp. | Gibellula aranearum | Cordycipitaceae | England (F) | 52 |
| Clubiona sp. | Gibellula leiopus | Cordycipitaceae | Peninsular Malaysia (F) | 53 |
| Clubiona sp. | Gibellula leiopus | Cordycipitaceae | Philippines (F) | 54 |
| Clubiona sp. | Gibellula pulchra | Cordycipitaceae | Belgium (F) | 55 |
| N/A | Gibellula leiopus | Cordycipitaceae | Philippines (F) | 56 |
| N/A | N/A | N/A | Germany (F) | 57 |
| **CORINNIDAE** | | | | |
| N/A | Gibellula aurea | Cordycipitaceae | Brazil (F) | 58 |
| Protoorthobula Wunderlich, 2004 | N/A | N/A | Baltic amber (F) | 59 |
| **CTENIDAE** | | | | |
| N/A | Gibellula sp. | Cordycipitaceae | Ecuador (F) | 60 |
| N/A | N/A | Order Hypocreales | Costa Rica (F) | 61 |
| N/A | N/A | Order Hypocreales | Costa Rica (F) | 62 |
| N/A | N/A | N/A | Brazil (F) | 63 |
| **CTENIZIDAE** | | | | |
| Cteniza sp. | N/A | N/A | Southern Europe (F) | 64 |
| Cyrtocarenum cunicularium (Olivier, 1811) | N/A | N/A | Greece (F) | 65 |

| | | | | |
|---|---|---|---|---|
| **CYBAEIDAE** | | | | |
| N/A | N/A | Cordycipitaceae | USA (F) | 66 |
| Cybaeus reticulatus Simon, 1886 | N/A | N/A | Canada (F) | 67 |
| **CYCLOCTENIDAE** | | | | |
| N/A | Beauveria sp. | Cordycipitaceae | New Zealand (F) | 68 |
| **DEINOPIDAE** | | | | |
| N/A | Gibellula fusiformispora | Cordycipitaceae | Thailand (F) | 69 |
| **DESIDAE** | | | | |
| Cambridgea sp. | Hyperparasite of a Gibellula | Cordycipitaceae + N/A | New Zealand (F) | 70 |
| **DICTYNIDAE** | | | | |
| Argyroneta aquatica (Clerck, 1757) | N/A | N/A | Germany (L) | 71 |
| **DIPLURIDAE** | | | | |
| Linothele megatheloides Paz & Raven, 1990 | N/A | Fungi imperfecti | Colombia (F) | 72 |
| **DYSDERIDAE** | | | | |
| Dysdera sp. | N/A | Unidentified (Metarhizium or Penicillium) | USA? (F) | 73 |
| **ERESIDAE** | | | | |
| Stegodyphus dumicola Pocock, 1898 | N/A | N/A | Africa (F) | 74 |
| **EUCTENIZIDAE** | | | | |
| Myrmekiaphila sp. | N/A | Probably a fungus hyperparasite overgrowing an unknown fungus | USA (F) | 75 |
| **GNAPHOSIDAE?** | | | | |
| N/A | N/A | N/A | Baltic amber (F) | 76 |
| **HAHNIIDAE** | | | | |
| Antistea elegans (Blackwall, 1841) | Torrubiella albolanata | Cordycipitaceae | England (F) | 77 |
| Antistea elegans (Blackwall, 1841) | Gibellula aranearum | Cordycipitaceae | England (F) | 78 |
| Cicurina sp. | N/A | Fungi Imperfecti / Class & Order unknown / Family unknown | USA (F) | 79 |
| Eocryphoeca ligula Wunderlich, 2004 | N/A | N/A | Baltic amber (F) | 80 |
| **HALONOPROCTIDAE** | | | | |
| Cyclocosmia truncata (Hentz, 1841) | Purpureocillium atypicola [= Nomuraea atypicola] | Ophiocordycipitaceae | USA (F) | 81 |
| Latouchia hyla Haupt & Shimojana, 2001 | Purpureocillium atypicola [= Nomuraea atypicola] | Ophiocordycipitaceae | Japan (F) | 82 |
| Latouchia japonica Strand, 1910 | Purpureocillium atypicola [= Nomuraea atypicola] | Ophiocordycipitaceae | Japan (F) | 83 |
| Latouchia typica (Kishida, 1913) | Purpureocillium atypicola [= Nomuraea | Ophiocordycipitaceae | Japan (F) | 84 |

| | | | | |
|---|---|---|---|---|
| | atypicola] | | | |
| Latouchia typica (Kishida, 1913) [= Latouchia swinhoeitypica] | Cordyceps cylindrica | Cordycipitaceae | Japan (F) | 85 |
| Latouchia typica (Kishida, 1913) [= Latouchia swinhoeitypica] | Purpureocillium atypicola [= Nomuraea atypicola] | Ophiocordycipitaceae | Japan (F) | 86 |
| Latouchia sp. | Purpureocillium atypicola [= Nomuraea atypicola] | Ophiocordycipitaceae | Japan (F) | 87 |
| N/A | Cordyceps cylindrica | Cordycipitaceae | Caribic / Trinidad (F) | 88 |
| N/A | Purpureocillium atypicola [= Nomuraea atypicola] | Ophiocordycipitaceae | Japan (F) | 89 |
| **HEPTATHELIDAE** | | | | |
| Heptathela kimurai (Kishida, 1920) | Purpureocillium atypicola [= Nomuraea atypicola] | Ophiocordycipitaceae | Japan (F) | 90 |
| **HERSILIIDAE** | | | | |
| N/A | Gibellula cf. pulchra | Cordycipitaceae | Vietnam (F) | 91 |
| N/A | Gibellula sp. | Cordycipitaceae | Peru (F) | 92 |
| N/A | Gibellula sp. | Cordycipitaceae | Thailand (F) | 93 |
| **HYPOCHILIDAE** | | | | |
| Hypochilus pococki Platnick, 1987 | Hevansia cf. aranearum | Cordycipitaceae | USA (F) | 94 |
| **IDIOPIDAE** | | | | |
| Arbanitis rapax (Karsch, 1878) | Purpureocillium atypicola [= Nomuraea atypicola] | Ophiocordycipitaceae | Australia (F) | 95 |
| Idiops sp. | Cordyceps sp. | Cordycipitaceae | Uruguay (F) | 96 |
| Prothemenops irineae Schwendinger & Hongpadharakiree, 2014 | Purpureocillium atypicola [= Nomuraea atypicola] | Ophiocordycipitaceae | Thailand (L) | 97 |
| Prothemenops siamensis Schwendinger, 1991 | Purpureocillium atypicola [= Nomuraea atypicola] | Ophiocordycipitaceae | Thailand (F) | 98 |
| N/A | Cordyceps nidus | Cordycipitaceae | Colombia (F) | 99 |
| N/A | Cordyceps nidus | Cordycipitaceae | Colombia (F) | 100 |
| **ISCHNOTHELIDAE** | | | | |
| Ischnothele guianensis (Walckenaer, 1837) | Mucor hiemalis | Mucoraceae | Germany (L) | 101 |
| **LAMPONIDAE** | | | | |
| Lampona sp. | Purpureocillium atypicola [= Nomuraea atypicola] | Ophiocordycipitaceae | New Zealand (F) | 102 |
| **LINYPHIIDAE** | | | | |
| Agyneta nigripes (Simon, 1884) | Cordyceps sp. | Cordycipitaceae | Arctic Island, Norway (F) | 103 |
| Atypena formosana (Oi, 1977) [= Callitrichia formosana] | Gibellula leiopus | Cordycipitaceae | Philippines (F) | 104 |
| Atypena sp. [= Callitrichia sp.] | Gibellula leiopus | Cordycipitaceae | Philippines (F) | 105 |
| Centromerita concinna (Thorell, 1875) | N/A | Class Hyphomycetes / unknown family | Netherlands (F) | 106 |
| Centromerus prudens (O. Pickard-Cambridge, 1873) | N/A | Class Hyphomycetes / unknown family | Netherlands (F) | 107 |
| Centromerus sylvaticus | N/A | Class Hyphomycetes / | Netherlands (F) | 108 |

| | | | | |
|---|---|---|---|---|
| (Blackwall, 1841) | | unknown family | | |
| Collinsia holmgreni (Thorell, 1871) | Cordyceps sp. | Cordycipitaceae | Arctic Island, Norway (F) | 109 |
| Erigone tirolensis L. Koch, 1872 | Cordyceps sp. | Cordycipitaceae | Arctic Island, Norway (F) | 110 |
| Linyphiidae-Erigoninae | Torrubiella albolanata | Cordycipitaceae | Denmark (F) Jutland | 111 |
| Frontinella pyramitela (Walckenaer, 1841) | Purpureocillium atypicola [= Nomuraea atypicola] | Ophiocordycipitaceae | USA (L) | 112 |
| Gongylidium rufipes (Linnaeus, 1758) | Gibellula pulchra | Cordycipitaceae | England (F) | 113 |
| Gongylidium rufipes (Linnaeus, 1758) | Torrubiella albolanata | Cordycipitaceae | England (F) | 114 |
| Leptorhoptrum robustum (Westring, 1851) | Gibellula aranearum | Cordycipitaceae | England (F) | 115 |
| Leptorhoptrum robustum (Westring, 1851) | Torrubiella albolanata | Cordycipitaceae | England (F) | 116 |
| Palliduphantes pallidus (O. Pickard-Cambridge, 1871) [= Lepthyphantes pallidus] | N/A | Class Hyphomycetes / unknown family | Netherlands (F) | 117 |
| Linyphiidae-Linyphiinae | Gibellula or Torrubiella | Cordycipitaceae | USA, TN (F) | 118 |
| Linyphiidae-Erigoninae | Gibellula or Torrubiella | Cordycipitaceae | USA, TN (F) | 119 |
| Oreoneta frigida (Thorell, 1872) | Cordyceps sp. | Cordycipitaceae | Arctic Island, Norway (F) | 120 |
| Styloctetor romanus (O. Pickard-Cambridge, 1873) [= Ceratinopsis romana] | N/A | Class Hyphomycetes / unknown family | Netherlands (F) | 121 |
| Walckenaeria antica (Wider, 1834) | N/A | Class Hyphomycetes / unknown family | Netherlands (F) | 122 |
| Walckenaeria monoceros (Wider, 1834) | N/A | Class Hyphomycetes / unknown family | Netherlands (F) | 123 |
| N/A – study year 1 | Beauveria bassiana | Cordycipitaceae | Denmark (F) | 124 |
| N/A | Gibellula aranearum | Cordycipitaceae | England (F) | 125 |
| N/A – record 1 | Gibellula aranearum | Cordycipitaceae | England (F) | 126 |
| N/A – record 2 | Gibellula aranearum | Cordycipitaceae | England (F) | 127 |
| N/A | Gibellula leiopus | Cordycipitaceae | Poland (F) | 128 |
| N/A | Gibellula nigelii | Cordycipitaceae | Thailand (F) | 129 |
| N/A | Gibellula pulchra | Cordycipitaceae | Belgium (F) | 130 |
| N/A – study year 1 | Gibellula spp. | Cordycipitaceae | Denmark (F) | 131 |
| N/A – study year 2 | Gibellula spp. | Cordycipitaceae | Denmark (F) | 132 |
| N/A | Gibellula spp. | Cordycipitaceae | Brazil (F) | 133 |
| N/A | Torrubiella albolanata | Cordycipitaceae | Denmark (F) Bog | 134 |
| N/A | Torrubiella albolanata | Cordycipitaceae | England (F) | 135 |
| **LIOCRANIDAE** | | | | |
| Agraecina cristiani (Georgescu, 1989) | Aspergillus baeticus | Aspergillaceae | Romania (F) | 136 |
| **LYCOSIDAE** | | | | |
| Rabidosa rabida (Walckenaer, 1837) [= Lycosa rabida] | Purpureocillium atypicola [= Nomuraea atypicola] | Ophiocordycipitaceae | USA (L) | 137 |
| Pardosa amentata (Clerck, 1757) | Conidiobolus sp. | Conidiobolaceae | Switzerland (F) | 138 |
| Pardosa lugubris (Walckenaer, 1802) | N/A | N/A | Scotland (F) | 139 |
| Trochosa terricola Thorell, 1856 | N/A | Class Hyphomycetes / unknown family | Netherlands (F) | 140 |

| | | | | |
|---|---|---|---|---|
| N/A | Cordyceps thaxteri | Cordycipitaceae | Republic of Serbia (F) | 141 |
| N/A | Gibellula aranearum | Cordycipitaceae | South Africa (F) | 142 |
| N/A | Gibellula sp. | Cordycipitaceae | South Africa (F) | 143 |
| N/A | Gibellula sp. | Cordycipitaceae | USA (F) | 144 |
| N/A | Purpureocillium atypicola [= Nomuraea atypicola] | Ophiocordycipitaceae | Thailand (F) | 145 |
| N/A | N/A | Order Hypocreales / unknown family | Costa Rica (F) | 146 |
| **MYSMENIDAE** | | | | |
| Palaeomysmena hoffeinsorum Wunderlich, 2004 | N/A | N/A | Baltic amber (F) | 147 |
| **NEMESIIDAE** | | | | |
| Nemesia meridionalis (Costa, 1835) | N/A | N/A | Italy (F) | 148 |
| **NEPHILIDAE** | | | | |
| Trichonephila clavipes (Linnaeus, 1767) | Beauveria bassiana | Cordycipitaceae | USA (F) | 149 |
| Trichonephila clavipes (Linnaeus, 1767) | Purpureocillium atypicola [= Nomuraea atypicola] | Ophiocordycipitaceae | Panama (F) | 150 |
| Trichonephila clavipes (Linnaeus, 1767) | Purpureocillium atypicola [= Nomuraea atypicola] | Ophiocordycipitaceae | USA (F) | 151 |
| Trichonephila clavipes (Linnaeus, 1767) | Sporodiniella umbellata | Mucoraceae | USA (F) | 152 |
| Trichonephila clavipes (Linnaeus, 1767) | N/A | N/A | South America? (F) | 153 |
| Trichonephila clavipes (Linnaeus, 1767) | N/A | N/A | USA (F) | 154 |
| **OONOPIDAE** | | | | |
| N/A | Basidiobolus sp. | Basidiobolaceae | Tanzania (F) | 155 |
| **OXYOPIDAE** | | | | |
| Hamataliwa sp. | Hevansia sp. | Cordycipitaceae | Cambodia (F) | 156 |
| Peucetia viridans (Hentz, 1832) | Purpureocillium atypicola [= Nomuraea atypicola] | Ophiocordycipitaceae | USA, FL (F) | 157 |
| N/A | Unidentified (Engyodontium or Lecanicillium) | Cordycipitaceae | India (F) | 158 |
| N/A | Gibellula trimorpha | Cordycipitaceae | Thailand (F) | 159 |
| N/A | N/A | N/A | Singapore (F) | 160 |
| **PHILODROMIDAE** | | | | |
| Philodromus sp. | possibly Purpureocillium atypicola [= Nomuraea atypicola] | Ophiocordycipitaceae | USA (F) | 161 |
| Thanatus sp. | Aspergillus sp. | Aspergillaceae | South Africa (F) | 162 |
| Thanatus sp. | Engyodontium sp. | Cordycipitaceae | South Africa (F) | 163 |
| **PHOLCIDAE** | | | | |
| Metagonia taruma Huber, 2000 | Gibellula sp. | Cordycipitaceae | Brazil (F) | 164 |
| Metagonia sp. | Gibellula pulchra | Cordycipitaceae | Brazil (F) | 165 |
| Modisimus sp. | Mucor sp. | Mucoraceae | Cuba (F) | 166 |

| | | | | |
|---|---|---|---|---|
| Pholcus phalangioides (Fuesslin, 1775) | Engyodontium aranearum [= Lecanicillium tenuipes] | Cordycipitaceae | USA (F) | 167 |
| Pholcus phalangioides (Fuesslin, 1775) | Engyodontium aranearum [= Lecanicillium tenuipes] | Cordycipitaceae | USA (L) | 168 |
| Pholcus phalangioides (Fuesslin, 1775) | N/A | Cordycipitaceae* | UK (F) | 169 |
| Pholcus phalangioides (Fuesslin, 1775) | N/A | Cordycipitaceae* | Italy (F) | 170 |
| Pholcus phalangioides (Fuesslin, 1775) | N/A | Cordycipitaceae* | Germany (F) | 171 |
| Pholcus reevesi Huber, 2011 | N/A | Cordycipitaceae* | USA (F) | 172 |
| Pholcus sp. | Parengyodontium album [= Beauveria alba = Engyodontium album] | Cordycipitaceae | Ukraine (F) | 173 |
| Pholcus sp. | Engyodontium aranearum [= Lecanicillium tenuipes] | Cordycipitaceae | Netherlands (F) | 174 |
| Pholcus sp. | Engyodontium aranearum [= Lecanicillium tenuipes] | Cordycipitaceae | Poland (F) | 175 |
| Pholcus sp. | Torrubiella pulvinata | Cordycipitaceae | USA (F) | 176 |
| Pholcus sp. | N/A | Cordycipitaceae* | Poland (F) | 177 |
| Pholcus sp. | N/A | Cordycipitaceae* | Spain (F) | 178 |
| Pholcus sp. | N/A | Cordycipitaceae* | Island of Corse, France (F) | 179 |
| Pholcus sp. | N/A | Cordycipitaceae* | Denmark (F) | 180 |
| Pholcus sp. | N/A | Cordycipitaceae* | Italy (F9 | 181 |
| Pholcus sp. | N/A | Cordycipitaceae* | Slovakia (F) | 182 |
| Pholcus sp. | N/A | Cordycipitaceae* | Ukraine / Europe (F) | 183 |
| Pholcus sp. | N/A | Cordycipitaceae* | Russia / Europe (F) | 184 |
| Pholcus sp. | N/A | Cordycipitaceae* | Denmark (F) | 185 |
| Pholcus sp. | N/A | Cordycipitaceae* | Portugal (F) | 186 |
| Pholcus sp. | N/A | Cordycipitaceae* | Russia / Europe (F) | 187 |
| Pholcus sp. | N/A | Cordycipitaceae* | Russia / Europe (F) | 188 |
| Pholcus sp. | N/A | Cordycipitaceae* | Russia / Europe (F) | 189 |
| Pholcus sp. | N/A | Cordycipitaceae* | Spain (F) | 190 |
| Pholcus sp. | N/A | Cordycipitaceae* | Belgium (F) | 191 |
| Pholcus sp. | N/A | Cordycipitaceae* | Denmark (F) | 192 |
| Pholcus sp. | N/A | Cordycipitaceae* | Slovenija (F) | 193 |
| Pholcus sp. | N/A | Cordycipitaceae* | Lithuania / Europe (F) | 194 |
| Pholcus sp. | N/A | Cordycipitaceae* | Italy (F) | 195 |
| Pholcus sp. | N/A | Cordycipitaceae* | Russia / Europe (F) | 196 |
| Pholcus sp. | N/A | Cordycipitaceae* | Latvia / Europe (F) | 197 |
| Pholcus sp. | N/A | Cordycipitaceae* | Lithuania / Europe (F) | 198 |
| Pholcus sp. | N/A | Cordycipitaceae* | USA (F) | 199 |
| Pholcus sp. | N/A | Cordycipitaceae* | USA (F) | 200 |
| Pholcus sp. | N/A | Cordycipitaceae* | USA (F) | 201 |
| Pholcus sp. | N/A | Cordycipitaceae* | Canada (F) | 202 |
| Pholcus sp. | N/A | Cordycipitaceae* | USA (F) | 203 |
| Pholcus sp. | N/A | Cordycipitaceae* | USA (F) | 204 |
| Pholcus sp. | N/A | Cordycipitaceae* | Canada (F) | 205 |
| Pholcus sp. | N/A | Cordycipitaceae* | Canada (F) | 206 |
| Pholcus sp. | N/A | Cordycipitaceae* | Canada (F) | 207 |

| | | | | |
|---|---|---|---|---|
| Pholcus sp. | N/A | Cordycipitaceae* | USA (F) | 208 |
| Pholcus sp. | N/A | Cordycipitaceae* | Canada (F) | 209 |
| Pholcus sp. | N/A | Cordycipitaceae* | USA (F) | 210 |
| Pholcus sp. | N/A | Cordycipitaceae* | USA (F) | 211 |
| Pholcus sp. | N/A | Cordycipitaceae* | USA (F) | 212 |
| Pholcus sp. | N/A | Cordycipitaceae* | USA (F) | 213 |
| Pholcus sp. | N/A | Cordycipitaceae* | USA (F) | 214 |
| Pholcus sp. | N/A | Cordycipitaceae* | Hungary (F) | 215 |
| N/A | Parengyodontium album [= Beauveria alba = Engyodontium album] | Cordycipitaceae | Solomon Islands (F) | 216 |
| N/A | Gibellula unica | Cordycipitaceae | Thailand (F) | 217 |
| N/A | Engyodontium aranearum [= Lecanicillium tenuipes] | Cordycipitaceae | Hungary (F) | 218 |
| N/A | Engyodontium aranearum [= Lecanicillium tenuipes] | Cordycipitaceae | USA (F) | 219 |
| N/A | Engyodontium aranearum [= Lecanicillium tenuipes] | Cordycipitaceae | USA (F) | 220 |
| N/A | Torrubiella pulvinata | Cordycipitaceae | USA, Hawaii (F) | 221 |
| N/A | N/A | N/A | USA (F) | 222 |
| N/A | Engyodontium aranearum [= Lecanicillium tenuipes] | Cordycipitaceae | Czech Republik (F) | 223 |
| N/A | Engyodontium aranearum [= Lecanicillium tenuipes] | Cordycipitaceae | Denmark (F) | 224 |
| N/A | Engyodontium aranearum [= Lecanicillium tenuipes] | Cordycipitaceae | France (F) | 225 |
| N/A | Engyodontium aranearum [= Lecanicillium tenuipes] | Cordycipitaceae | Poland (F) | 226 |
| N/A | Engyodontium aranearum [= Lecanicillium tenuipes] | Cordycipitaceae | England (F) | 227 |
| **PISAURIDAE** | | | | |
| N/A | Purpureocillium atypicola [= Nomuraea atypicola] | Ophiocordycipitaceae | Ecuador (F) | 228 |
| **PORRHOTHELIDAE** | | | | |
| Porrhothele antipodiana (Walckenaer, 1837) | N/A | N/A | New Zealand (F) | 229 |
| **PYCNOTHELIDAE** | | | | |
| Stenoterommata platensis Holmberg, 1881 | Lecanicillium aphanocladii | Cordycipitaceae | Argentina (F) | 230 |
| Stenoterommata platensis Holmberg, 1881 | Cordyceps caloceroides [= Ophiocordyceps caloceroides] | Ophiocordycipitaceae | Argentina (F) | 231 |
| Stenoterommata platensis Holmberg, 1881 | Purpureocillium lilacinum | Ophiocordycipitaceae | Argentina (F) | 232 |
| **SALTICIDAE** | | | | |

| | | | | |
|---|---|---|---|---|
| Afraflacilla venustula (Wesołowska & Haddad, 2009) | N/A | N/A | South Africa (F) | 233 |
| Anasaitis banksi (Roewer, 1951) [= Prostheclina signata Banks, 1901] | Gibellula arachnophila | Cordycipitaceae | Puerto Rico (F) | 234 |
| Anasaitis canosa (Walckenaer, 1837) | Gibellula pulchra | Cordycipitaceae | USA, LA (F) | 235 |
| Colonus sylvanus (Hentz, 1846) | N/A | N/A | USA (F) | 236 |
| Colonus sp. | Gibellula pulchra | Cordycipitaceae | USA, LA (F) | 237 |
| Colonus sp. | Gibellula cf. leiopus | Cordycipitaceae | USA (F) | 238 |
| Corythalia sp. | Gibellula spp. | Cordycipitaceae | Brazil (F) | 239 |
| Euophrys sp. | Torrubiella ratticaudata | Cordycipitaceae | Solomon Islands (F) | 240 |
| Euophrys sp. | Gibellula clavulifera var. alba | Cordycipitaceae | Solomon Islands (F) | 241 |
| Heliophanus pistaciae Wesołowska, 2003 | N/A | N/A | South Africa (F) | 242 |
| Hentzia palmarum (Hentz, 1832) [= Hentzia ambiguus] | Purpureocillium atypicola [= Nomuraea atypicola] | Ophiocordycipitaceae | USA (L) | 243 |
| Lyssomanes viridis (Walckenaer, 1837) | Gibellula sp. | Cordycipitaceae | USA (F) | 244 |
| Pelegrina galathea (Walckenaer, 1837) [= Metaphidippus galathea] | Purpureocillium atypicola [= Nomuraea atypicola] | Ophiocordycipitaceae | USA (L) | 245 |
| Pelegrina proterva (Walckenaer, 1837) | Gibellula cf. leiopus | Cordycipitaceae | USA (F) | 246 |
| Mopsus mormon Karsch, 1878 | Cordyceps sp. | Cordycipitaceae | Australia (F) | 247 |
| Myrmaplata plataleoides (O. Pickard-Cambridge, 1869) [= Myrmarachne plataleoides] | N/A | N/A | India (F) | 248 |
| Myrmarachne sp. | Gibellula longispora | Cordycipitaceae | Thailand (F) | 249 |
| Neon nelli G. W. Peckham & E. G. Peckham, 1888 | Gibellula pulchra | Cordycipitaceae | Canada (F) | 250 |
| Phidippus audax (Hentz, 1845) | Gibellula leiopus | Cordycipitaceae | USA (F) | 251 |
| Phidippus audax (Hentz, 1845) | Purpureocillium atypicola [= Nomuraea atypicola] | Ophiocordycipitaceae | USA (L) | 252 |
| Phidippus clarus Keyserling, 1885 | Purpureocillium atypicola [= Nomuraea atypicola] | Ophiocordycipitaceae | USA (L) | 253 |
| Phidippus clarus Keyserling, 1885 | Purpureocillium atypicola [= Nomuraea atypicola] | Ophiocordycipitaceae | USA (F) | 254 |
| Phidippus otiosus (Hentz, 1846) | Acrodontium crateriforme | Teratosphaeriaceae | USA (F) | 255 |
| Phidippus putnami (G. W. Peckham & E. G. Peckham, 1883) | Gibellula sp. | Cordycipitaceae | USA (F) | 256 |
| Phidippus regius C. L. Koch, 1846 | Acrodontium crateriforme | Teratosphaeriaceae | USA (F) | 257 |
| Phidippus regius C. L. Koch, 1846 | Purpureocillium atypicola [= Nomuraea atypicola] | Ophiocordycipitaceae | USA (F) | 258 |
| Phidippus sp. | Purpureocillium atypicola [= Nomuraea atypicola] | Ophiocordycipitaceae | USA (L) | 259 |

| | | | | |
|---|---|---|---|---|
| Platycryptus sp. | Gibellula pulchra | Cordycipitaceae | USA, LA (F) | 260 |
| Portia sp. | Gibellula scorpioides | Cordycipitaceae | Thailand (F) | 261 |
| N/A | Hymenostilbe sp. | Ophiocordycipitaceae | Brazil (F) | 262 |
| N/A | Akanthomyces aranearum | Cordycipitaceae | Thailand (F) | 263 |
| N/A | Akanthomyces aranearum | Cordycipitaceae | Thailand (F) | 264 |
| N/A | Akanthomyces koratensis | Cordycipitaceae | Thailand (F) | 265 |
| N/A | Cordyceps sp. | Cordycipitaceae | Ecuador (F) | 266 |
| N/A | Cordyceps sp. | Cordycipitaceae | Indonesia (F) | 267 |
| N/A | Cordyceps sp. | Cordycipitaceae | Singapore ? (F) | 268 |
| N/A | Cordyceps sp. | Cordycipitaceae | Unknown location (F) | 269 |
| N/A | Gibellula brunnea | Cordycipitaceae | Brazil (F) | 270 |
| N/A | Gibellula clavata | Cordycipitaceae | Ecuador (F) | 271 |
| N/A | Torrubiella clavata | Cordycipitaceae | Ecuador (F) | 272 |
| N/A | Gibellula leiopus | Cordycipitaceae | Philippines (F) | 273 |
| N/A | Gibellula leiopus | Cordycipitaceae | USA (F) | 274 |
| N/A | Gibellula mainsii | Cordycipitaceae | Brazil (F) | 275 |
| N/A | Gibellula mirabilis | Cordycipitaceae | Ecuador (F) | 276 |
| N/A | Gibellula pilosa | Cordycipitaceae | Thailand (F) | 277 |
| N/A | Gibellula pulchra | Cordycipitaceae | Thailand (F) | 278 |
| N/A | Gibellula pulchra | Cordycipitaceae | South Africa (F) | 279 |
| N/A | Gibellula pulchra | Cordycipitaceae | Taiwan (F) | 280 |
| N/A | Gibellula pulchra | Cordycipitaceae | Ghana (F) | 281 |
| N/A | Gibellula pulchra | Cordycipitaceae | South Africa (F) | 282 |
| N/A | Gibellula pulchra? | Cordycipitaceae | USA (F) | 283 |
| N/A | Gibellula pulchra? | Cordycipitaceae | Malaysia (F) | 284 |
| N/A | Gibellula trimorpha | Cordycipitaceae | Thailand (F) | 285 |
| N/A | Gibellula spp. | Cordycipitaceae | Brazil (F) | 286 |
| N/A | Gibellula spp. | Cordycipitaceae | Tropical region (F) | 287 |
| N/A | Gibellula sp. | Cordycipitaceae | Peru (F) | 288 |
| N/A | Gibellula sp. | Cordycipitaceae | Trinidad (F) | 289 |
| N/A | Gibellula cf. leiopus | Cordycipitaceae | USA (F) | 290 |
| N/A | Gibellula leiopus | Cordycipitaceae | Unknown location (F) | 291 |
| N/A | Gibellula cf. leiopus | Cordycipitaceae | USA (F) | 292 |
| N/A | Gibellula or Torrubiella | Cordycipitaceae | USA (F) | 293 |
| N/A | Granulomanus sp. | Cordycipitaceae | Peru (F) | 294 |
| N/A | Parahevansia koratensis [= Hevansia koratensis] | Cordycipitaceae | Thailand (F) | 295 |
| N/A | Pseudogibellula sp. | Cordycipitaceae | Peru (F) | 296 |
| N/A | Torrubiella sp. | Cordycipitaceae | Ecuador (F) | 297 |
| N/A | N/A | N/A | Australia (F) | 298 |
| N/A | Purpureocillium atypicola [= Nomuraea atypicola] | Ophiocordycipitaceae | USA (F) | 299 |
| N/A | Gibellula cf. leiopus | Cordycipitaceae | USA (F) | 300 |
| N/A | Gibellula sp. | Cordycipitaceae | USA (F) | 301 |
| N/A | Possibly Engyodontium aranearum [= Lecanicillium tenuipes] | Cordycipitaceae | Probably Singapore (F) | 302 |
| N/A | Gibellula cf. pulchra | Cordycipitaceae | Papua, Indonesia (F) | 303 |
| **SICARIIDAE** | | | | |
| Loxosceles reclusa Gertsch & | Purpureocillium | Ophiocordycipitaceae | USA (L) | 304 |

| | | | | |
|---|---|---|---|---|
| Mulaik, 1940 | atypicola [= Nomuraea atypicola] | | | |
| Loxosceles sp. | Metarhizium anisopliae | Clavicipitaceae | Brazil (L) | 305 |
| Loxosceles sp. | N/A | N/A | Peru (F) | 306 |
| **SPARASSIDAE** | | | | |
| Caayguara sp. | Gibellula spp. | Cordycipitaceae | Brazil (F) | 307 |
| Caayguara sp. | Gibellula sp. | Cordycipitaceae | Brazil (F) | 308 |
| Heteropoda jugulans (L. Koch, 1876) | Purpureocillium atypicola [= Nomuraea atypicola] | Ophiocordycipitaceae | Australia (F) | 309 |
| Palystes castaneus (Latreille, 1819) | Purpureocillium atypicola [= Nomuraea atypicola] | Ophiocordycipitaceae | South Africa (F) | 310 |
| N/A | Gibellula spp. | Cordycipitaceae | Brazil (F) | 311 |
| N/A | Purpureocillium atypicola [= Nomuraea atypicola] | Ophiocordycipitaceae | Australia (F) | 312 |
| N/A | Purpureocillium atypicola [= Nomuraea atypicola] | Ophiocordycipitaceae | Australia (F) | 313 |
| N/A | Purpureocillium atypicola [= Nomuraea atypicola] | Ophiocordycipitaceae | Australia (F) | 314 |
| N/A | Purpureocillium atypicola [= Nomuraea atypicola] | Ophiocordycipitaceae | Australia (F) | 315 |
| N/A | Purpureocillium atypicola [= Nomuraea atypicola] | Ophiocordycipitaceae | Papua New Guinea (F) | 316 |
| N/A | Purpureocillium atypicola [= Nomuraea atypicola] | Ophiocordycipitaceae | Peru (F) | 317 |
| N/A | Purpureocillium atypicola [= Nomuraea atypicola] | Ophiocordycipitaceae | Taiwan (F) | 318 |
| N/A | Purpureocillium atypicola [= Nomuraea atypicola] | Ophiocordycipitaceae | N/A (F) | 319 |
| **SYNOTAXIDAE** | | | | |
| Acrometa Petrunkevitch, 1942 | N/A | N/A | Baltic amber (F) | 320 |
| N/A | N/A | N/A | Baltic amber (F) | 321 |
| **TETRAGNATHIDAE** | | | | |
| Leucauge granulata (Walckenaer, 1841) | N/A | N/A | Australia (F) | 322 |
| Meta menardi (Latreille, 1804) | Gibellula sp. | Cordycipitaceae | Scotland (F) | 323 |
| Meta menardi (Latreille, 1804) | Gibellula bang-bangus | Cordycipitaceae | Scotland (F) | 324 |
| Meta menardi (Latreille, 1804) | Engyodontium rectidentatum | Cordycipitaceae | Czech Republic (F) | 325 |
| Meta menardi (Latreille, 1804) | Penicillium vulpinum | Aspergillaceae | Slovakia (F) | 326 |
| Meta menardi (Latreille, 1804) | Torrubiella arachnophila var. leiopus [= Torrubiella leiopus] | Cordycipitaceae | Germany (F) | 327 |
| Meta ovalis (Gertsch, 1933) | Beauveria spp. | Cordycipitaceae | USA (F) | 328 |
| Meta ovalis (Gertsch, 1933) | A pathogen in the | Ascomycota – | USA (F) | 329 |

| | | | | |
|---|---|---|---|---|
| | original publication termed as Paecilomyces | Incertae sedis | | |
| Metellina merianae (Scopoli, 1763) | Gibellula cf. leiopus | Cordycipitaceae | Wales, UK (F) | 330 |
| Metellina merianae (Scopoli, 1763) | Torrubiella arachnophila var. leiopus [= Torrubiella leiopus] | Cordycipitaceae | Germany (F) | 331 |
| Meta sp. | Torrubiella arachnophila var. leiopus [= Torrubiella leiopus] | Cordycipitaceae | Germany (F) | 332 |
| Pachygnatha degeeri Sundevall, 1830 | N/A | Class Hyphomycetes / unknown family | Netherlands (F) | 333 |
| Tetragnatha laboriosa Hentz, 1850 | Purpureocillium atypicola [= Nomuraea atypicola] | Ophiocordycipitaceae | USA (L) | 334 |
| N/A | Purpureocillium atypicola [= Nomuraea atypicola] | Ophiocordycipitaceae | Thailand /F) | 335 |
| **THERAPHOSIDAE** | | | | |
| Aphonopelma gabeli Smith, 1995 | Unidentified (Lecanicillium or Engyodontium) | Cordycipitaceae | England (L) | 336 |
| Avicularia juruensis Mello-Leitão, 1923 | Aspergillus niger | Aspergillaceae | Brazil (F) | 337 |
| Avicularia juruensis Mello-Leitão, 1923 | Beauveria bassiana | Cordycipitaceae | Brazil (F) | 338 |
| Grammostola sp. | Cordyceps caloceroides [= Ophiocordyceps caloceroides] | Ophiocordycipitaceae | Brazil (F) | 339 |
| Pamphobeteus ferox (Ausserer, 1875) | Cordyceps sp. | Cordycipitaceae | Colombia (F) | 340 |
| Phormictopus auratus Ortiz & Bertani, 2005 | N/A | N/A | Cuba (F) | 341 |
| Pterinopelma vitiosum (Keyserling, 1891) | Cordyceps caloceroides [= Ophiocordyceps caloceroides] | Ophiocordycipitaceae | Brazil (F) | 342 |
| N/A (subfamjily Theraphosinae) | Cordyceps caloceroides [= Ophiocordyceps caloceroides] | Cordycipitaceae | Colombia (F) | 343 |
| N/A | Cordyceps ignota | Cordycipitaceae | Argentina (F) | 344 |
| N/A | Cordyceps nidus | Cordycipitaceae | Colombia (L) | 345 |
| N/A | N/A | N/A | Ecuador (F) | 346 |
| **THERIDIIDAE** | | | | |
| Parasteatoda tepidariorum (C. L. Koch, 1841) [= Achaearanea tepidariorum] | Purpureocillium atypicola [= Nomuraea atypicola] | Ophiocordycipitaceae | USA (L) | 347 |
| Achaearanea sp. | Parengyodontium album [= Beauveria alba = Engyodontium album] | Cordycipitaceae | India (F) | 348 |
| Achaearanea sp. | Parengyodontium | Cordycipitaceae | Ukraine (F) | 349 |

| | | | | |
|---|---|---|---|---|
| | album [= Beauveria alba = Engyodontium album] | | | |
| Neopisinus cognatus (O. Pickard-Cambridge, 1893) [= Episinus cognatus] | Gibellula sp. | Cordycipitaceae | Brazil (F) | 350 |
| Helvibis longicauda Keyserling, 1891 | Gibellula pulchra | Cordycipitaceae | Brazil (F) | 351 |
| Hetschkia gracilis Keyserling, 1886 | Gibellula sp. | Cordycipitaceae | Brazil (F) | 352 |
| Janula bicornigera (Simon, 1894) | Gibellula sp. | Cordycipitaceae | Brazil (F) | 353 |
| Latrodectus geometricus C. L. Koch, 1841 | Mucor fragilis | Mucoraceae | USA (F) | 354 |
| Latrodectus geometricus C. L. Koch, 1841 | Mucor fragilis | Mucoraceae | USA (L) | 355 |
| Meotipa sp. | Hevansia minuta | Cordycipitaceae | Thailand (F) | 356 |
| Nesticodes rufipes (Lucas, 1846) | Clathroconium sp. | Incertae sedis | Cuba (F) | 357 |
| Theridion evexum Keyserling, 1884 | Gibellula sp. | Cordycipitaceae | Brazil (F) | 358 |
| N/A | Gibellula parvula | Cordycipitaceae | Thailand (F) | 359 |
| N/A | Gibellula solita | Cordycipitaceae | Thailand (F) | 360 |
| N/A | Hevansia novoguineensis | Cordycipitaceae | Thailand (F) | 361 |
| **THOMISIDAE** | | | | |
| Amyciaea sp. | Jenniferia cinerea [= Hevansia cinerea] | Cordycipitaceae | Thailand (F) | 362 |
| Cebrenninus cf. magnus Benjamin, 2016 | Gibellula cebrennini | Cordycipitaceae | Thailand (F) | 363 |
| Diaea cf. dorsata (Fabricius, 1777) | Jenniferia griseocinerea | Cordycipitaceae | Thailand (F) | 364 |
| Diaea cf. dorsata (Fabricius, 1777) | Jenniferia thomisidarum | Cordycipitaceae | Thailand (F) | 365 |
| Indoxysticus sp. | Gibellula longicaudata | Cordycipitaceae | Thailand (F) | 366 |
| Misumenops sp. | Purpureocillium atypicola [= Nomuraea atypicola] | Ophiocordycipitaceae | USA (L) | 367 |
| Synema parvulum (Hentz, 1847) | Most likely Purpureocillium atypicola [= Nomuraea atypicola] | Ophiocordycipitaceae | USA (F) | 368 |
| Tmarus sp. | Gibellula mainsii | Cordycipitaceae | Brazil (F) | 369 |
| Xysticus sp. | Purpureocillium atypicola [= Nomuraea atypicola] | Ophiocordycipitaceae | USA (L) | 370 |
| N/A | Cordyceps sp. ? | Cordycipitaceae | USA (F) | 371 |
| N/A | Gibellula brevistipitata | Cordycipitaceae | Thailand (F) | 372 |
| N/A | Gibellula pulchra | Cordycipitaceae | Poland (F) | 373 |
| N/A | Gibellula pulchra | Cordycipitaceae | Poland (F) | 374 |
| N/A | Gibellula sp. | Cordycipitaceae | Ecuador (F) | 375 |
| N/A | Gibellula or Torrubiella | Cordycipitaceae | USA, TN (F) | 376 |
| N/A ? | Purpureocillium atypicola [= Nomuraea atypicola] | Ophiocordycipitaceae | Indonesia (F) | 377 |
| N/A | Torrubiella albolanata | Cordycipitaceae | England (F) | 378 |
| N/A | Torrubiella | Cordycipitaceae | Japan (F) | 379 |

| | | | | |
|---|---|---|---|---|
| | neofusiformis | | | |
| **TRACHELIDAE** | | | | |
| Trachelas cf. robustus Keyserling, 1891 | Gibellula leiopus | Cordycipitaceae | Brazil (F) | 380 |
| Trachelas tranquillus (Hentz, 1847) | Gibellula leiopus | Cordycipitaceae | USA, LA (F) | 381 |
| Trachelas sp. | Gibellula leiopus | Cordycipitaceae | USA, MA (F) | 382 |
| Trachelas sp. | Gibellula leiopus | Cordycipitaceae | USA, SC (F) | 383 |
| Trachelas sp. | Immature Gibellula | Cordycipitaceae | USA, Georgia (F) | 384 |
| Trachelas sp. | immature Gibellula or Hevansia | Cordycipitaceae | USA, NC (F) | 385 |
| Trachelas sp. | N/A | N/A | USA, Maryland (F) | 386 |
| **TRECHALEIDAE** | | | | |
| Cupiennius salei (Keyserling, 1877) | Mucor hiemalis | Mucoraceae | Germany (L) | 387 |
| Cupiennius sp. | N/A | N/A | Ecuador (F) | 388 |
| Cupiennius sp. | Purpureocillium atypicola [= Nomuraea atypicola] | Ophiocordycipitaceae | Colombia (F) | 389 |
| **ULOBORIDAE** | | | | |
| Miagrammopes sp. | Gibellula dimorpha | Cordycipitaceae | Thailand (F) | 390 |
| N/A | Unidentified [Engyodontium or Lecanicillium] | Cordycipitaceae | Colombia (F) | 391 |
| **ZODARIIDAE** | | | | |
| Anniculus balticus Petrunkevitch 1942 | N/A | N/A | Baltic amber (F) | 392 |
| Epicratinus sp. | Gibellula sp. | Cordycipitaceae | Brazil (F) | 393 |
| Storenomorpha sp. | Gibellula pigmentosinum | Cordycipitaceae | Thailand (F) | 394 |
| **UNKNOWN SPIDERS** | | | | |
| N/A | Akanthomyces coccidioperitheciatus | Cordycipitaceae | Japan (F) | 395 |
| N/A | Akanthomyces kanyawimiae | Cordycipitaceae | Thailand (F) | 396 |
| N/A | Akanthomyces lecanii [= Lecanicillium lecanii = Verticillium lecanii] | Cordycipitaceae | Galapagos Islands, Ecuador, Brazil (F) | 397 |
| N/A | Akanthomyces ryukyuensis | Cordycipitaceae | Japan (F) | 398 |
| N/A | Akanthomyces sulphureus | Cordycipitaceae | Thailand () | 399 |
| N/A | Akanthomyces thailandicus | Cordycipitaceae | Thailand (F) | 400 |
| N/A | Akanthomyces waltergamsii | Cordycipitaceae | Thailand (F) | 401 |
| N/A | Beauveria araneola | Cordycipitaceae | China (F | 402 |
| N/A | Clonostachys aranearum | Bionectriaceae | China (F) | 403 |
| N/A | Cordyceps arachnogena | Cordycipitaceae | Papua New Guinea (F) | 404 |
| N/A | Cordyceps araneae | Cordycipitaceae | Thailand (F) | 405 |
| N/A | Cordyceps grenadensis | Cordycipitaceae | Grenada (F) | 406 |
| N/A | Cordyceps kuiburiensis | Cordycipitaceae | Thailand (F) | 407 |
| N/A | Cordyceps ogurasanensis | Cordycipitaceae | Japan (F) | 408 |

| | | | | |
|---|---|---|---|---|
| N/A | Cordyceps pseudonelumboides | Cordycipitaceae | Japan (F) | 409 |
| N/A | Cordyceps singeri | Cordycipitaceae | Argentina (F) | 410 |
| N/A | Gibellula alata | Cordycipitaceae | Sri Lanka (F) | 411 |
| N/A | Gibellula alata | Cordycipitaceae | Ghana (F) | 412 |
| N/A | Gibellula clavispora | Cordycipitaceae | China (F) | 413 |
| N/A | Gibellula clavulifera | Cordycipitaceae | Sri Lanka (F) | 414 |
| N/A | Gibellula clavulifera | Cordycipitaceae | Ghana (F) | 415 |
| N/A | Gibellula clavulifera | Cordycipitaceae | Thailand (F) | 416 |
| N/A | Gibellula clavulifera | Cordycipitaceae | China (F) | 417 |
| N/A | Gibellula curvispora | Cordycipitaceae | China (F) | 418 |
| N/A | Gibellula dabieshanensis | Cordycipitaceae | China (F) | 419 |
| N/A | Gibellula gamsii | Cordycipitaceae | Thailand (F) | 420 |
| N/A | Gibellula penicillioides | Cordycipitaceae | China (F) | 421 |
| N/A | Gibellula shennongjiaensis | Cordycipitaceae | China (F) | 422 |
| N/A | Hevansia arachnophila | Cordycipitaceae | Sri Lanka (F) | 423 |
| N/A | Hevansia arachnophila | Cordycipitaceae | Ghana (F) | 424 |
| N/A | Hevansia arachnophila | Cordycipitaceae | Thailand (F) | 425 |
| N/A | Hevansia arachnophila | Cordycipitaceae | Taiwan (F) | 426 |
| N/A | Hevansia arachnophila | Cordycipitaceae | Japan (F) | 427 |
| N/A | Jenniferia cinerea [= Hevansia cinerea] | Cordycipitaceae | Thailand (F) | 428 |
| N/A | Hevansia longispora | Cordycipitaceae | China (F) | 429 |
| N/A | Hevansia nelumboides | Cordycipitaceae | Japan (F) | 430 |
| N/A | Hevansia nelumboides | Cordycipitaceae | Taiwan (F) | 431 |
| N/A | Hevansia nelumboides | Cordycipitaceae | Thailand (F) | 432 |
| N/A | Hevansia nelumboides | Cordycipitaceae | China (F) | 433 |
| N/A | Hevansia ovalongata | Cordycipitaceae | Taiwan (F) | 434 |
| N/A | Hevansia ovalongata | Cordycipitaceae | Japan (F) | 435 |
| N/A | Hevansia websteri | Cordycipitaceae | Thailand (F) | 436 |
| N/A | Cordyceps farinosa [= Isaria farinosa= Poecilomyces farinosus] | Cordycipitaceae | Ghana (F) | 437 |
| N/A | Cordyceps farinosa [= Isaria farinosa= Poecilomyces farinosus] | Cordycipitaceae | Galapagos Islands (F) | 438 |
| N/A | Cordyceps javanica [= Isaria javanica] | Cordycipitaceae | Vietnam (F) | 439 |
| N/A | Lecanicillium aranearum | Cordycipitaceae | Sri Lanka (F) | 440 |
| N/A | Lecanicillium aranearum | Cordycipitaceae | Ghana (F) | 441 |
| N/A | Lecanicillium aranearum | Cordycipitaceae | India (F) | 442 |
| N/A | Lecanicillium araneicola | Cordycipitaceae | Indonesia (F) | 443 |
| N/A | Lecanicillium huhutii | Cordycipitaceae | China (F) | 444 |
| N/A | Akanthomyces lecanii [= Lecanicillium lecanii = Verticillium lecanii] | Cordycipitaceae | China (F) | 445 |
| N/A | Neoaraneomyces araneicola | Clavicipitaceae | China (F) | 446 |

| | | | | |
|---|---|---|---|---|
| N/A | Polystromomyces araneae | Cordycipitaceae | Thailand (F) | 447 |
| N/A | Pseudogibellula formicarum | Cordycipitaceae | Ghana (F) | 448 |
| N/A | Pseudometarhizium araneogenum | Clavicipitaceae | China (F) | 449 |
| N/A | Torrubiella alboglobosa | Cordycipitaceae | Japan (F) | 450 |
| N/A | Torrubiella aranicida | Cordycipitaceae | France (F) | 451 |
| N/A | Torrubiella aranicida | Cordycipitaceae | Cuba (F) | 452 |
| N/A | Torrubiella aranicida | Cordycipitaceae | China (F) | 453 |
| N/A | Torrubiella aranicida | Cordycipitaceae | UK (F) | 454 |
| N/A | Torrubiella aranicida | Cordycipitaceae | Japan (F) | 455 |
| N/A | Torrubiella aurantia | Cordycipitaceae | Japan (F) | 456 |
| N/A | Torrubiella aurantia | Cordycipitaceae | Thailand (F) | 457 |
| N/A | Torrubiella corniformis | Cordycipitaceae | Japan (F) | 458 |
| N/A | Torrubiella ellipsoidea | Cordycipitaceae | Japan (F) | 459 |
| N/A | Torrubiella falklandica | Cordycipitaceae | Falkland Islands (F) | 460 |
| N/A | Torrubiella farinacea | Cordycipitaceae | Japan (F) | 461 |
| N/A | Cordyceps flavoviridis [= Torrubiella flavoviridis] | Cordycipitaceae | Brazil (F) | 462 |
| N/A | Cordyceps flavoviridis [= Torrubiella flavoviridis] | Cordycipitaceae | Guyana (F) | 463 |
| N/A | Torrubiella formosana | Cordycipitaceae | Taiwan (F) | 464 |
| N/A | Torrubiella globosoides | Cordycipitaceae | Japan (F) | 465 |
| N/A | Cordyceps gonylepticida [= Torrubiella gonylepticida] | Cordycipitaceae | Brazil (F) | 466 |
| N/A | Cordyceps gonylepticida [= Torrubiella gonylepticida] | Cordycipitaceae | Trinidad (F) | 467 |
| N/A | Cordyceps gonylepticida [= Torrubiella gonylepticida] | Cordycipitaceae | Russian Caucasus (F) | 468 |
| N/A | Cordyceps gonylepticida [= Torrubiella gonylepticida] | Cordycipitaceae | Taiwan (F) | 469 |
| N/A | Torrubiella inegoensis | Cordycipitaceae | Japan (F) | 470 |
| N/A | Torrubiella longissima | Cordycipitaceae | Japan (F) | 471 |
| N/A | Torrubiella mammillata | Cordycipitaceae | Japan (F) | 472 |
| N/A | Torrubiella minuta | Cordycipitaceae | Japan (F) | 473 |
| N/A | Torrubiella miyagiana | Cordycipitaceae | Japan (F) | 474 |
| N/A | Torrubiella oblonga | Cordycipitaceae | Japan (F) | 475 |
| N/A | Torrubiella ooaniensis | Cordycipitaceae | Japan (F) | 476 |
| N/A | Torrubiella pallida | Cordycipitaceae | Japan (F) | 477 |
| N/A | Torrubiella plana | Cordycipitaceae | Japan (F) | 478 |
| N/A | Torrubiella plana | Cordycipitaceae | Taiwan (F) | 479 |
| N/A | Torrubiella rokkiana | Cordycipitaceae | Taiwan (F) | 480 |
| N/A | Torrubiella rosea | Cordycipitaceae | Japan (F) | 481 |

| | | | | |
|---|---|---|---|---|
| N/A | Torrubiella ryogamimontana | Cordycipitaceae | Japan (F) | 482 |
| N/A | Aphanocladium album | Nectriaceae | Khazakstan (F) | 483 |
| N/A | Hirsutella darwinii | Ophiocordycipitaceae | Galapagos Islands (F) | 484 |
| N/A | Hymenostilbe kedrovensis | Ophiocordycipitaceae | Russia (F) | 485 |
| N/A | Ophiocordyceps aranearum | Ophiocordycipitaceae | UK (F) | 486 |
| N/A | Ophiocordyceps engleriana | Ophiocordycipitaceae | Cameroon (F) | 487 |
| N/A | Ophiocordyceps engleriana | Ophiocordycipitaceae | Guyana (F) | 488 |
| N/A | Ophiocordyceps ghanensis | Ophiocordycipitaceae | Ghana (F) | 489 |
| N/A | Ophiocordyceps mrciensis | Ophiocordycipitaceae | Thailand (F) | 490 |
| N/A | Ophiocordyceps spiculata | Ophiocordycipitaceae | China (F) | 491 |
| N/A | Ophiocordyceps verrucosa | Ophiocordycipitaceae | USA (F) | 492 |
| N/A | Ophiocordyceps verrucosa | Ophiocordycipitaceae | UK (F) | 493 |
| N/A | Ophiocordyceps verrucosa | Ophiocordycipitaceae | China (F) | 494 |
| N/A | Tolypocladium cylindrosporum | Ophiocordycipitaceae | Europe (F) | 495 |
| N/A | Clathroconium arachnicola | Incertae sedis | Ghana (F) | 496 |
| N/A | Cryptococcus depauperatus [= Filobasidiella arachnophila] | Cryptococcaceae | Canada (F | 497 |
| N/A | Cladosporium cladosporioides | Cladosporiaceae | Canada (F) | 498 |
| N/A | Akanthomyces lecanii [= Lecanicillium lecanii = Verticillium lecanii] | Cordycipitaceae | Canada (F) | 499 |
| N/A | Penicillium tealii | Aspergillaceae | Australia (F) | 500 |
| N/A | Conidiobolus coronatus | Conidiobolaceae | Canada (F) | 501 |
| N/A | Cladosporium zixishanense | Cladosporiaceae | China (F) | 502 |
| N/A | Cordyceps cateniannulata | Cordycipitaceae | Thailand (F) | 503 |
| N/A | Aspergillus creber [= Aspergillus tennesseensis] | Aspergillaceae | Romania (F) | 504 |
| N/A | Clonostachys chuyangsinensis | Bionectriaceae | Vietnam (F) | 505 |
| N/A | Akanthomyces tiankengensis | Cordycipitaceae | China (F) | 506 |
| N/A | Akanthomyces bashanensis | Cordycipitaceae | China (F) | 507 |
| N/A | Akanthomyces beibeiensis | Cordycipitaceae | China (F) | 508 |
| N/A | Akanthomyces kunmingensis | Cordycipitaceae | China (F) | 509 |

| | | | | |
|---|---|---|---|---|
| N/A | Akanthomyces subaraneicola | Cordycipitaceae | China (F) | 510 |
| N/A egg sacs | Bhushaniella rubra | Cordycipitaceae | Thailand (F) | 511 |

016

https://www.reddit.com/r/Entomology/comments/o32x4f/can_anyone_help_id_this_spider_killed_by/

017  Mendes-Pereira T, De Araújo, JP, Mendes FC, Fonseca EO, Alves J, Sobczak JF et al. 2022. *Gibellula aurea* sp. nov. (Ascomycota, Cordycipitaceae): a new golden spider-devouring fungus from a Brazilian Atlantic Rainforest. *Phytotaxa* 573:85–102.

018  Arruda IDP. 2020. Manipulação comportamental da aranha *Macrophyes pacoti* (Araneae: Anyphaenidae) pelo fungo araneopatogênico *Gibellula* sp.(Hypocreales: Cordycipitaceae). Mestre Thesis, Universidade Federal do Ceará, Fortaleza, Brazil.

019  Arruda IDP, Villanueva-Bonilla GA, Faustino ML, Moura-Sobczak JCMS, Sobczak JF. 2021. Behavioral manipulation of the spider *Macrophyes pacoti* (Araneae: Anyphaenidae) by the araneopathogenic fungus *Gibellula* sp. (Hypocreales: Cordycipitaceae). *Canadian Journal of Zoology* 99:401–408.

020  Brescovit AD, Villanueva-Bonilla GA, Sobczak JCM, Nóbrega FADS, Oliveira LFMD, Arruda IDP et al. 2019. *Macrophyes pacoti* n. sp.(Araneae: Anyphaenidae) from Brazilian Atlantic Forest, with notes on an araneopathogenic fungus. *Zootaxa* 4629:294–300.

021  Costa PP. 2014. *Gibellula* spp. associadas a aranhas da Mata do Paraíso, Viçosa-MG. MSc Thesis, Universidade Federal de Viçosa, Brazil.

022  Hughes DP, Araújo JPM, Loreto RG, Quevillon L, De Bekker C, Evans HC. 2016. From so simple a beginning: the evolution of behavioral manipulation by fungi. *Advances in Genetics* 94:437–469.

023  Rose S. 2022. Spiders of North America. Princeton University Press, Princeton & Oxford.

024  Greenstone MH, Ignoffo CM, Samson RA. 1987. Susceptibility of spider species to the fungus *Nomuraea atypicola*. *Journal of Arachnology* 15:266–268.

025  Kobayasi Y. 1941. The genus *Cordyceps* and its allies. *Science Reports of the Tokyo Bunrika Daigaku, Section B* 84:53–260.

026  Chen WH, Liu C, Han YF, Liang JD, Liang ZQ. 2018. *Akanthomyces araneogenum*, a new Isaria-like araneogenous species. *Phytotaxa* 379:66–72.

027  Chen WH, Han YF, Liang ZQ, Jin DC. 2017. A new araneogenous fungus in the genus *Beauveria* from Guizhou, China. *Phytotaxa* 302:57–64.

028  Nentwig W. 1985a. Parasitic fungi as a mortality factor of spiders. *Journal of Arachnology* 13:272–274.

029  Greenstone MH, Ignoffo CM, Samson RA. 1987. Susceptibility of spider species to the fungus *Nomuraea atypicola*. *Journal of Arachnology* 15:266–268.

030  https://www.discoverlife.org/mp/20p?see=I_JP157820&res=640&flags=glean:

031–032   Nentwig W. 1985a. Parasitic fungi as a mortality factor of spiders. *Journal of Arachnology* 13:272–274.

033  Costa PP. 2014. *Gibellula* spp. associadas a aranhas da Mata do Paraíso, Viçosa-MG. MSc Thesis, Universidade Federal de Viçosa, Brazil.

034  Durkin ES, Cassidy ST, Gilbert R, Richardson EA, Roth AM, Shablin S et al. 2021. Parasites of spiders: Their impacts on host behavior and ecology. Journal of Arachnology 49:281–298.

035  Greenstone MH, Ignoffo CM, Samson RA. 1987. Susceptibility of spider species to the fungus *Nomuraea atypicola*. *Journal of Arachnology* 15:266–268.

036  Costa PP. 2014. *Gibellula* spp. associadas a aranhas da Mata do Paraíso, Viçosa-MG. MSc Thesis, Universidade Federal de Viçosa, Brazil.

037  Bishop L. 1990a. Entomophagous fungi as mortality agents of ballooning spiderlings. *Journal of Arachnology* 18:237–238

038  Humber RA, Hansen KS, Wheeler MM. 2014. USDA-ARS Collection of Entomopathogenic Fungal Cultures – Indexes to available isolates. Robert W. Holley Center for Agriculture and Health, Ithaca, New York.
https://www.ars.usda.gov/ARSUserFiles/80620520/ALL%20AVAIL%20indices%2016Jan014.pdf
Accessed 8 March 2023

039  https://whyevolutionistrue.com/2020/05/12/readers-wildlife-photos-1013/

040  https://aszk.org.au/wp-content/uploads/2020/03/Invertebrates.-Funnel-web-Spider-2009VB.pdf

041  Heneberg P, Řezáč M. 2013. Two *Trichosporon* species isolated from central-European mygalomorph spiders (Araneae: Mygalomorphae). *Antonie van Leeuwenhoek* 103:713–721.

042  Yasuda A. 1915. Purseweb spider parasitized by an *Isaria* fungus. *Botanical Magazine-Tokyo* 29:117.

043  Heneberg P, Řezáč M. 2013. Two *Trichosporon* species isolated from central-European mygalomorph spiders (Araneae: Mygalomorphae). *Antonie van Leeuwenhoek* 103:713–721.

044a–045a Pérez-Miles F., Perafán C. 2017. Behavior and biology of Mygalomorphae. Pp. 29–54. *In* Behaviour and Ecology of Spiders. (Viera C, Gonzaga MO, eds.). Springer, Cham.

044b–045b  Fernando Pérez-Miles, pers. comm.

046  Charles Haddad, pers. comm.

047  Steven Axford (photo) [ID Robert Raven]

048–049  Austin AD. 1984. Life history of *Clubiona robusta* L. Koch and related species (Araneae, Clubionidae) in South Australia. *Journal of Arachnology* 12:87–104.

050  van Helsdingen PJ. 2007. Sluipend gevaar. Nieuwsbrief Spined 23:35–35

051  Leatherdale D. 1970. The arthropod hosts of entomogenous fungi in Britain. *Entomophaga* 15:419–435.

052  McLean 1993 McLean IFG. 1993. A *Clubiona* spider infected with a parasitic fungus. *British Journal of Entomology and Natural History* 6:88.

053  van Vreden G, Ahmadzabidi AL. 1986. Pests of Rice and Their Natural Enemies in Peninsular Malaysia. Centre for Agricultural Publishing and Documentation (Pudoc), Wageningen, Netherlands.

054  Heinrichs EA. 1994. Biology and Management of Rice Insects. Wiley Eastern Limited, Delhi, India.

055  Mari S. 2017. 7. Observation particulière: Le champignon tueur d'araignées. Pp. 28–28. *In* Compte rendu des prospections «Araignées» 2016. https://oiseauxmaraisdharchies.be/onewebmedia/Rapport%20Inventaire%20arane%CC%81ologique%20Harchies%202016%20-finalized.pdf **

056  Barrion AT. 2001. Spiders: natural biological control agents against insect pests in Philippine rice fields. *Transactions of the National Academy of Science and Technology, Philippines* 23:121–130.

057  Bellmann H. 1997. Kosmos-Atlas Spinnentiere Europas. Franckh-Kosmos-Verlag, Stuttgart.

058  Mendes-Pereira T, De Araújo, JP, Mendes FC, Fonseca EO, Alves J, Sobczak JF et al. 2022. *Gibellula aurea* sp. nov. (Ascomycota, Cordycipitaceae): a new golden spider-devouring fungus from a Brazilian Atlantic Rainforest. *Phytotaxa* 573:85–102.

059  Wunderlich J. 2004. Fossil spiders in amber and copal: Conclusions, revisions, new taxa, family diagnoses of fossil and extant taxa. Beiträge zur Araneologie 3:1–1908.

060  https://www.alexanderwild.com/Insects/InsectKilling-Fungi/i-MhDvdgb/A

061–062  Sandler M. 2016. Zombie fungi: Occurrence of arthropod endoparasitic fungi at different altitudes in the Monteverde Region. Tropical Ecology Collection (Monteverde Institute). 641. https://digitalcommons.usf.edu/tropical_ecology/641

063  Hubert Höfer, pers. comm.

064–065  Arthur Decae, pers. comm.

066  https://www.inaturalist.org/observations/89047072

067  Robb Bennett, pers. comm.

068  https://www.alamy.com/scuttling-spider-infected-with-icing-sugar-fungus-image514844477.html

069  Kuephadungphan W, Tasanathai K, Petcharad B, Khonsanit A, Stadler M, Luangsa-ard JJ. 2020. Phylogeny- and morphology-based recognition of new species in the spider-parasitic genus *Gibellula* (Hypocreales, Cordycipitaceae) from Thailand. *MycoKeys* 72:17–42.

070  https://twitter.com/erincpow/status/1138628730803200002?lang=fr

071  Noordam AP, Samson RA, Sudhaus W. 1998. Fungi and Nematoda on *Centromerus sylvaticus* (Araneae, Linyphiidae). Pp. 343–347. *In* Proceedings of the 17th European Colloquium of Arachnology. (Selden PA, ed.). Edinburgh 1997.

072  Paz SN. 1993 Aspectos de la biologia reproductiva de *Linothele megatheloides* (Araneae: Dipluridae). *Journal of Arachnology* 21:40–49.

073
https://www.reddit.com/r/spiders/comments/ouo312/i_found_this_woodlouse_spider_overtaken_by_mold/

074 Henschel JR. 1998. Predation on social and solitary individuals of the spider *Stegodyphus dumicola* (Araneae, Eresidae). *Journal of Arachnology* 26:61–69.

075 https://bugguide.net/node/view/1209666

076 Wunderlich J. 2004. Fossil spiders in amber and copal: Conclusions, revisions, new taxa, family diagnoses of fossil and extant taxa. *Beiträge zur Araneologie* 3:1–1908.

077–078 Bristowe WL. 1958. The World of Spiders. Collins, London.

079 Cokendolpher JC. 2004. *Cicurina* spiders from caves in Bexar County, Texas (Araneae: Dictynidae). *Texas Memorial Museum Speleological Monographs* 6:13–58.

080 Wunderlich J. 2004. Fossil spiders in amber and copal: Conclusions, revisions, new taxa, family diagnoses of fossil and extant taxa. Beiträge zur Araneologie 3:1–1908.

081 https://www.inaturalist.org/observations/173218441

082 Kobayasi Y, Shimizu D. 1977. Some species of *Cordyceps* and its allies on spiders. *Kew Bulletin* 31:557–566.

083 Kobayasi Y. 1941. The genus *Cordyceps* and its allies. *Science Reports of the Tokyo Bunrika Daigaku, Section B* 84:53–260.

084 Petch T. 1939. Notes on entomogenous fungi. *Transactions of the British Mycological Society* 23:127–148.

085–086
https://www.naro.affrc.go.jp/org/fruit/epfdb/Ascomy/Cordyc/abcde/speci/OF0105.jpg

087 Haupt J. 2002. Fungal and rickettsial infections of some East Asian trapdoor spiders. Pp. 45–49. *In* European Arachnology 2000. (Toft S, Scharff N, eds.). Aarhus University Press, Aarhus.

088 Mains EB. 1954. Species of *Cordyceps* on spiders. *Bulletin of the Torrey Botanical Club* 81:492–500.

089 Kawamura S. 1929. On some new Japanese fungi. *Japanese Journal of Botany 4:* 291–302.

090a Yokoyama K, Ichikawa Y. 1984. Ecology of spider mushrooms at Omi Shrine in Otsu City. *Fuyu Kusa* 4:3–6. [in Japanese] Cited in:
https://www.wikieasy.wiki/de/%E3%82%AF%E3%83%A2%E3%82%BF%E3%82%B1

090b Yokoyama K, Hashiya M. 1994. Distribution survey of spider mushrooms. *Fuyu Kusa* 14:6–10. [in Japanese]. Cited in:
https://www.wikieasy.wiki/de/%E3%82%AF%E3%83%A2%E3%82%BF%E3%82%B1

091 https://rainforests.smugmug.com/Orders/Invertebrates/Orders/Cordyceps/i-FDVX5Kp/A

092 https://www.flickr.com/photos/rainforests/40151443503

093 Nigel Hywel-Jones (unpubl. data)

094 https://bugguide.net/node/view/1208519/bgimage

095 Orchard AE. 1996. Fungi of Australia. Australian Biological Resources Study, Canberra.

096a Pérez-Miles F., Perafán C. 2017. Behavior and biology of Mygalomorphae. Pp. 29–54. *In* Behaviour and Ecology of Spiders. (Viera C, Gonzaga MO, eds.). Springer, Cham.

096b Fernando Pérez-Miles, pers. comm.

097 Schwendinger PJ, Hongpadharakiree K. 2014. Three new *Prothemenops* species (Araneae: Idiopidae) from central Thailand. Zootaxa 3893:530-550.

098 Schwendinger PJ. 1996. The fauna of orthognathous spiders (Araneae: Mesothelae, Mygalomorphae) in Thailand. *Revue Suisse de Zoologie (Special Edition)* 2:577–584.

099 Castillo L, Sanjuan T, Restrepo S, Realpe E. 2015. Efecto del hongo aracnopatógeno *Cordyceps nidus* sp. nov. en tarántulas de la familia Theraphosidae en condiciones de laboratorio. Online at: https://repositorio.uniandes.edu.co/bitstream/handle/1992/17215/u703688.pdf?sequence=1&isAllowed=y Accessed 23 Sept 2022

100 Chirivi J, Danies G., Sierra R., Schauer N, Trenkamp S, Restrepo S et al. 2017. Metabolomic profile and nucleoside composition of *Cordyceps nidus* sp. nov. (Cordycipitaceae): a new source of active compounds. *PLoS One* 12:e0179428.

101 Nentwig W, Prillinger H. 1990. A zygomycetous fungus as a mortality factor in a laboratory stock of spiders. *Journal of Arachnology* 18:118–121.

102 https://www.stuff.co.nz/environment/17344/Fungus-to-kill-white-tail-spiders-discovered

103a Bristowe WS. 1948. Spiders from the arctic island of Jan Mayen. *Proceedings of the Zoological Society of London* 118:223–225.

103b Bristowe WL. 1958. The World of Spiders. Collins, London.

104 Barrion AT. 2001. Spiders: natural biological control agents against insect pests in Philippine rice fields. *Transactions of the National Academy of Science and Technology, Philippines* 23:121–130.

105 Heinrichs EA. 1994. Biology and Management of Rice Insects. Wiley Eastern Limited, Delhi, India.

106–108 Noordam AP, Samson RA, Sudhaus W. 1998. Fungi and Nematoda on *Centromerus sylvaticus* (Araneae, Linyphiidae). Pp. 343–347. *In* Proceedings of the 17th European Colloquium of Arachnology. (Selden PA, ed.). Edinburgh 1997.

109a-110a Bristowe WS. 1948. Spiders from the arctic island of Jan Mayen. *Proceedings of the Zoological Society of London* 118:223–225.

109b-110b Bristowe WL. 1958. The World of Spiders. Collins, London.

111  Læssøe T. 2015. Edderknoppe – Snyltekölle (*Torrubiella albolanata*) – en överraskelse fra de jyske sumpe. Svampe 71:23–27, 37.

112  Greenstone MH, Ignoffo CM, Samson RA. 1987. Susceptibility of spider species to the fungus *Nomuraea atypicola*. *Journal of Arachnology* 15:266–268.

113  Petch T. 1948. A revised list of British entomogenous fungi. *Transactions of the British Mycological Society* 31:286–304.

114  Petch T. 1944. Notes on entomogenous fungi. *Transactions of the British Mycological Society* 27:81–93.

115–116  Duffey E. 1997. Spider adaptation to artificial biotopes: the fauna of percolating filter beds in a sewage treatment works. *Journal of Applied Ecology* 34:1190–1202.

117  Noordam AP, Samson RA, Sudhaus W. 1998. Fungi and Nematoda on *Centromerus sylvaticus* (Araneae, Linyphiidae). Pp. 343–347. *In* Proceedings of the 17th European Colloquium of Arachnology. (Selden PA, ed.). Edinburgh 1997.

118–119  Bishop L. 1990a. Entomophagous fungi as mortality agents of ballooning spiderlings. *Journal of Arachnology* 18:237–238

120a  Bristowe WS. 1948. Spiders from the arctic island of Jan Mayen. *Proceedings of the Zoological Society of London* 118:223–225.

120b  Bristowe WL. 1958. The World of Spiders. Collins, London.

121–123  Noordam AP, Samson RA, Sudhaus W. 1998. Fungi and Nematoda on *Centromerus sylvaticus* (Araneae, Linyphiidae). Pp. 343–347. *In* Proceedings of the 17th European Colloquium of Arachnology. (Selden PA, ed.). Edinburgh 1997.

124  Meyling NV, Thorup-Kristensen K, Eilenberg J. 2011. Below-and aboveground abundance and distribution of fungal entomopathogens in experimental conventional and organic cropping systems. *Biological Control* 59:180–186.

125a  Bristowe WS. 1948. Spiders from the arctic island of Jan Mayen. *Proceedings of the Zoological Society of London* 118:223–225.

125b  Bristowe WL. 1958. The World of Spiders. Collins, London.

126–127  Leatherdale D. 1970. The arthropod hosts of entomogenous fungi in Britain. *Entomophaga* 15:419–435.

128  Ruszkiewicz-Michalska M, Tkaczuk C, Dynowska M, Sucharzewska E, Szkodzik J, Wrzosek M. 2012. Preliminary studies of fungi in the Biebrza National Park (NE Poland). I. Micromycetes. *Acta Mycologica* 47:213–234.

129  Kuephadungphan W, Petcharad B, Tasanathai K, Thanakitpipattana D, Kobmoo N, Khonsanit A. et al. 2022. Multi-locus phylogeny unmasks hidden species within the specialised spider-parasitic fungus, *Gibellula* (Hypocreales, Cordycipitaceae) in Thailand. Studies in Mycology 101:245–286.

130  Bosselaers JP. 1984. *Gibellula pulchra* (Sacc.) Cavara in het gebied van de Slangebeekbron te Zonhoven (België). *Natuurhistorisch Maandblad* 73:166–168.

131–132 Meyling NV, Thorup-Kristensen K, Eilenberg J. 2011. Below-and aboveground abundance and distribution of fungal entomopathogens in experimental conventional and organic cropping systems. *Biological Control* 59:180–186.

133 Costa PP. 2014. *Gibellula* spp. associadas a aranhas da Mata do Paraíso, Viçosa-MG. MSc Thesis, Universidade Federal de Viçosa, Brazil.

134 Thomas Laessoe (pers. comm./ photo)

135a Bristowe WS. 1948. Spiders from the arctic island of Jan Mayen. *Proceedings of the Zoological Society of London* 118:223–225.

135b Bristowe WL. 1958. The World of Spiders. Collins, London.

136a Nováková A, Kubátová A, Sklenář F, Hubka V. 2018a. Microscopic fungi on cadavers and skeletons from cave and mine environments. *Czech Mycology* 70:101–121.

136b Nováková A, Hubka V, Valinová Š, Kolařík M, Hillebrand-Voiculescu AM. 2018b. Cultivable microscopic fungi from an underground chemosynthesis-based ecosystem: a preliminary study. *Folia Microbiologica* 63:43–55.

137 Greenstone MH, Ignoffo CM, Samson RA. 1987. Susceptibility of spider species to the fungus *Nomuraea atypicola*. *Journal of Arachnology* 15:266–268.

138 Keller S, Wegensteiner R. 2010. A species of the fungus genus *Conidiobolus* as a pathogen of a lycosid spider. *Mitteilungen der Schweizerischen Entomologischen Gesellschaft* 83:227–231.

139 Edgar WD. 1969. Prey and predators of the wolf spider *Lycosa lugubris*. Journal of Zoology 159:405–411.

140 Noordam AP, Samson RA, Sudhaus W. 1998. Fungi and Nematoda on *Centromerus sylvaticus* (Araneae, Linyphiidae). Pp. 343–347. *In* Proceedings of the 17th European Colloquium of Arachnology. (Selden PA, ed.). Edinburgh 1997.

141 Savić D, Grbić G, Bošković E, Hänggi A. 2016. First records of fungi pathogenic on spiders for the Republic of Serbia. *Arachnology Letters* 52:31–34.

142 Doidge EM. 1950. Fungi on hosts other than vascular plants. Bothalia 5:58–63.

143 van der Bijl PA. 1922. A fungus - *Gibellula haygarthii*, sp. N. - on a spider of the family Lycosidae. *Transactions of the Royal Society of South Africa* 10:149–150.

144 https://bugguide.net/node/view/1707125

145 Hywel-Jones NL, Sivichai S. 1995. *Cordyceps cylindrica* and its association with *Nomuraea atypicola* in Thailand. *Mycological Research* 7:809–812.

146 Sandler M. 2016. Zombie fungi: Occurrence of arthropod endoparasitic fungi at different altitudes in the Monteverde Region. Tropical Ecology Collection (Monteverde Institute). 641. https://digitalcommons.usf.edu/tropical_ecology/641

147  Wunderlich J. 2004. Fossil spiders in amber and copal: Conclusions, revisions, new taxa, family diagnoses of fossil and extant taxa. Beiträge zur Araneologie 3:1–1908.

148  Isaia M, Decae A. 2012. Revalidation of *Nemesia meridionalis* Costa, 1835 (Araneae, Mygalomorphae, Nemesiidae), and first description of the male. *Arachnology* 15:280–284.

149  https://www.wikiwand.com/en/Beauveria_bassiana

150  Nentwig W. 1985a. Parasitic fungi as a mortality factor of spiders. *Journal of Arachnology* 13:272–274.

151  Humber RA, Hansen KS, Wheeler MM. 2014. USDA-ARS Collection of Entomopathogenic Fungal Cultures – Indexes to available isolates. Robert W. Holley Center for Agriculture and Health, Ithaca, New York.
https://www.ars.usda.gov/ARSUserFiles/80620520/ALL%20AVAIL%20indices%2016Jan014.pdf
Accessed 8 March 2023

152  https://mycocosm.jgi.doe.gov/Spoumb1/Spoumb1.home.html

153  https://twitter.com/phil_torres

154  https://bugguide.net/node/view/1573368/bgimage

155  Henriksen CB, Reboleira ASP, Scharff N, Enghoff H. 2018. First record of a Basidiobolus/Amphoromorpha fungus from a spider. *African Journal of Ecology* 56:153–156.

156  https://www.flickr.com/photos/rainforests/11307005053

157  https://bugguide.net/node/view/36643

158  https://jlrexplore.com/gallery/photostories/the-killer-fung

159  Kuephadungphan W, Petcharad B, Tasanathai K, Thanakitpipattana D, Kobmoo N, Khonsanit A. et al. 2022. Multi-locus phylogeny unmasks hidden species within the specialised spider-parasitic fungus, *Gibellula* (Hypocreales, Cordycipitaceae) in Thailand. Studies in Mycology 101:245–286.

160  https://www.flickr.com/photos/budak/41614482464/

161  Seth Ausubel, pers. comm.

162–163  Rong IH, Grobbelaar E. 1998. South African records of associations between fungi and arthropods. *African Plant Protection* 4:43–63.

164  https://www.flickr.com/photos/aracnologo/33172017034/

165  Costa PP. 2014. *Gibellula* spp. associadas a aranhas da Mata do Paraíso, Viçosa-MG. MSc Thesis, Universidade Federal de Viçosa, Brazil.

166  Mercado Sierra A, Alayo Soto R, Mena Portales J, de Armas LF. 1988. Hongos entomógenos de Cuba. Nueva especie de *Clathroconium* sobre arañas. *Acta Botánica Cubana* 56:1–5.

167–168  Jent D. 2013. Pathogenic Fungi Affecting Cellar Spiders, *Pholcus phalangioides*. Student project, Murray State University (Abstract). Online at:

https://digitalcommons.murraystate.edu/postersatthecapitol/2013/Murray/6/ Accessed 27 September 2022.

169 Ian Redding – https://www.alamy.com/stock-photo-daddy-longlegs-spider-pholcus-phalangioides-dead-and-covered-in-fungus-92890630.html

170 https://www.alamy.com/daddy-longlegs-spider-pholcus-phalangoides-infected-with-a-fungal-pathogen-possibly-gibellula-pulchra-in-a-cave-near-podere-montecucco-orvieot-umbria-italy-image262966101.html

171 https://www.youtube.com/watch?v=T9aw56z6jTg

172 https://www.flickr.com/photos/alan_cressler/22082836645

173 Martynenko SV, Kondratyukn TO, Sukhomlin MM. 2012. A hyphomycete, *Engyodontium album* (Limber) de Hoog, attacking spiders in underground headings of Kyiv-City. *Ukrainian Botanical Journal* 69:423–432.

174 Cokendolpher JC. 1993. Pathogens and parasites of opiliones (arthropoda: arachnida). *Journal of Arachnology* 21:120–146.

175 Dubiel G. 2015. Występowanie grzyba *Gibellula leiopus* (Vuil. ex Maubl.) Mains w Beskidzie Śląskim. *Przegląd Przyrodniczy* 26:30–38.

176 Eiseman C, Charney N, Carlson J. 2010. Tracks & Sign of Insects & Other Invertebrates: A Guide to North American Species. Stackpole Books, Mechanicsburg.

177 https://www.inaturalist.org/observations/124885254

178 https://www.inaturalist.org/observations/140295751

179 https://www.inaturalist.org/observations/130995484

180 https://www.inaturalist.org/observations/126673855

181 https://www.inaturalist.org/observations/124879856

182 https://www.inaturalist.org/observations/111079446

183 https://www.inaturalist.org/observations/110955846

184 https://www.inaturalist.org/observations/109295151

185 https://www.inaturalist.org/observations/106805425

186 https://www.inaturalist.org/observations/105038206

187 https://www.inaturalist.org/observations/102510550

188 https://www.inaturalist.org/observations/94084524

189 https://www.inaturalist.org/observations/82581919

190 https://www.inaturalist.org/observations/73504730

191 https://www.inaturalist.org/observations/71357780

192 https://www.inaturalist.org/observations/71017453

193 https://www.inaturalist.org/observations/66159361

194 https://www.inaturalist.org/observations/6492719538485

195 https://www.inaturalist.org/observations/64667449

196 https://www.inaturalist.org/observations/58144497

197 https://www.inaturalist.org/observations/47423815

198 https://www.inaturalist.org/observations/21948048

199 https://www.inaturalist.org/observations/141405381

200 https://www.inaturalist.org/observations/138848464

201 https://www.inaturalist.org/observations/138783675

202 https://www.inaturalist.org/observations/129578129

203 https://www.inaturalist.org/observations/114440448

204 https://www.inaturalist.org/observations/103106269

205 https://www.inaturalist.org/observations/101988244

206 https://www.inaturalist.org/observations/100592061

207 https://www.inaturalist.org/observations/97905071

208 https://www.inaturalist.org/observations/97615190

209 https://www.inaturalist.org/observations/96648400

210 https://www.inaturalist.org/observations/92354925

211 https://www.inaturalist.org/observations/65501463

212 https://www.inaturalist.org/observations/37388680

213 https://www.inaturalist.org/observations/36498174

214 https://www.inaturalist.org/observations/17493159

215 https://www.flickr.com/photos/nagysandor/29821742974

216 Humber RA, Hansen KS, Wheeler MM. 2014. USDA-ARS Collection of Entomopathogenic Fungal Cultures – Indexes to available isolates. Robert W. Holley Center for Agriculture and Health, Ithaca, New York.
https://www.ars.usda.gov/ARSUserFiles/80620520/ALL%20AVAIL%20indices%2016Jan014.pdf
Accessed 8 March 2023

217 Kuephadungphan W, Petcharad B, Tasanathai K, Thanakitpipattana D, Kobmoo N, Khonsanit A. et al. 2022. Multi-locus phylogeny unmasks hidden species within the specialised spider-parasitic fungus, *Gibellula* (Hypocreales, Cordycipitaceae) in Thailand. *Studies in Mycology* 101:245–286.

218 http://naturephoto-walter.blogspot.com/2017/02/miskolc-hungary-misk

219
https://www.reddit.com/r/ThatsInsane/comments/j4292e/this_is_the_fungus_engyodontium_aranearum_which/

220a https://blog.mycology.cornell.edu/2006/11/09/a-spiders-nightmare/

220b Kathie T. Hodge (pers. comm.)

221 Cokendolpher JC. 1993. Pathogens and parasites of opiliones (arthropoda: arachnida). *Journal of Arachnology* 21:120–146.

222 Rose S. 2022. Spiders of North America. Princeton University Press, Princeton & Oxford.

223 https://www.inaturalist.org/observations/166795631

224 https://www.inaturalist.org/observations/106805412

225 https://www.gbif.org/occurrence/4076204454

226 https://www.gbif.org/occurrence/4080930548

227 https://www.gbif.org/occurrence/4034771775

228a https://www.flickr.com/photos/gillesarbour/24503554811 no longer accessible June 2023

228b Gilles Arbour, pers. comm. (Fig. 5B, this paper)

229 https://www.inaturalist.org/observations/138327035

230–232 Manfrino RG, González A, Barneche J, Galván JT, Hywell-Jones N, Lastra CCL. 2017. Contribution to the knowledge of pathogenic fungi of spiders in Argentina. Southernmost record in the world. *Revista Argentina de Microbiología* 49:197–200.

233 Charles Haddad, pers. comm.

234 Wolcott GN. 1948. The insects of Puerto Rico. *Journal of Agriculture of the University of Puerto Rico* 32:1–224.

235 Saltamachia SJ. 2022. Theoretical and empirical evidence for extended phenotypes in a specialized parasite of spiders. *Authorea Preprint Repository* DOI: 10.22541/au.164604858.89088094/v1.

236  Rose S. 2022. Spiders of North America. Princeton University Press, Princeton & Oxford.

237  Saltamachia SJ. 2022. Theoretical and empirical evidence for extended phenotypes in a specialized parasite of spiders. *Authorea Preprint Repository* DOI: 10.22541/au.164604858.89088094/v1.

238  Carmen Champagne, pers. comm. (Fig. 4C, this paper).

239  Costa PP. 2014. *Gibellula* spp. associadas a aranhas da Mata do Paraíso, Viçosa-MG. MSc Thesis, Universidade Federal de Viçosa, Brazil.

240–241  Humber RA, Rombach MC. 1987. *Torrubiella ratticaudata* sp. nov.(Pyrenomycetes: Clavicipitales) and other fungi from spiders on the Solomon Islands. Mycologia 79:375–382.

242  Charles Haddad, pers. comm.

243  Greenstone MH, Ignoffo CM, Samson RA. 1987. Susceptibility of spider species to the fungus *Nomuraea atypicola*. *Journal of Arachnology* 15:266–268.

244  Durkin ES, Cassidy ST, Gilbert R, Richardson EA, Roth AM, Shablin S et al. 2021. Parasites of spiders: Their impacts on host behavior and ecology. Journal of Arachnology 49:281–298.

245  Greenstone MH, Ignoffo CM, Samson RA. 1987. Susceptibility of spider species to the fungus *Nomuraea atypicola*. *Journal of Arachnology* 15:266–268.

246  Mary Jane Hatfield, pers. comm. (Fig. 4A, this paper)

247  https://www.flickr.com/photos/72842252@N04/12691569924

248  Abhijith APC, Hill DE, Ramachandra P. 2002. Notes on biology of the ant-mimicking jumping spider *Myrmarachne plataleoides* (Araneae: Salticidae: Astioida) in south Asia. *Peckhamia* 287.1:1–12.

249  Kuephadungphan W, Petcharad B, Tasanathai K, Thanakitpipattana D, Kobmoo N, Khonsanit A. et al. 2022. Multi-locus phylogeny unmasks hidden species within the specialised spider-parasitic fungus, *Gibellula* (Hypocreales, Cordycipitaceae) in Thailand. *Studies in Mycology* 101:245–286.

250  Strongman DB. 1991. *Gibellula pulchra* from a spider (Salticidae) in Nova Scotia, Canada. *Mycologia* 83:816–817.

251  Edwards GB. 1980. Taxonomy, ethology, and ecology of *Phidippus* (Araneae: Salticidae) in eastern North America. PhD Dissertation, University of Florida, Gainesville, USA.

252–253  Greenstone MH, Ignoffo CM, Samson RA. 1987. Susceptibility of spider species to the fungus *Nomuraea atypicola*. *Journal of Arachnology* 15:266–268.

254–255  Edwards GB. 1980. Taxonomy, ethology, and ecology of *Phidippus* (Araneae: Salticidae) in eastern North America. PhD Dissertation, University of Florida, Gainesville, USA.

256  https://bugguide.net/node/view/828984

257–258  Edwards GB. 1980. Taxonomy, ethology, and ecology of *Phidippus* (Araneae: Salticidae) in eastern North America. PhD Dissertation, University of Florida, Gainesville, USA.

259  Greenstone MH, Ignoffo CM, Samson RA. 1987. Susceptibility of spider species to the fungus *Nomuraea atypicola*. *Journal of Arachnology* 15:266–268.

260  Saltamachia SJ. 2022. Theoretical and empirical evidence for extended phenotypes in a specialized parasite of spiders. *Authorea Preprint Repository* DOI: 10.22541/au.164604858.89088094/v1.

261  Kuephadungphan W, Tasanathai K, Petcharad B, Khonsanit A, Stadler M, Luangsa-Ard JJ. 2020. Phylogeny- and morphology-based recognition of new species in the spider-parasitic genus *Gibellula* (Hypocreales, Cordycipitaceae) from Thailand. *MycoKeys* 72:17–42.

262  Evans HC, Samson RA. 1987. Fungal pathogens of spiders. Mycologist 1:152–159.

263  Luangsa-Ard JJ, Tasanathai K, Mongkolsamrit S, Hywel-Jones N. 2008. Atlas of Invertebrate-Pathogenic Fungi of Thailand, Volume 2. National Center for Genetic Engineering and Biotechnology, National Science and Technology Development Agency, Pathumthani, Thailand.

264  http://www.thai2bio.net/museum/item.php?keyword=Akanthomyces%20aranearum

265  Hywel-Jones N. 1996. *Akanthomyces* on spiders in Thailand. *Mycological Research* 9:1065–1070.

266  https://www.alamy.de/fotos-bilder/spider-parasitized-cordyceps-fungus-in.html

267  https://rainforests.smugmug.com/Orders/Invertebrates/Orders/Cordyceps

268  https://www.deviantart.com/melvynyeo/art/Killer-Fungus-279883330

269  https://twitter.com/stephenmarek2/status/647981177056309248

270–272  Samson RA, Evans HC. 1992. New species of *Gibellula* on spiders (Araneida) from South America. *Mycologia* 84:300–314.

273  Barrion AT. 2001. Spiders: natural biological control agents against insect pests in Philippine rice fields. *Transactions of the National Academy of Science and Technology, Philippines* 23:121–130.

274  https://bugguide.net/node/view/236694

275–276  Samson RA, Evans HC. 1992. New species of *Gibellula* on spiders (Araneida) from South America. *Mycologia* 84:300–314.

277–278  Kuephadungphan W, Petcharad B, Tasanathai K, Thanakitpipattana D, Kobmoo N, Khonsanit A. et al. 2022. Multi-locus phylogeny unmasks hidden species within the specialised spider-parasitic fungus, *Gibellula* (Hypocreales, Cordycipitaceae) in Thailand. *Studies in Mycology* 101:245–286.

279  Rong IH, Grobbelaar E. 1998. South African records of associations between fungi and arthropods. *African Plant Protection* 4:43–63.

280  https://taieol.tw/pages/142073

281 Samson RA, Evans HC. 1973. Notes on entomogenous fungi from Ghana: I. The genera *Gibellula* and *Pseudogibellula*. *Acta Botanica Neerlandica* 22:522–528.

282 Rong IH, Botha A. 1993. New and interesting records of South African fungi XII. Synnematous Hyphomycetes. *South African Journal of Botany* 59:514–518.

283 https://www.marylandbiodiversity.com/view/15240

284 https://bugkeeping.tumblr.com/post/634138914125496321/a-jumping-spider-parasitized-by-a-cordyceps

285 Kuephadungphan W, Petcharad B, Tasanathai K, Thanakitpipattana D, Kobmoo N, Khonsanit A. et al. 2022. Multi-locus phylogeny unmasks hidden species within the specialised spider-parasitic fungus, *Gibellula* (Hypocreales, Cordycipitaceae) in Thailand. *Studies in Mycology* 101:245–286.

286 Costa PP. 2014. *Gibellula* spp. associadas a aranhas da Mata do Paraíso, Viçosa-MG. MSc Thesis, Universidade Federal de Viçosa, Brazil.

287 Evans HC, Samson RA. 1987. Fungal pathogens of spiders. *Mycologist* 1:152–159.

288 Pérez Meza P. 2004. Hongos entomopatógenos asociados a diferentes cultivos tropicales. Tesis para optar el título de Ingeniero Agrónomo, Universidad Nacional Agraria de la Selva, Peru.

289 Williams 1921 Williams CB. 1921. Report on the froghopper-blight of sugar-cane in Trinidad. *Trinidad and Tobago Department of Agriculture Memoirs* 1:1–170.

290 https://ozarkbill.com/2021/11/01/zombie-spider-bastards/

291
https://www.reddit.com/r/Parasitology/comments/q5lhbq/parasitic_gibellula_sp_fungus_in_a_jumping_spider/

292
https://www.reddit.com/r/natureismetal/comments/owcygc/spider_infected_with_gibellula_fungus/

293 Bishop L. 1990a. Entomophagous fungi as mortality agents of ballooning spiderlings. *Journal of Arachnology* 18:237–238.

294 Pérez Meza P. 2004. Hongos entomopatógenos asociados a diferentes cultivos tropicales. Tesis para optar el título de Ingeniero Agrónomo, Universidad Nacional Agraria de la Selva, Peru.

295 Hywel-Jones N. 1996. *Akanthomyces* on spiders in Thailand. *Mycological Research* 9:1065–1070.

296 Pérez Meza P. 2004. Hongos entomopatógenos asociados a diferentes cultivos tropicales. Tesis para optar el título de Ingeniero Agrónomo, Universidad Nacional Agraria de la Selva, Peru.

297 Evans HC, Samson RA. 1987. Fungal pathogens of spiders. *Mycologist* 1:152–159.

298 https://www.projectnoah.org/spottings/6423106

299 http://www.plantpath.cornell.edu/PhotoLab/PicOfMonth/POM3.htm

300 https://www.whatsthatbug.com/spider-with-fungus-infection/

301 https://www.marylandbiodiversity.com/view/15240

302 https://www.deviantart.com/melvynyeo/art/Killer-Fungus-279883330

303 https://www.projectnoah.org/spottings/7104616

304 Greenstone MH, Ignoffo CM, Samson RA. 1987. Susceptibility of spider species to the fungus *Nomuraea atypicola*. *Journal of Arachnology* 15:266–268.

305 Beys-da-Silva WO, Santi L, Berger M, Guimaraes JA, Schrank A, Vainstein MH. 2013. Susceptibility of Loxosceles sp. to the arthropod pathogenic fungus Metarhizium anisopliae: potential biocontrol of the brown spider. Transactions of the Royal Society of Tropical Medicine and Hygiene 107:59–61.

306 https://www.inaturalist.org/observations/34934503

307 Costa PP. 2014. *Gibellula* spp. associadas a aranhas da Mata do Paraíso, Viçosa-MG. MSc Thesis, Universidade Federal de Viçosa, Brazil.

308 Hughes DP, Araújo JPM, Loreto RG, Quevillon L, De Bekker C, Evans HC. 2016. From so simple a beginning: the evolution of behavioral manipulation by fungi. *Advances in Genetics* 94:437–469.

309 https://minibeastwildlife.blogspot.com/2011/03/killer-fungus.html

310 Rong IH, Grobbelaar E. 1998. South African records of associations between fungi and arthropods. *African Plant Protection* 4:43–63.

311 Costa PP. 2014. *Gibellula* spp. associadas a aranhas da Mata do Paraíso, Viçosa-MG. MSc Thesis, Universidade Federal de Viçosa, Brazil.

312 http://davidavid.blogspot.com/2008/

313 https://www.foxeslair.org/foxypress/archives/07-2020

314 https://twitter.com/gotlegsmacro/status/1472827370843168772?lang=ar

315 https://www.flickr.com/photos/72842252@N04/13997658232

316 https://www.whatsthatbug.com/2009/04/26/giant-crab-spider-riddled-with-fungus-we-believe/

317 https://www.flickr.com/photos/rainforests/51674043090/

318 https://www.123rf.com/photo_145569651_close-up-shot-of-a-huntsman-spiders-at-yilan-taiwan.html

319 https://www.shutterstock.com/image-photo/close-dead-huntsman-spider-fungus-on-1894778263

320–321 Wunderlich J. 2004. Fossil spiders in amber and copal: Conclusions, revisions, new taxa, family diagnoses of fossil and extant taxa. *Beiträge zur Araneologie* 3:1–1908.

322 https://www.flickr.com/photos/40132175@N06/4680651904

323 Harry Evans, pers. comm.

324 CABI 2022. Confirmed new fungus has mysterious origins. https://phys.org/news/2022-06-fungus-mysterious.html Accessed 3 August 2022.

325 Kubátová A. 2017. Entomopatogenní houby – nerovný souboj. *Ziva* 5:250–254.

326 Nováková A, Kubátová A, Sklenář F, Hubka V. 2018a. Microscopic fungi on cadavers and skeletons from cave and mine environments. Czech Mycology 70:101–121.

327 https://das-neue-naturforum.de/forum/index.php?thread/19258-die-spinnenkernkeule-torrubiella-leiopus-nebenfruchtform/

328–329 Yoder JA, Benoit JB, Christensen BS, Croxall TJ, Hobbs HH. 2009. Entomopathogenic fungi carried by the cave orb weaver spider, *Meta ovalis* (Araneae, Tetragnathidae), with implications for mycoflora transfer to cave crickets. *Journal of Cave and Karst Studies*, 71:116–120.

330 McNeil D. 2012. Entomogenous fungi. *Shropshire Entomology* 5:5–6.

331-332 https://das-neue-naturforum.de/forum/index.php?thread/19258-die-spinnenkernkeule-torrubiella-leiopus-nebenfruchtform/

333 Noordam AP, Samson RA, Sudhaus W. 1998. Fungi and Nematoda on *Centromerus sylvaticus* (Araneae, Linyphiidae). Pp. 343–347. *In* Proceedings of the 17th European Colloquium of Arachnology. (Selden PA, ed.). Edinburgh 1997.

334 Greenstone MH, Ignoffo CM, Samson RA. 1987. Susceptibility of spider species to the fungus *Nomuraea atypicola*. *Journal of Arachnology* 15:266–268.

335 Hywel-Jones NL, Sivichai S. 1995. *Cordyceps cylindrica* and its association with *Nomuraea atypicola* in Thailand. *Mycological Research* 7:809–812.

336 Sherwood D. 2021. Notes on a case of fungal pathogenesis on a juvenile of the theraphosid spider *Aphonopelma gabeli* Smith, 1995 in captivity (Araneae: Theraphosidae). *Serket* 18:27–30.

337–338 Ayroza G, Ferreira IL, Sayegh RS, Tashima AK, da Silva Jr, PI. 2012. Juruin: an antifungal peptide from the venom of the Amazonian Pink Toe spider, *Avicularia juruensis*, which contains the inhibitory cystine knot motif. *Frontiers in Microbiology* 3:324.

339 Barbosa BC, Maciel TT, Abegg AD, Borges LM, Rosa CD, Vargas-Peixoto D. 2016. Arachnids infected by arthropod-pathogenic fungi in an urban fragment of Atlantic Forest in southern Brazil. *Natureza Online* 14:11–14.

340 Ávila Guerrero C. 2019. Caracterización del microhábitat y distribución espacial de *Pamphobeteus ferox* Araneae. Theraphosidae en parches de bosque andino de San Antonio del Tequendama. Thesis por el titulo de Biologo, Universidad de La Salle, Bogota, Colombia.

341 Ortiz D, Bertani R. 2005. A new species in the spider genus *Phormictopus* (Theraphosidae: Theraphosinae) from Cuba. *Revista Ibérica de Aracnología* 11:29–36.

342 Barbosa BC, Maciel TT, Abegg AD, Borges LM, Rosa CD, Vargas-Peixoto D. 2016. Arachnids infected by arthropod-pathogenic fungi in an urban fragment of Atlantic Forest in southern Brazil. *Natureza Online* 14:11–14.

343 Daniel Winkler, pers. comm.

344 Mains EB. 1954. Species of *Cordyceps* on spiders. *Bulletin of the Torrey Botanical Club* 81:492–500.

345 Castillo L, Sanjuan T, Restrepo S, Realpe E. 2015. Efecto del hongo aracnopatógeno *Cordyceps nidus* sp. nov. en tarántulas de la familia Theraphosidae en condiciones de laboratorio. Online at:

346 James Christensen (Minden Pictures) MISSING !!!!

347 Greenstone MH, Ignoffo CM, Samson RA. 1987. Susceptibility of spider species to the fungus *Nomuraea atypicola*. *Journal of Arachnology* 15:266–268.

348 Chandrashekhar et al. 1981 Chandrashekhar S., Suryanarayanan TS, Narasimham CL. 1981. Occurrence of *Beauveria alba* on a spider. *Current Science* 50:248.

349 Martynenko SV, Kondratyukn TO, Sukhomlin MM. 2012. A hyphomycete, *Engyodontium album* (Limber) de Hoog, attacking spiders in underground headings of Kyiv-City. *Ukrainian Botanical Journal* 69:423–432.

350 Costa PP. 2014. *Gibellula* spp. associadas a aranhas da Mata do Paraíso, Viçosa-MG. MSc Thesis, Universidade Federal de Viçosa, Brazil.

351 Gonzaga MO, Leiner NO, Santos AJ. 2006. On the sticky cobwebs of two theridiid spiders (Araneae: Theridiidae). *Journal of Natural History* 40:293–306.

352–353 Costa PP. 2014. *Gibellula* spp. associadas a aranhas da Mata do Paraíso, Viçosa-MG. MSc Thesis, Universidade Federal de Viçosa, Brazil.

354–355 Bibbs CS, Vitoreli AM, Benny G, Harmon CL, Baldwin RW. 2013. Susceptibility of *Latrodectus geometricus* (Araneae: Theridiidae) to a *Mucor* strain discovered in north central Florida, USA. *Florida Entomologist* 96:1052–1061.

356 Mongkolsamrit S, Noisripoom W, Tasanathai K, Kobmoo N, Thanakitpipattana D, Khonsanit A. et al. 2022. Comprehensive treatise of *Hevansia* and three new genera *Jenniferia*, *Parahevansia* and *Polystromomyces* on spiders in Cordycipitaceae from Thailand. *MycoKeys* 91:113–149.

357 Mercado Sierra A, Alayo Soto R, Mena Portales J, de Armas LF. 1988. Hongos entomógenos de Cuba. Nueva especie de *Clathroconium* sobre arañas. *Acta Botánica Cubana* 56:1–5.

358 Costa PP. 2014. *Gibellula* spp. associadas a aranhas da Mata do Paraíso, Viçosa-MG. MSc Thesis, Universidade Federal de Viçosa, Brazil.

359–360 Kuephadungphan W, Petcharad B, Tasanathai K, Thanakitpipattana D, Kobmoo N, Khonsanit A. et al. 2022. Multi-locus phylogeny unmasks hidden species within the specialised spider-parasitic fungus, *Gibellula* (Hypocreales, Cordycipitaceae) in Thailand. *Studies in Mycology* 101:245–286.

361 Mongkolsamrit S, Noisripoom W, Tasanathai K, Kobmoo N, Thanakitpipattana D, Khonsanit A. et al. 2022. Comprehensive treatise of *Hevansia* and three new genera *Jenniferia*, *Parahevansia* and *Polystromomyces* on spiders in Cordycipitaceae from Thailand. *MycoKeys* 91:113–149.

362 Kuephadungphan W, Macabeo APG, Luangsa-Ard JJ, Tasanathai K, Thanakitpipattana D, Phongpaichit S. et al. 2019. Studies on the biologically active second-
ary metabolites of the new spider parasitic fungus *Gibellula gamsii*. Mycological Progress 18:135–146.

363 Kuephadungphan W, Tasanathai K, Petcharad B, Khonsanit A, Stadler M, Luangsa-ard JJ. 2020. Phylogeny- and morphology-based recognition of new species in the spider-parasitic genus *Gibellula* (Hypocreales, Cordycipitaceae) from Thailand. *MycoKeys* 72:17–42.

364–365 Mongkolsamrit S, Noisripoom W, Tasanathai K, Kobmoo N, Thanakitpipattana D, Khonsanit A. et al. 2022. Comprehensive treatise of *Hevansia* and three new genera *Jenniferia*, *Parahevansia* and *Polystromomyces* on spiders in Cordycipitaceae from Thailand. *MycoKeys* 91:113–149.

366 Kuephadungphan W, Petcharad B, Tasanathai K, Thanakitpipattana D, Kobmoo N, Khonsanit A. et al. 2022. Multi-locus phylogeny unmasks hidden species within the specialised spider-parasitic fungus, *Gibellula* (Hypocreales, Cordycipitaceae) in Thailand. *Studies in Mycology* 101:245–286.

367 Greenstone MH, Ignoffo CM, Samson RA. 1987. Susceptibility of spider species to the fungus *Nomuraea atypicola*. *Journal of Arachnology* 15:266–268.

368 Tony DeSantis, pers. comm. (Fig. 4E, this paper).

369 Costa PP. 2014. *Gibellula* spp. associadas a aranhas da Mata do Paraíso, Viçosa-MG. MSc Thesis, Universidade Federal de Viçosa, Brazil.

370 Greenstone MH, Ignoffo CM, Samson RA. 1987. Susceptibility of spider species to the fungus *Nomuraea atypicola*. *Journal of Arachnology* 15:266–268.

371 https://www.flickr.com/photos/128810613@N02/50070178022

372 Kuephadungphan W, Petcharad B, Tasanathai K, Thanakitpipattana D, Kobmoo N, Khonsanit A. et al. 2022. Multi-locus phylogeny unmasks hidden species within the specialised spider-parasitic fungus, *Gibellula* (Hypocreales, Cordycipitaceae) in Thailand. *Studies in Mycology* 101:245–286.

373 Ruszkiewicz-Michalska M, Tkaczuk C, Dynowska M, Sucharzewska E, Szkodzik J, Wrzosek M. 2012. Preliminary studies of fungi in the Biebrza National Park (NE Poland). I. Micromycetes. *Acta Mycologica* 47:213–234.

374 Ruszkiewicz-Michalska M, Balazy S, Chelkowski J, Dynowska M, Pawlowska J, Sucharzewska E et al. 2015. Preliminary studies of fungi in the Biebrza National Park (NE Poland). Part III. Micromycetes-new data. Acta Mycologica 50:1–28. http://dx.doi.org/10.5586/am.1067

375 Serrano Añazco YDL. 2016. Diversidad de hongos entomopatógenos del género Cordyceps sl (Hypocreales: Clavicipitaceae) en el Ecuador. Bachelor's Thesis, Pontificia Universidad Católica del Ecuador, Quito.**

376 Bishop L. 1990a. Entomophagous fungi as mortality agents of ballooning spiderlings. *Journal of Arachnology* 18:237–238

377 https://www.alamy.com/fungus-torrubiella-sp-on-dead-spider-aranaea-order-on-crane-flower-strelitzia-reginae-klungkung-bali-indonesia-image432872997.html

378 Petch T. 1944. Notes on entomogenous fungi. *Transactions of the British Mycological Society* 27:81–93.

379 Kobayasi Y, Shimizu D. 1982. Monograph of the genus *Torrubiella. Bulletin of National Science Museum Tokyo, Series B* 8:43–78.

380 Costa 2014 Costa PP. 2014. *Gibellula* spp. associadas a aranhas da Mata do Paraíso, Viçosa-MG. MSc Thesis, Universidade Federal de Viçosa, Brazil.

381 Saltamachia SJ. 2022. Theoretical and empirical evidence for extended phenotypes in a specialized parasite of spiders. *Authorea Preprint Repository* DOI: 10.22541/au.164604858.89088094/v1.

382 https://www.whatsthatbug.com/2008/10/26/fungus-riddled-spider/

383 https://www.flickr.com/photos/myriorama/14996980578

384 https://bugguide.net/node/view/500724

385 https://bugguide.net/node/view/1723599

386 https://bugguide.net/node/view/1001126

387 Nentwig W, Prillinger H. 1990. A zygomycetous fungus as a mortality factor in a laboratory stock of spiders. *Journal of Arachnology* 18:118–121.

388 https://www.inaturalist.org/observations/89466583

389 https://www.inaturalist.org/observations/93148799

390 Kuephadungphan W, Petcharad B, Tasanathai K, Thanakitpipattana D, Kobmoo N, Khonsanit A. et al. 2022. Multi-locus phylogeny unmasks hidden species within the specialised spider-parasitic fungus, *Gibellula* (Hypocreales, Cordycipitaceae) in Thailand. Studies in Mycology 101:245–286.

391 Daniel Winkler, pers. comm. (Fig. 5F, this paper).

392 Wunderlich J. 2004. Fossil spiders in amber and copal: Conclusions, revisions, new taxa, family diagnoses of fossil and extant taxa. *Beiträge zur Araneologie* 3:1–1908.

393 Costa PP. 2014. *Gibellula* spp. associadas a aranhas da Mata do Paraíso, Viçosa-MG. MSc Thesis, Universidade Federal de Viçosa, Brazil.

394 Kuephadungphan W, Tasanathai K, Petcharad B, Khonsanit A, Stadler M, Luangsa-ard JJ. 2020. Phylogeny- and morphology-based recognition of new species in the spider-parasitic genus *Gibellula* (Hypocreales, Cordycipitaceae) from Thailand. *MycoKeys* 72:17–42.

395–401 Shrestha B, Kubátová A, Tanaka E, Oh J, Yoon DH, Sung JM et al. 2019. Spider-pathogenic fungi within Hypocreales (Ascomycota): their current nomenclature, diversity, and distribution. *Mycological Progress* 18:983–1003.

402 Chen WH, Han YF, Liang ZQ, Jin DC. 2017. A new araneogenous fungus in the genus *Beauveria* from Guizhou, China. *Phytotaxa* 302:57–64.

403 Wang Y, Tang DX, Luo R, Wang YB, Thanarut C, Dao VM, et al. 2023a. Phylogeny and systematics of the genus *Clonostachys*. *Frontiers in Microbiology* 14:1117753.

404 Shrestha B, Kubátová A, Tanaka E, Oh J, Yoon DH, Sung JM et al. 2019. Spider-pathogenic fungi within Hypocreales (Ascomycota): their current nomenclature, diversity, and distribution. *Mycological Progress* 18:983–1003.

405 Mongkolsamrit S, Noisripoom W, Tasanathai K, Khonsanit A, Thanakitpipattana D, Himaman W et al. 2020. Molecular phylogeny and morphology reveal cryptic species in *Blackwellomyces* and *Cordyceps* (Cordycipitaceae) from Thailand. Mycological Progress 19:957–983.

406 Shrestha B, Kubátová A, Tanaka E, Oh J, Yoon DH, Sung JM et al. 2019. Spider-pathogenic fungi within Hypocreales (Ascomycota): their current nomenclature, diversity, and distribution. *Mycological Progress* 18:983–1003.

407 Mongkolsamrit S, Noisripoom W, Luangsa-Ard JJ, Himaman W. 2019. *Cordyceps kuiburiensis*. Persoonia 43:358–359.**

408–420 Shrestha B, Kubátová A, Tanaka E, Oh J, Yoon DH, Sung JM et al. 2019. Spider-pathogenic fungi within Hypocreales (Ascomycota): their current nomenclature, diversity, and distribution. *Mycological Progress* 18:983–1003.

421–422 Chen M, Wang T, Lin Y, Huang B. 2022. Morphological and molecular analyses reveal two new species of *Gibellula* (Cordycipitaceae, Hypocreales) from China. MycoKeys 90:53–69.

423–436 Shrestha B, Kubátová A, Tanaka E, Oh J, Yoon DH, Sung JM et al. 2019. Spider-pathogenic fungi within Hypocreales (Ascomycota): their current nomenclature, diversity, and distribution. *Mycological Progress* 18:983–1003.

437 Evans HC. 2013. Fungal pathogens of spiders. Pp. 107–121. In Spider Ecophysiology. (Nentwig W, ed.). Springer, Berlin, Heidelberg.

438 Evans HC, Samson RA. 1982. Entomogenous fungi from the Galápagos Islands. *Canadian Journal of Botany* 60:2325–2333.

439 Thúy NT, Tùng NV, Lân TN, Lam TTN. 2015. Some biological characteristics of *Isaria javanica* (Frider. & Bally) Samsom & Hywel-Jones distributing at Pu Mat National Park, Nghe An. Journal of Science & Developpement 1:687–693 [in Vietnamese]

440–443 Shrestha B, Kubátová A, Tanaka E, Oh J, Yoon DH, Sung JM et al. 2019. Spider-pathogenic fungi within Hypocreales (Ascomycota): their current nomenclature, diversity, and distribution. *Mycological Progress* 18:983–1003.

444–445 Zhou et al. 2022 Zhou YM, Zhi JR, Qu JJ, Zou X. 2022. Estimated divergence times of *Lecanicillium* in the Family Cordycipitaceae provide insights into the attribution of *Lecanicillium*. *Frontiers in Microbiology*:1379.

446 Chen WH, Liang JD, Ren XX, Zhao JH, Han YF, Liang ZQ. 2022. Phylogenetic, ecological and morphological characteristics reveal two new spider-associated genera in Clavicipitaceae. MycoKeys 91:49–66.

447  Mongkolsamrit S, Noisripoom W, Tasanathai K, Kobmoo N, Thanakitpipattana D, Khonsanit A. et al. 2022. Comprehensive treatise of *Hevansia* and three new genera *Jenniferia*, *Parahevansia* and *Polystromomyces* on spiders in Cordycipitaceae from Thailand. *MycoKeys* 91:113–149.

448  Samson RA, Evans HC. 1973. Notes on entomogenous fungi from Ghana: I. The genera *Gibellula* and *Pseudogibellula*. *Acta Botanica Neerlandica* 22:522–528.

449  Chen WH, Liang JD, Ren XX, Zhao JH, Han YF, Liang ZQ. 2022. Phylogenetic, ecological and morphological characteristics reveal two new spider-associated genera in Clavicipitaceae. MycoKeys 91:49–66.

450  Shrestha B, Kubátová A, Tanaka E, Oh J, Yoon DH, Sung JM et al. 2019. Spider-pathogenic fungi within Hypocreales (Ascomycota): their current nomenclature, diversity, and distribution. *Mycological Progress* 18:983–1003.

451  Johnson D, Sung GH, Hywel-Jones NL, Luangsa-Ard JJ, Bischoff JF et al. 2009. Systematics and evolution of the genus *Torrubiella* (Hypocreales, Ascomycota). Mycological Research 113:279–289.**

452–482  Shrestha B, Kubátová A, Tanaka E, Oh J, Yoon DH, Sung JM et al. 2019. Spider-pathogenic fungi within Hypocreales (Ascomycota): their current nomenclature, diversity, and distribution. *Mycological Progress* 18:983–1003.

483  Humber RA, Hansen KS, Wheeler MM. 2014. USDA-ARS Collection of Entomopathogenic Fungal Cultures – Indexes to available isolates. Robert W. Holley Center for Agriculture and Health, Ithaca, New York.
https://www.ars.usda.gov/ARSUserFiles/80620520/ALL%20AVAIL%20indices%2016Jan014.pdf
Accessed 8 March 2023

484  Evans & Samson 1982 Evans HC, Samson RA. 1982. Entomogenous fungi from the Galápagos Islands. *Canadian Journal of Botany* 60:2325–2333.

485–494  Shrestha B, Kubátová A, Tanaka E, Oh J, Yoon DH, Sung JM et al. 2019. Spider-pathogenic fungi within Hypocreales (Ascomycota): their current nomenclature, diversity, and distribution. *Mycological Progress* 18:983–1003.

495  Montalva C, Silva JJ, Rocha LFN, Luz C, Humber RA. 2019. Characterization of *Tolypocladium cylindrosporum* (Hypocreales, Ophiocordycipitaceae) isolates from Brazil and their efficacy against *Aedes aegypti* (Diptera, Culicidae). *Journal of Applied Microbiology* 126:266–276.**

496  Samson RA, Evans HC. 1982. *Clathroconium*, a new helicosporous hyphomycete genus from spiders. *Canadian Journal of Botany* 60:1577–1580.

497  Malloch D, Kane J, Lahaie DG. 1978. Filobasidiella arachnophila sp. nov. *Canadian Journal of Botany* 56:1823–1826.

498–499  Greif MD, Currah RS. 2007. Patterns in the occurrence of saprophytic fungi carried by arthropods caught in traps baited with rotted wood and dung. *Mycologia* 99:7–19.

500  Tan YP, Bishop-Hurley SL, Shivas RG, Cowan DA, Maggs-Kölling G, Maharachchikumbura SSN et al. 2022. Fungal Planet description sheets: 1436–1477. *Persoonia* 49:261–350.

501 Greif MD, Currah RS. 2007. Patterns in the occurrence of saprophytic fungi carried by arthropods caught in traps baited with rotted wood and dung. *Mycologia* 99:7–19.

502 Wang Y, Liu Y, Zhang G, Zhang M, Zhu K, Wang Y. et al. 2020. Complete mitochondrial genome of *Cladosporium zixishanense* sp. nov. YFCC 8620 isolated from the spider in Yunnan, southwestern China. Mitochondrial DNA Part B 5:210–211.**

503 Aini AN, Mongkolsamrit S, Wijanarka W, Thanakitpipattana D, Luangsa-Ard JJ, Budiharjo A. 2020. Diversity of *Akanthomyces* on moths (Lepidoptera) in Thailand. MycoKeys 71, 1.**

504 Nováková A, Kubátová A, Sklenář F, Hubka V. 2018a. Microscopic fungi on cadavers and skeletons from cave and mine environments. *Czech Mycology* 70:101–121.

505 Wang Y, Tang DX, Luo R, Wang YB, Thanarut C, Dao VM et al. 2023. Phylogeny and systematics of the genus *Clonostachys*. *Frontiers in Microbiology* 14:1117753.

506-508 Chen WH, Liang JD, Ren XX, Zhao JH, Han YF. 2023. Study on species diversity of *Akanthomyces* (Cordycipitaceae, Hypocreales) in the Jinyun Mountains, Chongqing, China. *MycoKeys* 98:299.

509-510 Wang Y, Wang ZQ, Luo R, Souvanhnachit S, Thanarut C, Dao VM, et al. 2023. Species diversity and major host–substrate associations of the genus *Akanthomyces*. *Research Square* https://doi.org/10.21203/rs.3.rs-2907259/v1.

511 Mongkolsamrit S, Sandargo B, Ebada SS, Noisripoom W, Jaiyen S, Luangsa-ard JJ, et al. 2023. *Bhushaniella* gen. nov. (Cordycipitaceae) on spider eggs sac: a new genus from Thailand and its bioactive secondary metabolites. *Mycological Progress* 22:1–16.

6 September 2023
